# TCR signaling via NFATc1 constrains IL-15-induced NK-like activation of human memory CD8^+^ T cells

**DOI:** 10.1101/2025.01.13.632884

**Authors:** Hoyoung Lee, So-Young Kim, Sang-Hoon Kim, Seongju Jeong, Kyung Hwan Kim, Chang Gon Kim, June-Young Koh, Hyung-Don Kim, Ji Won Han, Hosun Yu, Sunwoo Min, Su-Hyung Park, Hyuk Soo Eun, Eui-Cheol Shin

## Abstract

Here we investigated the regulatory mechanisms of TCR-independent bystander activation and NK-like cytotoxicity of human memory CD8^+^ T cells. We found that TCR signals suppressed characteristic features of IL-15-induced CD8^+^ T-cell activation, including increased NKG2D expression and upregulation of genes related to NK cytotoxicity and IFN response. Moreover, ionomycin suppressed IL-15-induced bystander activation and NK-like cytotoxicity, indicating that Ca^2+^-calcineurin signaling is responsible for TCR-mediated suppression of IL-15-induced bystander activation. In detail, NFATc1 suppressed IL-15-induced bystander activation via binding to AP-1 that is necessary for the IL-15-induced upregulation of NK cytotoxicity-related genes. Consistent with these results, calcineurin inhibitors enhanced IL-15-induced NKG2D expression in the presence of TCR signals. Additionally, we defined genes upregulated by IL-15 and downregulated by concurrent TCR signals as an IL-15-induced bystander activation gene set, and found that this gene signature was upregulated in bystander CD8^+^ T cells from patients with hepatitis A virus infection. This study paves the way for further investigation of bystander CD8^+^ T-cell activation in various pathological conditions, and its regulation.

**Graphical abstract:** 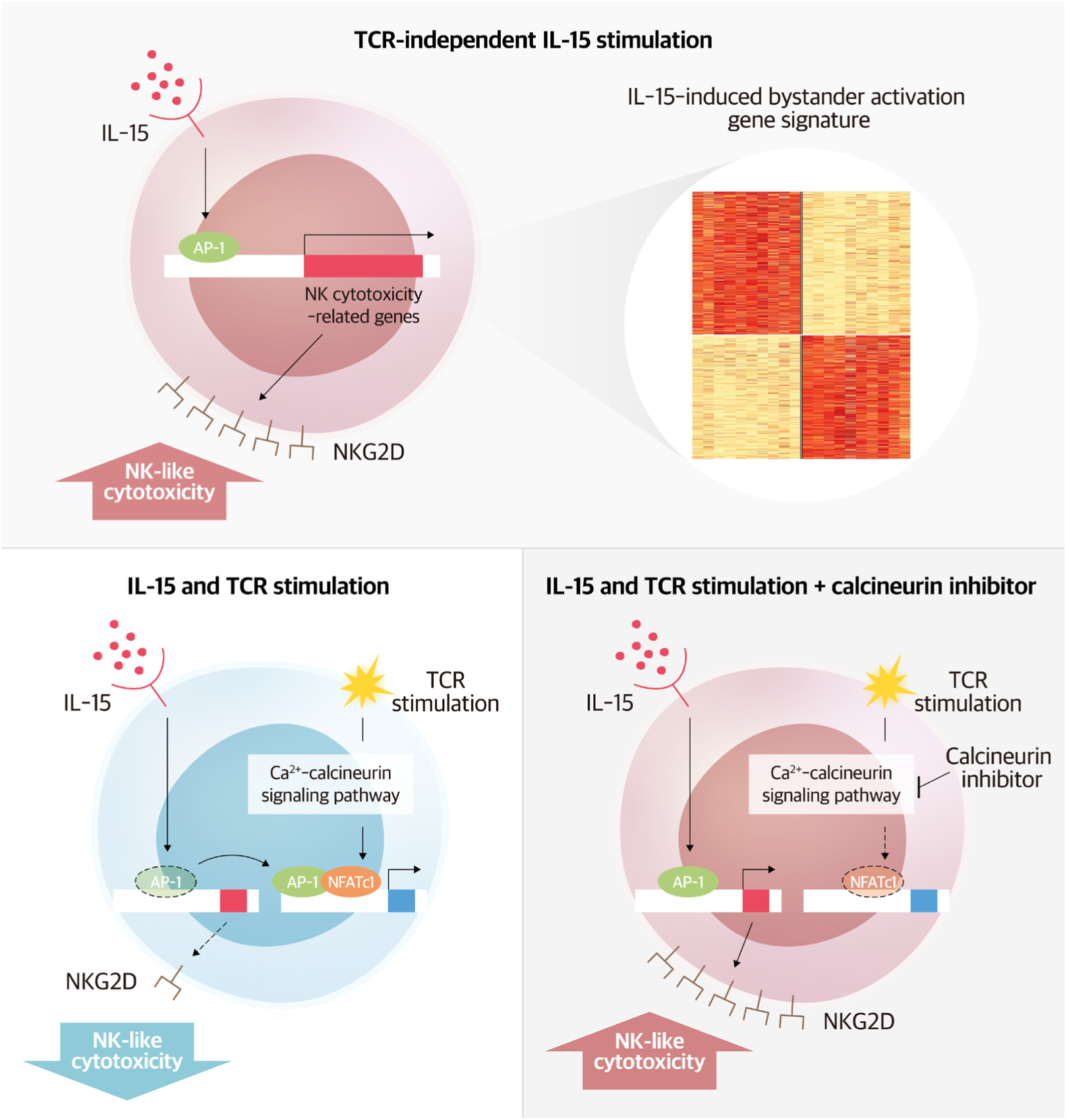

**Highlights:**

- TCR signaling downregulates IL-15-induced transcriptomic features related to NK-like bystander activation of human memory CD8^+^ T cells via the Ca^2+^-calcineurin pathway.
- NFATc1 suppresses IL-15-induced bystander activation via binding to AP-1, which is responsible for IL-15-induced NK-like cytotoxic activity.
- Calcineurin inhibitors cannot suppress IL-15-induced bystander CD8^+^ T-cell activation, and paradoxically increase IL-15-induced NKG2D expression in the presence of TCR signals.
- Genes upregulated by IL-15 and downregulated by concurrent TCR signals define the gene set of bystander CD8^+^ T-cell activation, which was validated in disease contexts.

## Introduction

Memory CD8^+^ T cells provide prompt and enhanced protection against re-exposures to previously encountered pathogens. These cells are primarily activated by T-cell receptor (TCR) signals, which are triggered by binding to epitope peptides presented by major histocompatibility complex class I (MHC-I).^1^ However, memory CD8^+^ T cells can also be activated without TCR stimulation by cytokines, a process termed bystander activation.^2,3^ Interleukin 15 (IL-15) is reportedly the key driver triggering bystander activation of memory _CD8_^+^ T cells.^4,5,6,7,8,9,10^

Bystander activation of memory CD8^+^ T cells can play either protective^6,9,11,12,13,14^ or detrimental^8,15,16,17,18^ roles in microbial infections, depending on the infecting pathogens. For example, in acute hepatitis A virus (HAV) infection, upregulated IL-15 causes bystander activation of pre-existing memory CD8^+^ T cells that are specific for HAV-unrelated viruses.^8^ These bystander-activated memory CD8^+^ T cells are characterized by increased expression of the NK-activating receptor NKG2D, together with cytotoxic effector molecules, such as perforin and granzyme B. Consequently, the bystander-activated memory CD8^+^ T cells recognize and kill target cells via NKG2D-dependent NK-like cytotoxicity, leading to non-specific killing of hepatocytes during acute HAV infection. Notably, this IL-15-induced NK-like cytotoxicity is significantly correlated with liver damage severity in patients with acute HAV infection, indicating the immunopathological role of IL-15-driven bystander-activated memory CD8^+^ T cells.^8^

IL-15-induced bystander-activated memory CD8^+^ T cells are characterized by upregulated NKG2D and CCR5, which is counteracted by the concurrent TCR stimulation.^8,19^ Thus, we can distinguish *bona fide* TCR-independent bystander activation from TCR-mediated conventional activation by examining the expression levels of NKG2D^8^ and CCR5^19^ on CD8^+^ T cells. Additionally, studies of a mouse model of bystander activation revealed that IL-15-induced upregulation of IFN-induced transmembrane protein 3 (IFITM3) is also counteracted by concurrent TCR stimulation.^20^ These findings indicate that antigen-specific TCR signals restrain cytokine-driven nonspecific activation of memory CD8^+^ T cells. However, the regulatory mechanisms of IL-15-induced bystander activation of memory CD8^+^ T cells have not yet been investigated at the molecular level.

In the present study, we investigated specific transcriptional features and regulatory mechanisms of IL-15-induced bystander CD8^+^ T-cell activation. We identified a unique set of genes that were upregulated by IL-15 and downregulated by concurrent TCR stimulation. This gene signature, which is related to NK-cell-mediated cytotoxicity and IFN response, was significantly enriched in bystander CD8^+^ T cells from patients with acute HAV infection. We demonstrated that TCR-induced Ca^2+^-calcineurin signaling counteracted IL-15-induced bystander activation and NK-like cytotoxic activity. Particularly, NFATc1 suppressed IL-15-induced bystander activation via AP-1 binding. Accordingly, calcineurin inhibitors increased IL-15-induced NKG2D upregulation in the presence of TCR signaling. The current findings reveal a regulatory mechanism of bystander CD8^+^ T-cell activation, and provide a bystander activation-specific gene signature that will be useful for further investigations of bystander CD8^+^ T-cell activation in various human diseases.

## Results

### TCR stimulation counteracts IL-15-induced NKG2D upregulation on memory CD8^+^ T cells

We analyzed the expression of NKG2D, a marker that is upregulated upon bystander activation^8^, on human CD8^+^ T cells following *ex vivo* stimulation of healthy donor peripheral blood mononuclear cells (PBMCs) with common gamma chain cytokines, including IL-2, IL-4, IL-7, IL-9, IL-15, and IL-21. The NKG2D expression on CCR7^−^ memory CD8^+^ T cells was increased by IL-2, but was more significantly upregulated by IL-15 (**Figure 1A**). On the other hand, any of the tested cytokines did not increase NKG2D expression in naïve CD8^+^ T cells (**Figure S1**). This IL-15-mediated increase in NKG2D expression on memory CD8^+^ T cells was dose-dependent, and the effect was saturated at 10 ng/ml (**Figure 1B**).

**Figure 1.**
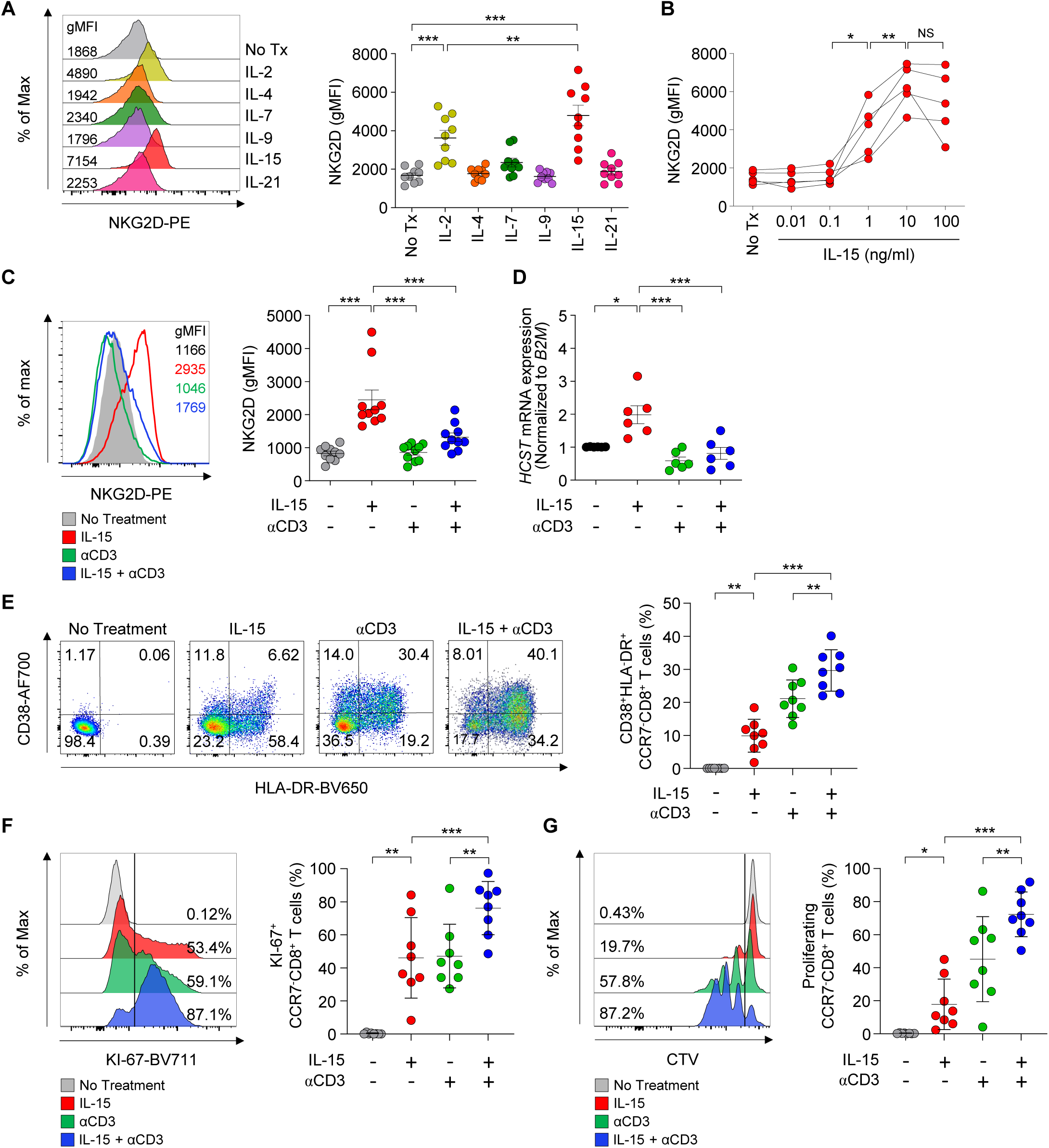
IL-15-induced upregulation of NKG2D on memory CD8^+^ T cells, which is counteracted by concurrent TCR stimulation. **(A and B)** Peripheral blood mononuclear cells (PBMCs) from healthy donors (*n* = 9) were stimulated for 48 hours with IL-2 (10 ng/ml), IL-4 (10 ng/ml), IL-7 (10 ng/ml), IL-9 (10 ng/ml), IL-15 (10 ng/ml) or IL-21 (50 ng/ml). (A) Representative flow cytometry stacked histograms and cumulative data presenting NKG2D expression on CCR7^−^ memory CD8^+^ T cells from healthy donor PBMCs following stimulation with the indicated cytokines. (B) Cumulative data presenting NKG2D expression on CCR7^−^ memory CD8^+^ T cells stimulated for 48 hours with varying doses of IL-15 (0.01, 0.1, 1, 10, and 100 ng/ml). **(C and D)** Sorted CCR7^−^ memory CD8^+^ T cells from healthy donors were stimulated with IL-15 (10 ng/ml), anti-CD3 (coated, 1 µg/ml), or the combination of both for 48 hours. (C) Representative stacked flow cytometry histograms and cumulative data presenting the expression of NKG2D (*n* = 10). (D) mRNA levels of *HCST* (*n* = 6) measured by qPCR with specific primer sets and normalized to *B2M* expression. **(E)** Representative flow cytometry plots and cumulative data presenting the expression CD38 and HLA-DR (*n* = 8) on CCR7^−^ memory CD8^+^ T cells stimulated in the same way as in (C), but for 96 hours. **(F and G)** Sorted CCR7^−^ memory CD8^+^ T cells from healthy donors (*n* = 8) were stained with CellTrace Violet, and stimulated with IL-15 (10 ng/ml), anti-CD3 (coated, 1 µg/ml), or the combination of both for 96 hours. Representative stacked flow cytometry histograms and cumulative data present (F) Ki-67 expression and (G) proliferation of CCR7^−^ memory CD8^+^ T cells. Error bars represent mean ± SD.

Next, we examined how TCR stimulation affected NKG2D expression on memory CD8^+^ T cells. From healthy donor PBMCs, we sorted CCR7^−^ memory CD8^+^ T cells to a purity above 99% (**Figure S2**). Following treatment with IL-15, anti-CD3, or both, the NKG2D expression on memory CD8^+^ T cells was significantly upregulated by IL-15 but not by anti-CD3, and was rather restrained by concurrent anti-CD3 stimulation (**Figure 1C**), confirming our previous findings.^8^ In addition, we found mRNA expression of *HCST*, which encodes DAP10, an adaptor molecule for NKG2D, exhibited the same pattern as surface NKG2D expression on memory CD8^+^ T cells in response to IL-15, anti-CD3, or both (**Figure 1D**). This aligns with previous findings in NK cells, where surface NKG2D expression was shown to correlate with *HCST* (DAP10) mRNA levels following IL-15 stimulaiton.^21^

On the other hand, memory CD8^+^ T-cell activation—as indicated by increased co-expression of CD38 and HLA-DR—was synergistically increased by co-stimulation with IL-15 and anti-CD3, compared to by their individual treatment (**Figure 1E**). Upon co-treatment with IL-15 and anti-CD3, memory CD8^+^ T cells also exhibited synergized increases of Ki-67 expression (**Figure 1F**) and actual proliferation (**Figure 1G**). Taken together, our results demonstrated that TCR and IL-15 synergistically activate memory CD8^+^ T cells while certain aspects of IL-15-induced bystander activation (e.g., NKG2D upregulation) is suppressed by TCR stimulation.

### TCR stimulation suppresses the IL-15-induced upregulation of genes associated with NK-cell-mediated cytotoxicity and IFN response in memory CD8^+^ T cells

To investigate distinct transcriptional signatures induced by IL-15 and TCR stimulation, we performed bulk RNA-seq of sorted memory CD8^+^ T cells following stimulation with IL-15, anti-CD3, or the combination of both. Principal component analysis (PCA) revealed clear segregation of IL-15-stimulated memory CD8^+^ T cells from memory CD8^+^ T cells stimulated with anti-CD3 or the combination of IL-15 and anti-CD3 (**Figure 2A**). This indicated that IL-15-stimulated memory CD8^+^ T cells exhibited unique transcriptional features compared to the others. We identified upregulated differentially expressed genes (DEGs) (**Figure 2B**) and clustered them according to their upregulation patterns (clusters 1–7) (**Figure 2C and Table S1**). Among the 1,401 DEGs upregulated by IL-15 stimulation, 1,037 were additively or synergistically upregulated by IL-15 and anti-CD3 co-stimulation (Cluster 7) indicating additive/synergistic interaction between TCR- and IL-15-induced signals. On the other hand, we found 118 DEGs were upregulated by IL-15 and not further regulated by concurrent anti-CD3 stimulation (Cluster 6), and 214 were upregulated by IL-15 and downregulated by concurrent anti-CD3 stimulation (Cluster 4).

**Figure 2.**
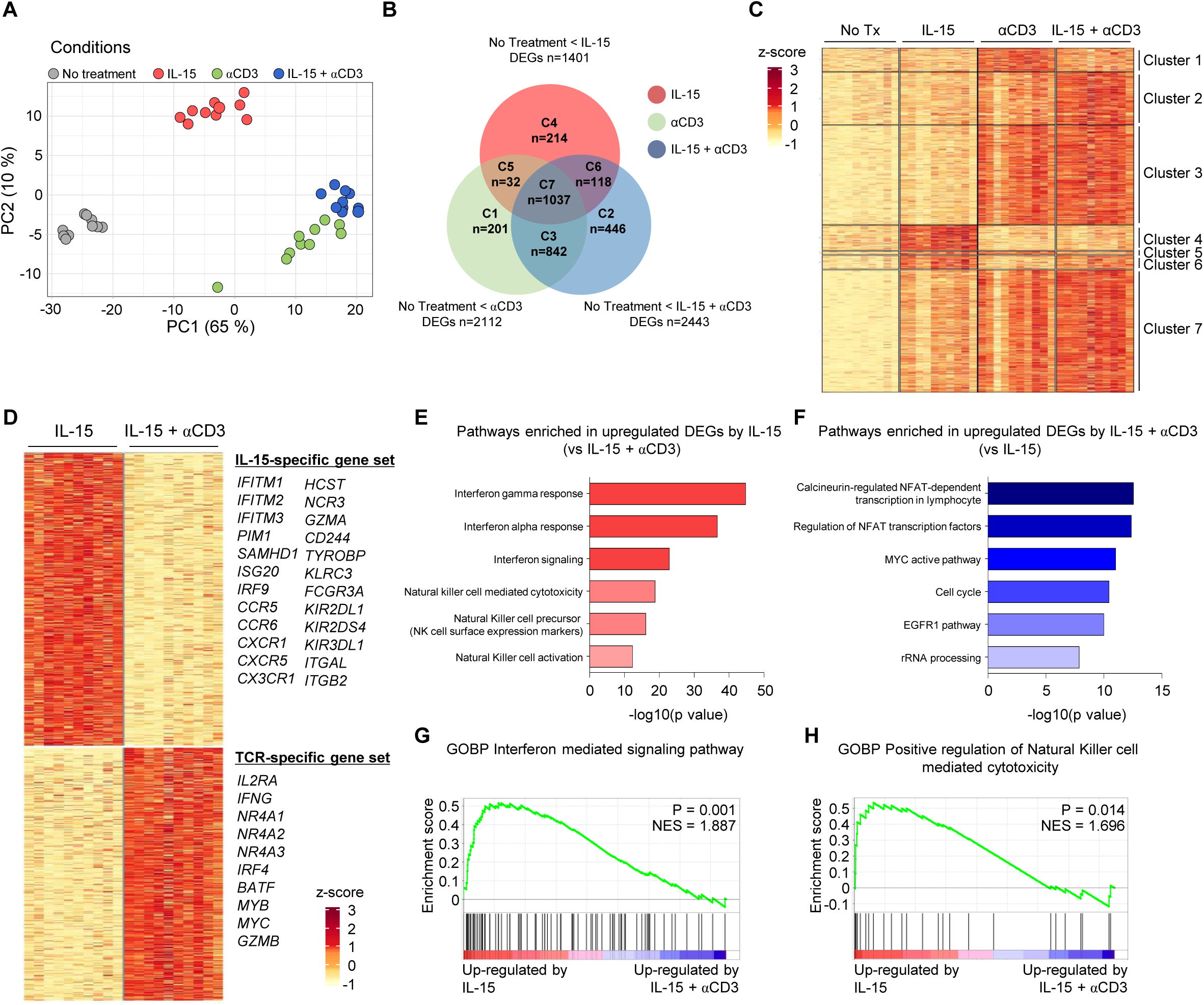
Transcriptional profiles of memory CD8^+^ T cells stimulated with IL-15, anti-CD3, or the combination of both. **(A-H)** Sorted CCR7^−^ memory CD8^+^ T cells from healthy donors (*n* = 10) were stimulated for 48 hours with IL-15 (10 ng/ml), anti-CD3 (coated, 1 µg/ml), or the combination of both. Total RNA was isolated and RNA-seq was performed. **(A)** Principal component analysis (PCA) indicating segregation of the gene expression patterns of memory CD8^+^ T cells stimulated with IL-15, anti-CD3, or the combination of both. **(B)** Venn diagram of individual and overlapping differentially expressed genes of memory CD8^+^ T cells stimulated with IL-15, anti-CD3, or the combination of both, compared to unstimulated cells and **(C)** Heat-map showing the z scores of normalized read counts (adjusted *P* < 0.05, Log2 fold-change > 1). **(D)** Heat-map of differentially expressed genes between memory CD8^+^ T cells stimulated with IL-15 alone or IL-15 plus anti-CD3 (adjusted *P* < 0.05, Log2 fold-change > 1). **(E and F)** Bar-plots showing the enrichment scores (−log10[*p* value]) of various pathways enriched in DEGs upregulated by (E) IL-15 or (F) IL-15 plus anti-CD3. **(G and H)** GSEA of (G) GOBP interferon-mediated signaling pathway (GO: 0140888) and (H) GOBP positive regulation of natural killer-cell-mediated cytotoxicity (GO: 0045954) in memory CD8^+^ T cells stimulated with IL-15 alone versus IL-15 plus anti-CD3.

We further directly compared data between IL-15 stimulation and IL-15 plus anti-CD3 co-stimulation (**Figure 2D and Table S2**). We identified a gene set that was upregulated by IL-15 and downregulated by concurrent anti-CD3 (designated as the IL-15-specific gene set), and a gene set that was upregulated by IL-15 and anti-CD3 co-stimulation, but not by IL-15 alone (designated as the TCR-specific gene set). The IL-15-specific gene set included IFN-responsive genes (*IFITM1*, *IFITM2*, *IFITM3*, *PIM1* and *SAMHD1*) and genes associated with NK-cell-mediated cytotoxicity (*HCST*, *NCR3*, *GZMA, CD244 and TYROBP*). The TCR-specific gene set included genes upregulated by TCR stimulation (*IL2RA*, *IFNG*, *NR4A1*, *NR4A2*, *NR4A3*, *IRF4*, *BATF* and *MYC*). Pathway analysis confirmed that the IL-15-specific gene set was significantly enriched with genes related to IFN response and NK-cell-mediated cytotoxicity (**Figure 2E**), and that the TCR-specific gene set was significantly enriched with genes associated with calcineurin-regulated NFAT-dependent transcription, MYC pathway, and cell cycle (**Figure 2F**). Gene set enrichment analysis also demonstrated that the IL-15-specific gene set was significantly enriched with genes associated with the IFN-mediated signaling pathway (**Figure 2G**) and NK-cell-mediated cytotoxicity (**Figure 2H**). Importantly, the IL-15-specific gene set included genes for DAP10 (*HCST*), CCR5 (*CCR5*), and IFITM3 (*IFITM3*), which have been shown to be upregulated by IL-15, but downregulated by concurrent TCR stimulation in memory CD8^+^ T cells.^19,20^

### Ca^2+^-mediated signaling suppresses characteristic features of IL-15-induced bystander CD8^+^ T-cell activation

We further examined why TCR stimulation suppresses characteristic features of IL-15-induced bystander CD8^+^ T-cell activation, including the upregulation of genes associated with NK-cell-mediated cytotoxicity and IFN response. We hypothesized that the Ca^2+^-calcineurin signaling pathway might be responsible for this unique phenomenon, because this pathway is activated by TCR stimulation,^22^ but not by IL-15 stimulation. We first confirmed that the calcineurin inhibitor FK506 abrogated the TCR-induced activation and proliferation of memory CD8^+^ T cells (**Figure S3A**), but not IL-15-induced activation and proliferation (**Figure S3B**). On the other hand, the mTOR inhibitor everolimus abolished both TCR- and IL-15-induced activation and proliferation. Consistent with these findings, treatment with another type of calcineurin inhibitor, cyclosporin A (CsA), significantly abrogated the TCR-induced activation and proliferation of memory CD8^+^ T cells (**Figure S3C**) but did not observably affect IL-15-induced activation and proliferation (**Figure S3D**).

To investigate how the Ca^2+^-calcineurin signaling pathway influenced IL-15-induced bystander CD8^+^ T-cell activation, we used the Ca^2+^ ionophore ionomycin.^23^ Sorted human memory CD8^+^ T cells were treated with IL-15, ionomycin, or the combination of both, and then subjected to bulk RNA-seq. PCA revealed clear segregation of IL-15-stimulated memory CD8^+^ T cells from the memory CD8^+^ T cells stimulated with ionomycin or a combination of IL-15 plus ionomycin (**Figure 3A**). We identified upregulated DEGs (**Figure 3B**) and clustered them according to their upregulation patterns, from cluster 1 to 7 (**Figure 3C and Table S3**). Like TCR co-stimulation, concurrent ionomycin co-stimulation downregulated a subset of DEGs that were upregulated by IL-15 stimulation (cluster 4). We directly compared IL-15 stimulation and co-stimulation with IL-15 plus ionomycin. This led to identification of a gene set that was upregulated by IL-15 and downregulated by concurrent ionomycin treatment (**Figure 3D and Table S4**), which was significantly enriched with genes related to IFN response and NK-cell-mediated cytotoxicity (**Figure 3E**).

**Figure 3.**
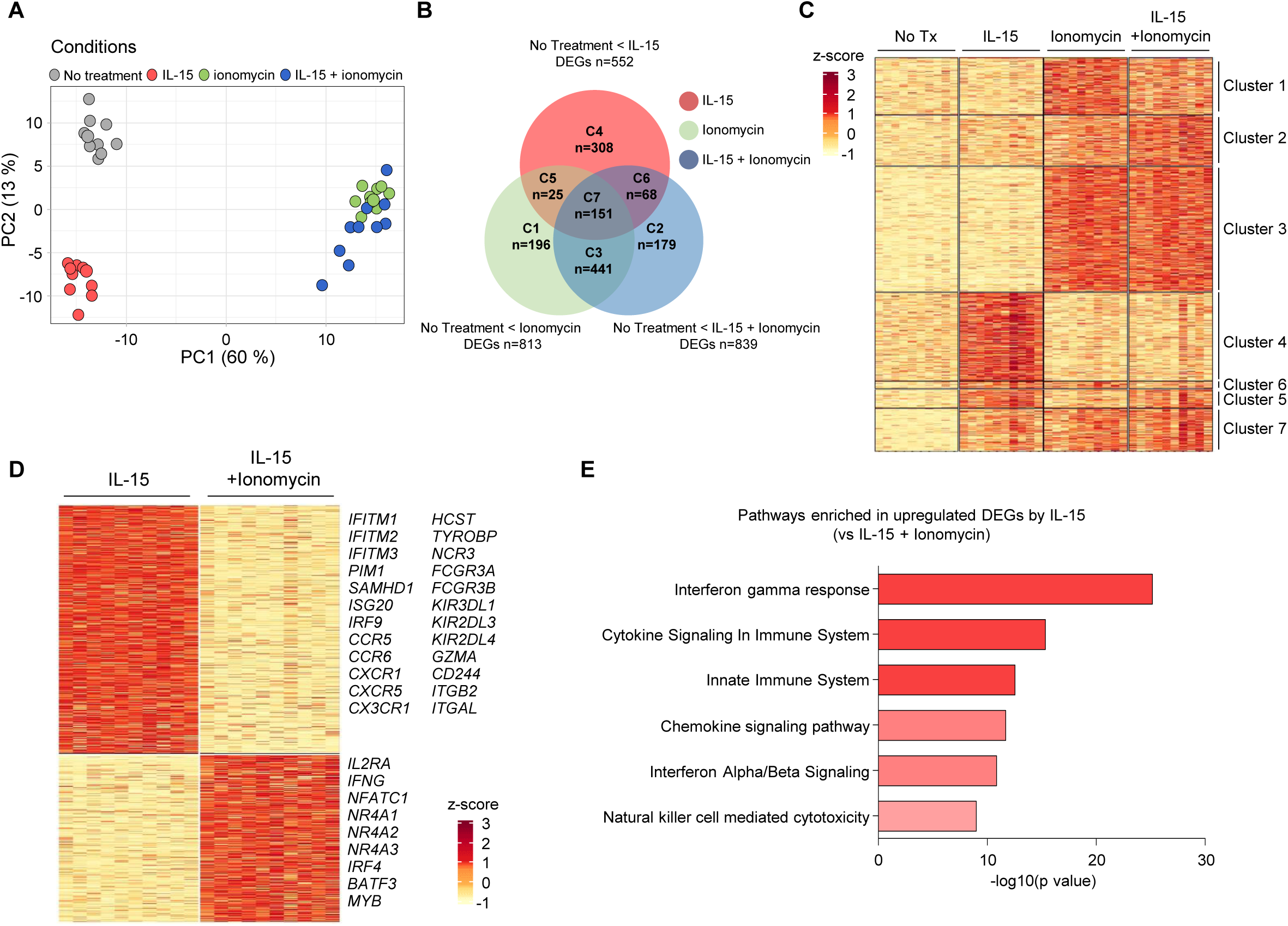
Transcriptional profiles of memory CD8^+^ T cells stimulated with IL-15, ionomycin, or the combination of both. **(A-E)** Sorted CCR7^−^ memory CD8^+^ T cells from healthy donors (*n* = 10) were stimulated for 12 hours with IL-15 (10 ng/ml), ionomycin (500 ng/ml), or the combination of both. Total RNA was isolated and RNA-seq was performed. **(A)** PCA indicating segregation of gene expression patterns of memory CD8^+^ T cells stimulated with IL-15, ionomycin, or the combination of both. **(B)** Venn diagram of individual and overlapping differentially expressed genes of memory CD8^+^ T cells stimulated with IL-15, ionomycin, or the combination of both compared to unstimulated cells and **(C)** Heat-map showing the z scores of normalized read counts (adjusted *P* < 0.05, Log2 fold-change > 1). **(D)** Heat-map of differentially expressed genes between memory CD8^+^ T cells stimulated with IL-15 or with IL-15 plus ionomycin (adjusted *P* < 0.05, Log2 fold-change > 1). **(E)** Bar-plots showing the enrichment scores (−log10[*p* value]) of various pathways enriched in DEGs upregulated by IL-15 and down-regulated by IL-15 plus ionomycin. Statistical analysis was performed using the paired Student’s t test. \*\*\*\**P* < 0.0001.

### Ca^2+^-mediated signaling suppresses IL-15-induced NK-like cytotoxic activity of memory CD8^+^ T cells

We examined the effects of Ca^2+^-mediated signals on the IL-15-induced NK-like cytotoxic activity of memory CD8^+^ T cells, which is mediated by IL-15-induced upregulation of NKG2D.^8^ First, we analyzed NKG2D expression on sorted memory CD8^+^ T cells from healthy donors following *ex vivo* stimulation with IL-15, ionomycin, or the combination of both. In line with how TCR stimulation affected IL-15-induced NKG2D expression, ionomycin inhibited IL-15-induced expression of NKG2D on memory CD8^+^ T cells (**Figure 4A**). However, such ionomycin-induced inhibition did not affect upregulation of the activation markers CD38 and HLA-DR (**Figure 4B**). These findings indicated that Ca^2+^-mediated signals counteracted only certain aspects of IL-15-induced bystander CD8^+^ T-cell activation, including the increased surface expression of NKG2D, and the upregulation of genes associated with NK-cell-mediated cytotoxicity and IFN response. We further examined the effect of calcineurin inhibitors on NKG2D expression. As expected, FK506 did not abrogate IL-15-induced NKG2D upregulation on memory CD8^+^ T cells (**Figure S4**). However, FK506 increased the NKG2D expression when memory CD8^+^ T cells were stimulated with IL-15 in the presence of anti-CD3 (**Figure 4C**). This observation was consistent with the results of CsA treatment (**Figure S5**). This paradoxical effect of calcineurin inhibitors can be explained by inhibition of the counteracting effect of the Ca^2+^-calcineurin pathway on IL-15-induced NKG2D upregulation.

**Figure 4.**
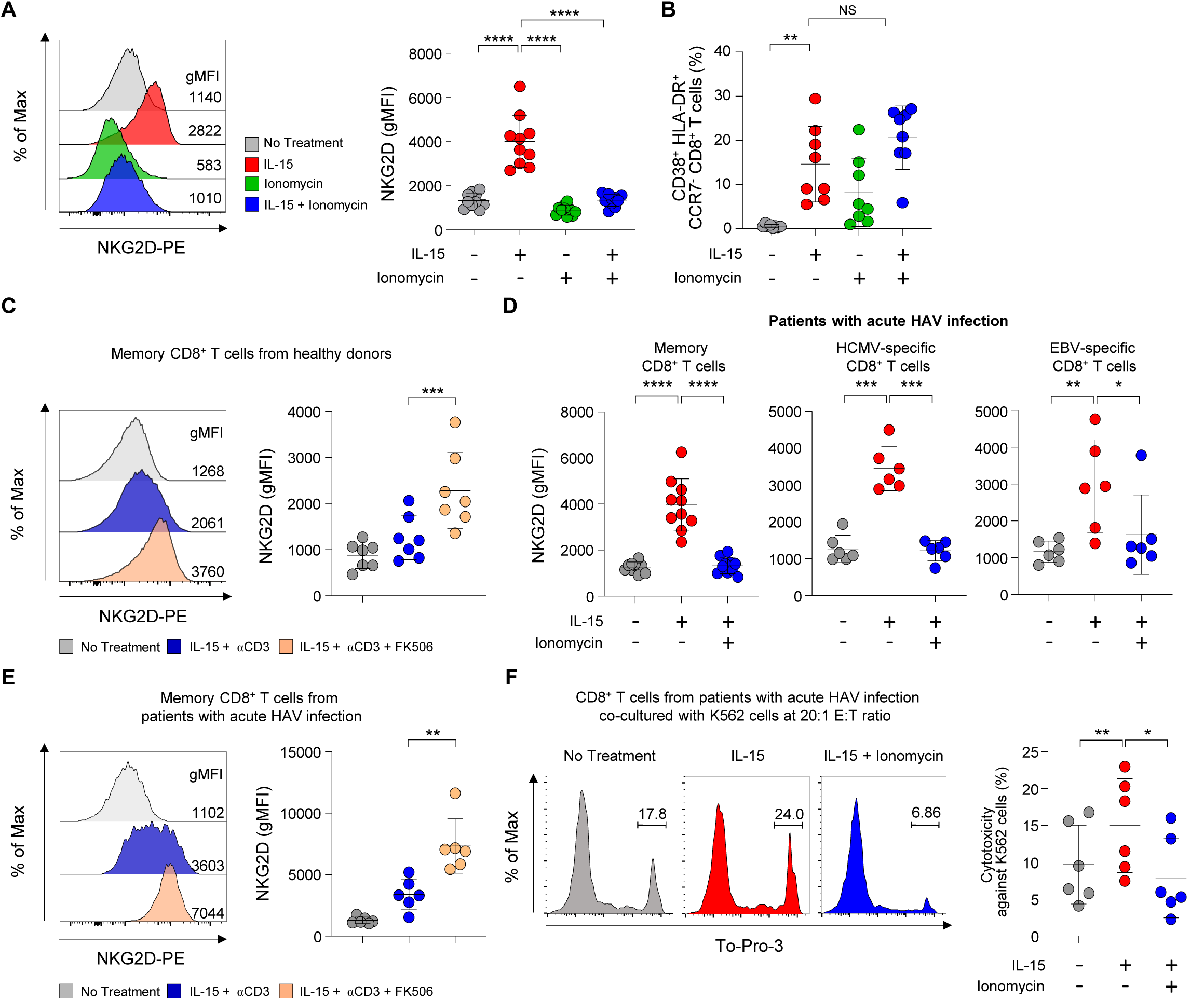
The role of Ca^2+^-calcineurin signaling on IL-15-induced NK-like cytotoxicity of CD8^+^ T cells. **(A and B)** Sorted CCR7^−^ memory CD8^+^ T cells from healthy donors were stimulated with IL-15 (10 ng/ml), ionomycin (500 ng/ml), or the combination of both. (A) Representative stacked flow cytometry histograms and cumulative data presenting the expression of NKG2D (*n* = 10) on CCR7^−^ memory CD8^+^ T cells after 48 hours of stimulation. (B) Cumulative data presenting expression of CD38 and HLA-DR (*n* = 8) on CCR7^−^ memory CD8^+^ T cells after 96 hours of stimulation. **(C)** Sorted CCR7^−^ memory CD8^+^ T cells from healthy donors (*n* = 7) were pretreated with FK506 (10 ng/ml) for 1 hour, followed by stimulation with IL-15 (10 ng/ml) and anti-CD3 (coated, 1 µg/ml) for 48 hours. Representative stacked flow cytometry histograms and cumulative data present NKG2D expression on CCR7^−^ memory CD8^+^ T cells. **(D)** PBMCs from patients with acute HAV infection were stimulated with IL-15 (10 ng/ml) or a combination of IL-15 plus ionomycin (500 ng/ml) for 48 hours. Cumulative data present NKG2D expression on CCR7^−^ memory (*n* = 10), HCMV-specific (*n* = 6), and EBV-specific (*n* = 6) CD8^+^ T cells. **(E)** Sorted CCR7^−^ memory CD8^+^ T cells from patients with acute HAV infection (*n* = 6) were pretreated with FK506 (10 ng/ml) for 1 hour, followed by stimulation with IL-15 (10 ng/ml) and anti-CD3 (coated, 1 µg/ml) for 48 hours. Representative stacked flow cytometry histograms and cumulative data present NKG2D expression on CCR7^−^ memory CD8^+^ T cells. **(F)** Sorted total CD8^+^ T cells from patients with acute HAV infection (*n* = 6) were stimulated with IL-15 (10 ng/ml) or a combination of IL-15 plus ionomycin (500 ng/ml) for 24 hours, and co-cultured with PKH-26-labeled K562 cells at a 20:1 effector:target cell (E:T) ratio for 12 hours. Cytotoxicity was analyzed by flow cytometry following staining of the dead cells with TO-PRO-3 and presented as representative histograms and cumulative data. Error bars represent mean ± SD. Statistical analysis was performed using the paired Student’s t test. \**P* < 0.05, \*\**P* < 0.01, \*\*\**P* < 0.001, \*\*\*\**P* < 0.0001.

We extended our analysis to examine CD8^+^ T cells obtained from patients with acute HAV infection, in which the bystander activation of memory CD8^+^ T cells is well-exemplified.^8^ Analyzing the NKG2D expression after one day of *ex vivo* stimulation with IL-15, with or without ionomycin, revealed that IL-15-induced NKG2D upregulation on memory CD8^+^ T cells was significantly inhibited by the concurrent ionomycin stimulation (**Figure 4D**). We found the same result when we analyzed human cytomegalovirus (HCMV)- and Epstein-Barr virus (EBV)-specific MHC-I multimer^+^CD8^+^ T cells. The paradoxical effect of FK506 on NKG2D expression was also observed when memory CD8^+^ T cells from patients with acute HAV infection were treated with IL-15 in the presence of anti-CD3 (**Figure 4E**). Finally, we investigated the effect of ionomycin on IL-15-induced NK-like cytotoxic activity, which is mediated by NKG2D. We sorted CD8^+^ T cells from PBMCs of patients with acute HAV infection (**Figure S6**), and evaluated the NK-like cytotoxic activity against K562 target cells that lacked MHC-I expression. We found that ionomycin significantly decreased the IL-15-induced NK-like cytotoxic activity of CD8^+^ T cells from patients with acute HAV infection (**Figure 4F**). Collectively, our findings demonstrated that the IL-15-induced NK-like cytotoxic activity of bystander-activated memory CD8^+^ T cells was restrained by the Ca^2+^-calcineurin signaling pathway.

### NFATc1 suppresses IL-15-induced NKG2D upregulation via AP-1 binding

We further investigated a mechanism how the Ca^2+^-calcineurin signaling pathway constrains IL-15-induced NK-like activation of memory CD8^+^ T cells at the transcription factor level. First, we found that SP600125, a JNK inhibitor, abrogated IL-15-induced NKG2D upregulation (**Figure 5A**). In addition, SR11302, an AP-1 inhibitor, also abolished IL-15-induced NKG2D upregulation, indicating that IL-15 upregulates NKG2D expression via AP-1 (**Figure 5B**). Furthermore, we performed siRNA transfection for knocking-down the expression of c-Jun and c-Fos, and IL-15-induced NKG2D upregulation was successfully abrogated by transfection of siRNAs for c-Jun and c-Fos (**Figure 5C**).

**Figure 5.**
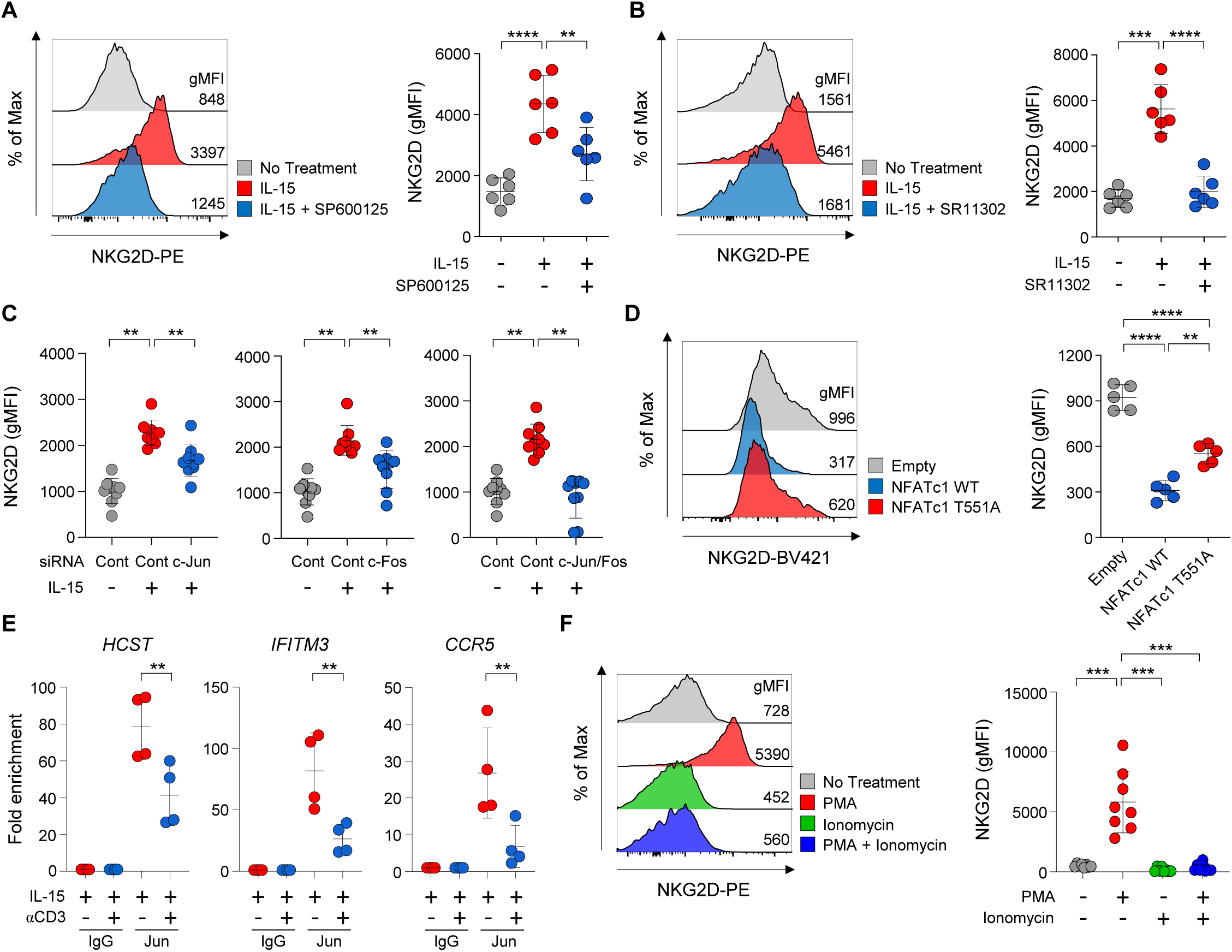
NFATc1-mediated suppression of IL-15-induced NKG2D upregulation through AP-1 binding. **(A and B)** PBMCs from healthy donors (*n* = 6) were pretreated with SP600125 (10 μM) or SR11302 (10 μM) for 1 hour, followed by stimulation with IL-15 (10 ng/ml) for 48 hours. Representative stacked flow cytometry histograms and cumulative data present NKG2D expression on CCR7^−^ memory CD8^+^ T cells pretreated with (A) SP600125 or (B) SR11302. **(C)** PBMCs from healthy donors (*n* = 9) were transfected with c-Jun siRNA (200 nM), c-fos siRNA (200 nM) or the combination of both siRNAs and rested for 4 days. Subsequently, PBMCs were stimulated with IL-15 (10 ng/ml) for 48 hours. Cumulative data present NKG2D expression on CCR7^−^ memory CD8^+^ T cells. **(D)** PBMCs from healthy donors (*n* = 5) were transfected with pcDNA3.1 plasmids containing wild-type or T551A mutant NFATc1. Subsequently, PBMCs were stimulated with IL-15 (10 ng/ml) for 48 hours. Representative stacked flow cytometry histograms and cumulative data present NKG2D expression on CCR7^-^ memory CD8^+^ T cells. **(E)** ChIP-qPCR assays for Jun occupancy at the highly open chromatin genetic regions of *HCST*, *IFITM3* or *CCR5* in sorted CCR7^-^ memory CD8^+^ T cells from healthy donors (*n* = 4) stimulated for 48 hours with IL-15 (10 ng/ml), anti-CD3 (coated, 1 µg/ml), or the combination of both. The y-axis represents fold enrichment, calculated as the fold change of immunoprecipitated DNA with c-Jun relative to immunoprecipitated DNA with IgG, analyzed by ChIP-qPCR using the primers described in the methods section. **(F)** Sorted CCR7^−^ memory CD8^+^ T cells from healthy donors (*n* = 8) were stimulated with PMA (10 ng/ml), ionomycin (500 ng/ml), or the combination of both for 48 hours. Representative stacked flow cytometry histograms and cumulative data present NKG2D expression on CCR7^−^ memory CD8^+^ T cells. Error bars represent mean ± SD. Statistical analysis was performed using the paired Student’s t test for (A) to (D) and (F) and one-way ANOVA test for (E). \*\**P* < 0.01, \*\*\**P* < 0.001, \*\*\*\**P* < 0.0001.

Next, we hypothesized that IL-15-induced AP-1 cannot upregulate NKG2D when it is bound by NFATc1 activated by the Ca^2+^-calcineurin signaling pathway in the presence of concurrent TCR stimulation. To validate this hypothesis, we performed transfection with plasmids coding for wild-type NFATc1 as well as T551A mutant NFATc1 that cannot bind to AP-1.^24^ NKG2D was upregulated by IL-15 in memory CD8^+^ T cells transfected with control plasmids, and wild-type NFATc1 transfection abrogated IL-15-induced NKG2D upregulation (**Figure 5D**). Importantly, the NKG2D expression in T551A NFATc1-transfected cells was significantly higher than that in wild-type NFATc1-transfected cells although it was not fully restored to the level in control plasmid-transfected cells. This result supports our hypothesis that TCR-induced NFATc1 suppresses IL-15-induced NKG2D upregulation via binding to AP-1. We also investigated whether AP-1 directly binds to its predicted binding sites in genes associated with IL-15-induced bystander activation of memory CD8^+^ T cells and whether this binding diminishes with concurrent TCR stimulation. Chromatin immunoprecipitation followed by quantitative PCR (ChIP-qPCR) confirmed that AP-1 binding at *HCST*, *IFITM3*, and *CCR5* under IL-15 stimulation, which was significantly reduced by concurrent TCR stimulation (**Figure 5E**).

Finally, we validated our hypothesis by using PMA that activates AP-1 via protein kinase C activation. In this experiment, we used PMA and ionomycin instead of IL-15 and TCR stimulation. As expected, The NKG2D expression on memory CD8^+^ T cells was significantly upregulated by PMA but not by ionomycin, and was restrained by combined treatment with PMA and ionomycin (**Figure 5F**). Taken together, we conclude that TCR signaling via NFATc1 constrains IL-15-induced NK-like activation of memory CD8^+^ T cells.

### The gene set of IL-15-induced bystander activation is defined and validated

In Figure 2D, the IL-15-specific gene set was identified from genes upregulated by IL-15 and downregulated by concurrent anti-CD3 stimulation, suggesting that the IL-15-specific gene set can be used to distinguish IL-15-induced bystander activation from TCR-induced conventional activation. We investigated whether the IL-15-specific gene set was actually upregulated in TCR-independently IL-15-activated bystander CD8^+^ T cells in acute HAV infection, where the bystander activation of memory CD8^+^ T cells is well-exemplified.^8^

To this end, we performed scRNA-seq using CD8^+^ T cells obtained from patients with acute HAV infection or healthy donors. We sorted CD8^+^ T cells from PBMCs, and stained them with DNA-barcoded MHC-I multimers (dCODE dextramers) to detect CD8^+^ T cells specific for HAV, HCMV, EBV, and influenza A virus (IAV) (**Figure 6A**). We further sorted dCODE dextramer-positive and -negative cells, and then re-mixed them at a 1:9 ratio. Next, the cells were labeled with antibody-derived tags (ADTs), and scRNA-seq was performed using the Chromium system. In uniform manifold approximation and projection (UMAP) visualization, we successfully detected dCODE^+^ HAV-specific CD8^+^ T cells, together with HCMV-, EBV- and IAV-specific CD8^+^ T cells, which we defined as bystander CD8^+^ T cells (**Figure 6B**). Compared to those from healthy donors, the bystander CD8^+^ T cells from patients with acute HAV infection exhibited increased surface protein levels of activation markers including CD38 and HLA-DR, and the genes for cytotoxic molecules *PRF1* and *GZMB* (**Figure 6C**). Moreover, we confirmed the upregulation of NKG2D, CCR5 and *HCST* in bystander CD8^+^ T cells compared to HAV-specific CD8^+^ T cells (**Figure 6D**). This validated our previous finding^8, 19^ that bystander CD8^+^ T cells are activated during acute HAV infection.

**Figure 6.**
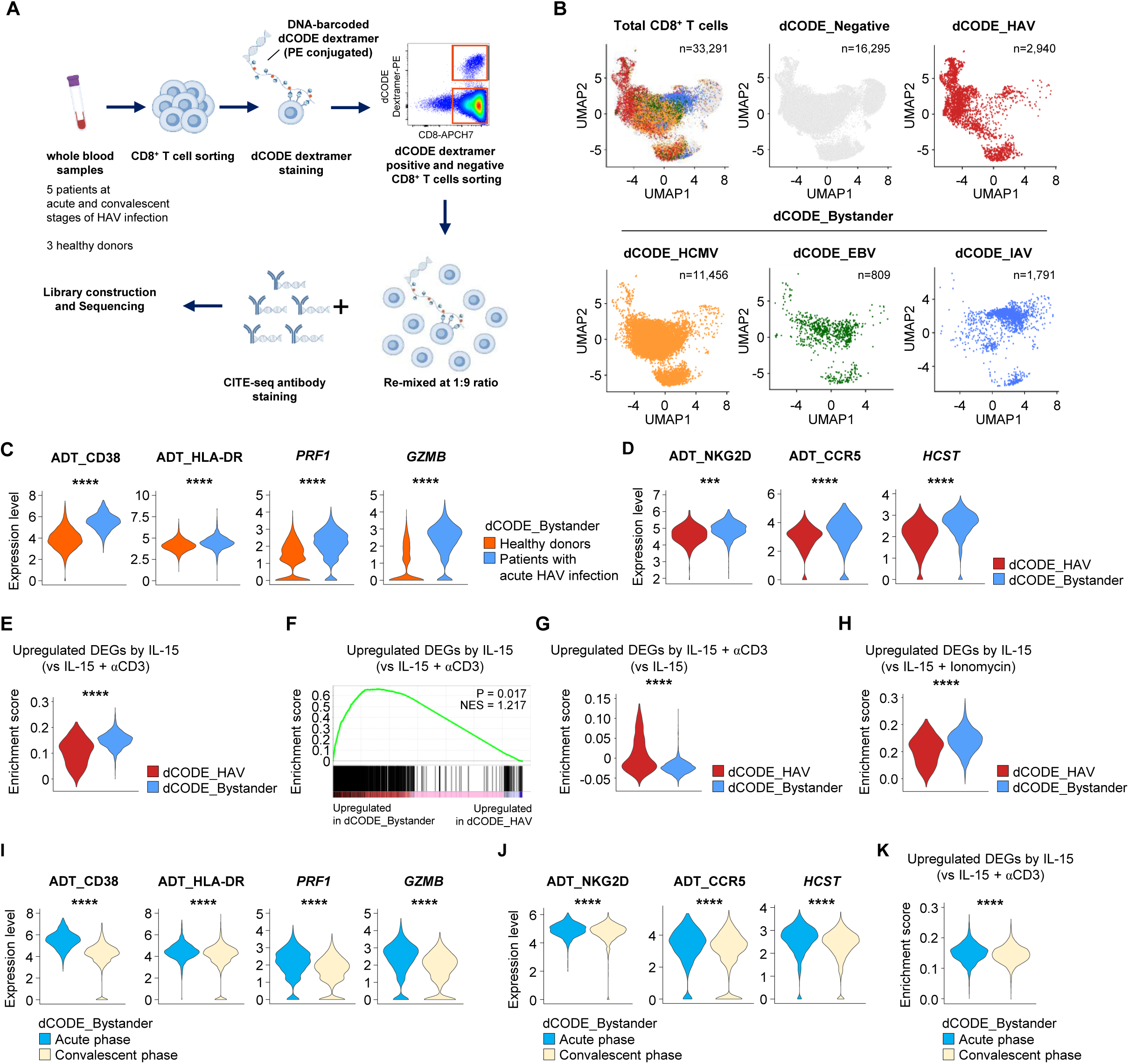
High-throughput single-cell analysis of virus-specific CD8^+^ T cells from patients with HAV infection. **(A-K)** Total CD8^+^ T cells were isolated from patients at acute and convalescent stages of HAV infection (*n* = 5) and healthy donors (*n* = 3) by magnetic separation, and stained with DNA barcode-labeled MHC class I dextramers conjugated with PE (dCODE dextramers). dCODE dextramer-positive and - negative CD8^+^ T cells were flow-sorted, and then re-mixed at a 1:9 ratio and further stained with ADTs. Subsequently, these cells were processed for single-cell RNA sequencing. **(A)** Schematic overview of the experimental design. **(B)** UMAP projections of 33,291 total CD8^+^ T cells, including dCODE-negative (*n* = 16,295) and HAV-specific CD8^+^ T cells (*n* = 2,940), and bystander CD8^+^ T cells, including HCMV-specific (*n* = 11,456), EBV-specific (*n* = 809) and IAV-specific (*n* = 1,791) CD8^+^ T cells. **(C)** Violin plots showing ADT expression of CD38 and HLA-DR, and normalized gene expression of *PRF1* and *GZMB* in bystander (CMV-, EBV- and IAV-specific) CD8^+^ T cells from patients with acute HAV infection versus those from healthy donors. **(D)** ADT expression of NKG2D and CCR5, and normalized gene expression of *HCST* in HAV-specific versus bystander CD8^+^ T cells from patients with acute HAV infection. **(E)** Violin plot showing enrichment score for DEGs upregulated by IL-15 compared to IL-15 plus anti-CD3 in memory CD8^+^ T cells, analyzed in HAV-specific and bystander CD8^+^ T cells from patients with acute HAV infection. **(F)** GSEA of upregulated DEGs from IL-15-stimulated memory CD8^+^ T cells, using the transcriptomes of HAV-specific versus bystander CD8^+^ T cells from patients with acute HAV infection obtained using a pseudo bulk approach. **(G)** Violin plot showing enrichment score for DEGs upregulated by IL-15 plus anti-CD3 compared to IL-15 alone in memory CD8^+^ T cells, analyzed in HAV-specific and bystander CD8^+^ T cells from patients with acute HAV infection. **(H)** Violin plot showing enrichment score for DEGs upregulated by IL-15 compared to IL-15 plus ionomycin in memory CD8^+^ T cells, analyzed in HAV-specific and bystander CD8^+^ T cells from patients with acute HAV infection. **(I and J)** Violin plots showing (I) ADT expression of CD38 and HLA-DR, and normalized gene expression of *PRF1* and *GZMB*, and (J) ADT expression of NKG2D and CCR5, and normalized gene expression of *HCST* in bystander CD8^+^ T cells from patients at the acute and convalescent stages of HAV infection. **(K)** Violin plot showing enrichment score for DEGs upregulated by IL-15 compared to IL-15 plus anti-CD3 in memory CD8^+^ T cells, analyzed in bystander CD8^+^ T cells from patients at the acute and convalescent stages of HAV infection. Statistical analysis was performed using the Mann-Whitney U-test \*\*\**P* < 0.001, \*\*\*\**P* < 0.0001.

Next, we applied the IL-15-specific gene set (which was upregulated by IL-15 and downregulated by concurrent anti-CD3) in bulk RNA-seq to scRNA-seq analysis. The IL-15-specific gene set was significantly enriched in bystander CD8^+^ T cells compared to HAV-specific CD8^+^ T cells from patients with acute HAV infection (**Figures 6E and F**). Additionally, we found that the IL-15-specific gene set was significantly enriched in dCODE^−^ effector/memory CD8^+^ T cell cluster compared to HAV-specific CD8^+^ T cells from patients with acute HAV infection (**Figure S7**). This indicated that the dCODE^−^ effector/memory CD8^+^ T-cell population included a considerable proportion of non-HAV-specific bystander cells. On the other hand, the TCR-specific gene set (a gene set that was upregulated by IL-15 and anti-CD3 co-stimulation, but not by IL-15 alone in Figure 2D) was significantly upregulated in HAV-specific CD8^+^ T cells compared to bystander CD8^+^ T cells (**Figure 6G**). These findings suggest that the IL-15-specific gene set and TCR-specific gene set will be useful for discriminating between IL-15-induced bystander activation and TCR-induced conventional activation. We also analyzed a gene set that was upregulated by IL-15 and downregulated by concurrent ionomycin treatment, and found that it was significantly enriched in bystander CD8^+^ T cells compared to HAV-specific CD8^+^ T cells (**Figure 6H**), indicating that genes upregulated by IL-15 and counteracted by Ca^2+^ are characteristic features of bystander-activated memory CD8^+^ T cells during acute HAV infection. Finally, we found that activation markers (**Figure 6I**), expression of NKG2D, CCR5 and *HCST* (**Figure 6J**), and enrichment of the IL-15-specific gene set (**Figure 6K**) in bystander CD8^+^ T cells were significantly downmodulated during the convalescent phase.

Detailed analysis revealed that IFN response-related genes (*IFI44L*, *IFIT3*, *IFITM1*, *IFITM2*, *IFNAR1*, *IFNAR2*, *IFNGR1, PIM1 and SAMHD1*) were significantly enriched in bystander CD8^+^ T cells compared to HAV-specific CD8^+^ T cells (**Figure S8A**). Furthermore, surface proteins (**Figure S8B**) and genes (**Figure S8C**) associated with NK-cell-mediated cytotoxicity were also significantly increased on bystander CD8^+^ T cells compared to HAV-specific CD8^+^ T cells. On the other hand, compared to bystander CD8^+^ T cells, HAV-specific CD8^+^ T cells exhibited upregulation of CD38, HLA-DR, PD-1, LAG-3, TIM-3, and CD39, together with *IFNG, PRF1, GZMB,* and *MKI67* (**Figure S9**).

We also validated the IL-15-specific gene set in a mouse model of bystander CD8^+^ T-cell activation, in which OT-1 bystander memory cells are activated by PR8 IAV infection, with or without SIINFEKL co-stimulation.^20^ We identified the upregulated DEGs in OT-1 memory cells during PR8 infection, compared to in those cells during PR8 infection and SIINFEKL co-stimulation, and found that the IL-15-specific gene set was significantly enriched among the DEGs from the PR8 mouse model (**Figure S10)**. On the other hand, the TCR-specific gene set was significantly enriched among the upregulated DEGs in OT-1 memory cells during PR8 infection and SIINFEKL co-stimulation, compared to those cells during PR8 infection without SIINFEKL co-stimulation. These findings demonstrate that the IL-15-specific gene set and TCR-specific gene set are good indicators of IL-15-induced bystander activation and TCR-induced conventional activation, respectively. We further conclude that the IL-15-specific gene set, which was upregulated by IL-15 and counteracted by concurrent TCR stimulation, can be used to identify memory CD8^+^ T cells that are activated by an IL-15-induced bystander mechanism under various conditions.

## Discussion

The characteristic features of IL-15-induced bystander activation—including NK-like cytotoxicity and the upregulation of NKG2D and CCR5—are differentially regulated by IL-15 and TCR stimulation in memory CD8^+^ T cells.^2,8,19^ However, the molecular characteristics and regulatory mechanisms governing this phenomenon have not been previously explored. Here, we identified a specific set of genes related to NK-cell-mediated cytotoxicity and IFN response, which was upregulated by IL-15 but restrained by concurrent TCR stimulation in memory CD8^+^ T cells. Moreover, we used ionomycin to demonstrate that Ca^2+^-calcineurin signaling restrained characteristic features of IL-15-induced bystander activation and NK-like cytotoxicity. Furthermore, we showed that NFATc1 suppressed IL-15-induced bystander activation via AP-1 binding. Consistent with these results, calcineurin inhibitors increased the IL-15-induced NKG2D upregulation on memory CD8^+^ T cells in the presence of TCR signals. We further defined the genes that were upregulated by IL-15 and suppressed by concurrent TCR stimulation as the gene set of IL-15-induced bystander activation. This gene signature was significantly enriched in bystander CD8^+^ T cells from patients with acute HAV infection, a prototype disease of bystander CD8^+^ T-cell activation.

IL-15 is a homeostatic cytokine that plays essential roles in the development, proliferation, and maintenance of memory CD8^+^ T cells.^25,26,27,28,29^ However, accumulating evidence has expanded our understanding of IL-15 from a homeostatic cytokine to a potent inducer of bystander CD8^+^ T-cell activation.^2^ Previous studies have shown that IL-15 itself elicits multiple functions of memory CD8^+^ T cells, independently of TCR signals, including proliferation, synthesis of effector molecules, and cytotoxicity.^30,31,32^ It was later demonstrated that IL-15 can activate bystander CD8^+^ T cells and enhance their NK-like cytotoxic features that contribute to host tissue damage.^8,18^ However, limited information is available regarding the specific transcriptional signatures induced by bystander activation in memory CD8^+^ T cells, which are distinguished from those induced by TCR stimulation. Notably, the specific transcriptional profile of IL-15-induced bystander activation has not been previously profiled in human memory CD8^+^ T cells.

In the present study, we found a unique set of genes upregulated by IL-15 and downregulated by concurrent TCR stimulation, which were enriched with genes associated with NK-cell-mediated cytotoxicity and IFN response. This finding supports previous reports of NK-like cytotoxic activity of IL-15-activated memory CD8^+^ T cells.^5,8,10,31,33,34^ Moreover, the enrichment of IFN response genes among genes upregulated by IL-15 and downregulated by concurrent TCR stimulation also confirms our previous findings that IFN response genes were enriched in bystander-activated memory CD8^+^ T cells from a mouse model.^20^ The gene set of IL-15-induced bystander activation, which was upregulated by IL-15 and counteracted by concurrent TCR stimulation, can be used to identify memory CD8^+^ T cells that are activated by an IL-15-induced bystander mechanism under various conditions.

To validate the IL-15-induced bystander activation gene set in a human disease setting, we performed a scRNA-seq analysis that incorporated dCODE MHC-I multimers. Using this approach, we analyzed single-cell transcriptomes of HAV-specific and bystander (HCMV-, EBV-, and IAV-specific) CD8^+^ T cells from PBMCs of patients with acute HAV infection. Importantly, the IL-15-induced bystander activation gene set that was upregulated by TCR-independent IL-15 stimulation was significantly enriched in bystander CD8^+^ T cells from these patients. However, this enrichment was not observed in HAV-specific CD8^+^ T cells during acute HAV infection. This finding demonstrates that gene set will be useful for distinguishing IL-15-induced bystander CD8^+^ T-cell activation from TCR-mediated conventional activation. This validated IL-15-induced bystander activation gene set provides a compelling opportunity to explore its use in various other diseases with elevated IL-15 levels that correlate with disease severity, in which the phenomenon of TCR-independent CD8^+^ T-cell activation is implicated.^2,35^ For example, the IL-15-induced bystander activation gene set could be used to study alopecia areata,^33^ in which virtual memory CD8^+^ T cells activated by IL-15 exert NKG2D-dependent NK-like cytotoxicity. Identifying and analyzing TCR-independent bystander CD8^+^ T-cell activation in other diseases could enhance our understanding of the roles of bystander-activated CD8^+^ T cells, elucidating their potential contribution to shaping disease outcomes. Moreover, future research could uncover potential therapeutic targets, which could be used to modulate the immunopathological role of TCR-independent bystander CD8^+^ T-cell activation in various diseases.

Binding of IL-15 to its cellular receptor delivers intracellular signals,^36,37^ some of which (e.g., mTOR and MAPK signaling pathways) overlap with those induced by TCR stimulation.^38^ Due to the overlapping components of signaling pathways, previous reports have focused on the role of IL-15 in amplifying TCR signaling, thereby enhancing various TCR-mediated effector functions of CD8^+^ T cells.^39,40,41^ However, the Ca^2+^-calcineurin signaling pathway discriminates between TCR and IL-15 signals. Upon TCR ligation, Ca^2+^ is released into the cytosol, leading to a rapid influx of extracellular Ca^2+^ until its level is highly sustainable for T-cell activation.^42^ The accumulated cytosolic Ca^2+^ binds to calmodulin, leading to a subsequent conformational change in calmodulin for its interaction with the protein phosphatase calcineurin.^43^ This interaction dephosphorylates NFATc1, which leads to its translocation into the nucleus, where it binds to AP-1 and regulates various T-cell functions.^44, 45^ However, IL-15 stimulation does not activate the Ca^2+^-calcineurin signaling pathway, as indicated by our present finding that calcineurin inhibitors had no effect on IL-15-induced CD8^+^ T-cell activation. Instead, Ca^2+^ signals suppress characteristic features of the IL-15-induced bystander activation of memory CD8^+^ T cells, including upregulation of NKG2D the IL-15-induced bystander activation gene set, and NK-like cytotoxic activity.

AP-1 is a transcription factor induced by diverse immune signaling pathways, including IL-15.^36^ It typically exists as a dimeric complex comprising proteins from the JUN, FOS, ATF, and MAF families.^46^ Notably, c-JUN and c-FOS are essential for inducing NKG2D expression by upregulating DAP10 expression.^47^ In fact, in our study, we demonstrated that IL-15 upregulates *HCST* mRNA expression and NKG2D surface expression on memory CD8^+^ T cells. Moreover, we found that AP-1 drives the IL-15-induced upregulation of NKG2D on memory CD8⁺ T cells. Given that NFATc1 binds AP-1 to form a transcriptional complex,^44^ we further investigated whether this interaction under concurrent TCR stimulation abrogates the action of AP-1 regarding the IL-15-induced upregulation of NKG2D and other NK-related genes, including *HCST*. Notably, TCR stimulation promotes NFATc1 binding to AP-1, thereby preventing the AP-1-dependent transcription of key genes involved in IL-15-induced NK-like bystander activation. Our findings reveal a crucial regulatory mechanism, in which TCR signaling counteracts cytokine-induced T-cell activation. This discovery offers new insights into the balance between antigen-specific adaptive and cytokine-induced innate-like T-cell activation.

Calcineurin inhibitors, such as FK506 and CsA, are widely used as immunosuppressants in organ transplantation settings and for the treatment of autoimmune diseases.^48^ However, previous studies have reported that calcineurin inhibitors do not affect the cytokine-induced production of IL-17A and IFN-γ from mouse memory-like CD4^+^ T cells.^49^ Consistently, we demonstrated that calcineurin inhibitors failed to suppress the IL-15-induced activation and proliferation of memory CD8^+^ T cells. Moreover, calcineurin inhibitors paradoxically upregulated the IL-15-induced NKG2D expression on memory CD8^+^ T cells in the presence of TCR co-stimulation. This suggests that calcineurin inhibitors may enhance NK-like cytotoxic activity of memory CD8^+^ T cells, and exacerbate CD8^+^ T cell-mediated immunopathology in IL-15-associated pathological conditions. The data from our current study suggests that immunosuppressive regimens must be carefully selected for the treatment of chronic inflammatory diseases, depending on the mechanism of CD8^+^ T-cell activation.

The recent recognition that bystander activation of memory CD8^+^ T cells can potentially impact disease outcomes has attracted great attention. Our present study highlights characteristic features and regulatory mechanisms underlying IL-15-induced bystander activation in human memory CD8^+^ T cells. We demonstrated that IL-15-induced bystander activation exhibits NK-like qualities, rather than being an inherent aspect of the adaptive immune response. Moreover, we found that TCR stimulation plays a role in preventing such antigen-nonspecific NK-like activation through Ca^2+^-calcineurin-NFATc1 signaling, thereby preserving the intrinsic nature of the adaptive immune response. Increasing our understanding of the mechanisms underlying the IL-15-induced bystander activation of memory CD8^+^ T cells will have important clinical implications for the appropriate treatment of patients whose tissues are damaged by CD8^+^ T cell-mediated immunopathogenesis.

## Acknowledgement

This work was supported by the Bio&Medical Technology Development Program of the National Research Foundation (NRF) funded by the Korean government (MSIT) (RS-2024-00439160 and RS-2024-00512914) and by the Institute for Basic Science (IBS), Republic of Korea (IBS-R801-D2).

## Author contributions

H.L. and E.C.S. designed the study. H.S.E. collected and provided human clinical samples and information. H.L., S.Y.K., S.H.K., and K.H.K. performed the experiments and collected data. H.L., S.Y.K., S.H.K., H.Y., S.J., K.H.K., C.G.K., J.Y.K., H.D.K., J.W.H., S.H.P., and E.C.S were involved in data analysis and interpretation. H.L., S.Y.K., and H.Y. performed statistical analysis. H.L., S.Y.K., and E.C.S. drafted the manuscript with comments from all authors.

## INCLUSION AND DIVERSITY

We support inclusive, diverse, and equitable conduct of research.

## STAR★METHODS

### KEY RESOURCES TABLE

| REAGENT or RESOURCE | SOURCE | IDENTIFIER |
| --- | --- | --- |
| **Antibodies** |  |  |
| BV510 Anti-human CD3 (clone UCHT1), | BD Biosciences | Cat# 563109<br>RRID:AB_2732053 |
| BV605 Anti-human CD8 (clone SK1) | BD Biosciences | Cat# 564116<br>RRID:AB_2869551 |
| BV711 Anti-human Ki-67 (clone B56) | BD Biosciences | Cat# 563755<br>RRID:AB_2738406 |
| PerCP-Cy5.5 Anti-human CD4 (clone RPA-T4) | BD Biosciences | Cat# 560650<br>RRID:AB_1727476 |
| AlexaFluor 700 Anti-human CD3 (clone UCHT1) | BD Biosciences | Cat# 557943<br>RRID:AB_396952 |
| PerCP-Cy5.5 Anti-human CD14 (clone M5E2) | BD Biosciences | Cat# 550787<br>RRID:AB_393884 |
| PerCP-Cy5.5 Anti-human CD19 (clone H fIB19) | BD Biosciences | Cat# 561295<br>RRID:AB_10644017 |
| PE Anti-human NKG2D (clone 1D11) | BD Biosciences | Cat# 561815<br>RRID:AB_10896282 |
| PE-CF594 Anti-human CD14 (clone MPHIP9) | BD Biosciences | Cat# 562335<br>RRID:AB_11153663 |
| PE-CF594 Anti-human CD19 (clone HIB19) | BD Biosciences | Cat# 562294<br>RRID:AB_11154408 |
| Pe-Cy7 Anti-human CD45RA (clone HI100) | BD Biosciences | Cat# 560675<br>RRID:AB_1727498 |
| AlexaFluor 700 Anti-human CD45RA (clone HI100) | BD Biosciences | Cat# 560673<br>RRID:AB_1727496 |
| APC Anti-human HLA-DR (clone G46-6) | BD Biosciences | Cat# 559866<br>RRID:AB_398674 |
| BV650 Anti-Human HLA-DR (clone L203) | BD Biosciences | Cat# 752489<br>RRID:AB_2917482 |
| AlexaFluor 700 Anti-human CD3 (clone UCHT1) | BD Biosciences | Cat# 557943<br>RRID:AB_396952 |
| AlexaFluor 700 Anti-human CD38 (clone HIT2) | BD Biosciences | Cat# 560676<br>RRID:AB_1727472 |
| APC-H7 Anti-human CD8 (clone SK1) | BD Biosciences | Cat# 560273<br>RRID:AB_1645482 |
| FITC Anti-human CD16 (clone B73.1) | BD Biosciences | Cat# 561308<br>RRID:AB_10644202 |
| BV421 anti-human NKG2D (clone 1D11) | Biolegend | Cat# 320821<br>RRID:AB_2566510 |
| PerCP/cy5.5 Anti-human CD197 (CCR7) (clone G043H7) | Biolegend | Cat# 353220<br>RRID:AB_10916121 |
| APC/Cyanine7 Anti-human HLA-DR (clone L243) | Biolegend | Cat# 307618<br>RRID:AB_493586 |
| TotalSeq™-C0034 anti-human CD3 Antibody (clone UCHT1) | Biolegend | Cat# 300479<br>RRID: AB_2800723 |
| TotalSeq™-C0045 anti-human CD4 Antibody (clone SK3) | Biolegend | Cat# 344651<br>RRID:AB_2800921 |
| TotalSeq™-C0046 anti-human CD8 Antibody (clone SK1) | Biolegend | Cat# 344753<br>RRID:AB_2800922 |
| TotalSeq™-C0047 anti-human CD56 (NCAM) Antibody (clone 5.1H11) | Biolegend | Cat# 362559<br>RRID:AB_2801002 |
| TotalSeq™-C0051 anti-human CD14 Antibody (clone 63D3) | Biolegend | Cat# 367137<br>RRID:AB_2810576 |
| TotalSeq™-C0063 anti-human CD45RA Antibody | Biolegend | Cat# 304163 |
| (clone HI100) |  | RRID:AB_2800764 |
| TotalSeq™-C0083 anti-human CD16 Antibody<br>(clone 3G8) | Biolegend | Cat# 302065<br>RRID:AB_2800738 |
| TotalSeq™-C0085 anti-human CD25 Antibody<br>(clone BC96) | Biolegend | Cat# 302649<br>RRID: AB_2800745 |
| TotalSeq™-C0087 anti-human CD45RO Antibody<br>(clone UCHL1) | Biolegend | Cat# 304259<br>RRID: AB_2800766 |
| TotalSeq™-C0088 anti-human CD279 (PD-1)<br>Antibody (clone EH12.2H7) | Biolegend | Cat# 329963<br>RRID: AB_2800862 |
| TotalSeq™-C0089 anti-human TIGIT (VSTM3)<br>Antibody (clone A15153G) | Biolegend | Cat# 372729<br>RRID: AB_2801021 |
| TotalSeq™-C0101 anti-human CD335 (NKp46)<br>Antibody (clone 9 E2) | Biolegend | Cat# 331941<br>RRID: AB_2800874 |
| TotalSeq™-C0138 anti-human CD5 Antibody<br>(clone UCHT2) | Biolegend | Cat# 300637<br>RRID: AB_2800726 |
| TotalSeq™-C0139 anti-human TCR $\gamma/\delta$ Antibody<br>(clone B1) | Biolegend | Cat# 331231<br>RRID: AB_2814199 |
| TotalSeq™-C0140 anti-human CD183 (CXCR3)<br>Antibody (clone G025H7) | Biolegend | Cat# 353747<br>RRID: AB_2800949 |
| TotalSeq™-C0141 anti-human CD195 (CCR5)<br>Antibody (clone J418F1) | Biolegend | Cat# 359137<br>RRID: AB_2810570 |
| TotalSeq™-C0145 anti-human CD103 (Integrin αE)<br>Antibody (clone BerACT8) | Biolegend | Cat# 350233<br>RRID: AB_2800933 |
| TotalSeq™-C0146 anti-human CD69 Antibody<br>(clone FN50) | Biolegend | Cat# 310951<br>RRID: AB_2800810 |
| TotalSeq™-C0147 anti-human CD62L Antibody<br>(clone DREG-56) | Biolegend | Cat# 304851<br>RRID: AB_2800770 |
| TotalSeq™-C0148 anti-human CD197 (CCR7)<br>Antibody (clone G043H7) | Biolegend | Cat# 353251<br>RRID: AB_2800943 |
| TotalSeq™-C0149 anti-human CD161 Antibody<br>(clone HP-3G10) | Biolegend | Cat# 339947<br>RRID: AB_2810532 |
| TotalSeq™-C0151 anti-human CD152 (CTLA-4)<br>Antibody (clone BNI3) | Biolegend | Cat# 369621<br>RRID: AB_2801015 |
| TotalSeq™-C0152 anti-human CD223 (LAG-3)<br>Antibody (clone 11C3C65) | Biolegend | Cat# 369335<br>RRID: AB_2814327 |
| TotalSeq™-C0154 anti-human CD27 Antibody<br>(clone O323) | Biolegend | Cat# 302853<br>RRID: AB_2800747 |
| TotalSeq™-C0155 anti-human CD107a (LAMP-1)<br>Antibody (clone H4A3) | Biolegend | Cat# 328649<br>RRID: AB_2800854 |
| TotalSeq™-C0159 anti-human HLA-DR Antibody<br>(clone L243) | Biolegend | Cat# 307663<br>RRID: AB_2800795 |
| TotalSeq™-C0165 anti-human CD314 (NKG2D)<br>Antibody (clone 1D11) | Biolegend | Cat# 320837<br>RRID: AB_2800844 |
| TotalSeq™-C0168 anti-human CD57 Recombinant<br>Antibody (clone QA17A04) | Biolegend | Cat# 393321<br>RRID: AB_2801030 |
| TotalSeq™-C0169 anti-human CD366 (Tim-3)<br>Antibody (clone F38-2E2) | Biolegend | Cat# 345049<br>RRID: AB_2800925 |
| TotalSeq™-C0170 anti-human CD272 (BTLA)<br>Antibody (clone MIH26) | Biolegend | Cat# 344527<br>RRID: AB_2800919 |
| TotalSeq™-C0176 anti-human CD39 Antibody<br>(clone A1) | Biolegend | Cat# 328237<br>RRID: AB_2800853 |
| TotalSeq™-C0179 anti-human CX3CR1 Antibody<br>(clone K0124E1) | Biolegend | Cat# 355705<br>RRID: AB_2800960 |
| TotalSeq™-C0224 anti-human TCR $\alpha/\beta$ Antibody<br>(clone IP26) | Biolegend | Cat# 306743<br>RRID: AB_2800793 |
| TotalSeq™-C0250 anti-mouse/human KLRG1<br>(MAFA) Antibody (clone 2F1/KLRG1) | Biolegend | Cat# 138433<br>RRID: AB_2800649 |
| TotalSeq™-C0355 anti-human CD137 (4-1BB) | Biolegend | Cat# 309839 |
| Antibody (clone 4B4-1) |  | RRID: AB_2800807 |
| TotalSeq™-C0368 anti-human CD226 (DNAM-1)<br>Antibody (clone 11A8) | Biolegend | Cat# 338337<br>RRID: AB_2800899 |
| TotalSeq™-C0386 anti-human CD28 Antibody<br>(clone CD28.2) | Biolegend | Cat# 302963<br>RRID: AB_2800751 |
| TotalSeq™-C0389 anti-human CD38 Antibody<br>(clone HIT2) | Biolegend | Cat# 303543<br>RRID: AB_2800758 |
| TotalSeq™-C0390 anti-human CD127 (IL-7Rα)<br>Antibody (clone A019D5) | Biolegend | Cat# 351356<br>RRID: AB_2800937 |
| TotalSeq™-C0420 anti-human CD158<br>(KIR2DL1/S1/S3/S5) Antibody (clone HP-MA4) | Biolegend | Cat# 339517<br>RRID: AB_2814252 |
| TotalSeq™-C0581 anti-human TCR Vα7.2 Antibody<br>(clone 3C10) | Biolegend | Cat# 351735<br>RRID: AB_2810556 |
| TotalSeq™-C0584 anti-human TCR Vα24-Jα18<br>(iNKT cell) Antibody (clone 6B11) | Biolegend | Cat# 342925<br>RRID: AB_2810529 |
| TotalSeq™-C0592 anti-human CD158b<br>(KIR2DL2/L3, NKAT2) Antibody (clone DX27) | Biolegend | Cat# 312619<br>RRID: AB_2819944 |
| TotalSeq™-C0599 anti-human CD158e1 (KIR3DL1,<br>NKB1) Antibody (clone DX9) | Biolegend | Cat# 312725<br>RRID: AB_2814161 |
| TotalSeq™-C0801 anti-human CD337 (NKp30)<br>Antibody (clone P30-15) | Biolegend | Cat# 325219<br>RRID: AB_2800851 |
| TotalSeq™-C0802 anti-human CD336 (NKp44)<br>Antibody (clone P44-8) | Biolegend | Cat# 325119<br>RRID: AB_2810484 |
| TotalSeq™-C0804 anti-human CD186 (CXCR6)<br>Antibody (clone K041E5) | Biolegend | Cat# 356023<br>RRID: AB_2876677 |
| TotalSeq™-C0867 anti-human CD94 Antibody<br>(clone DX22) | Biolegend | Cat# 305523<br>RRID: AB_2814143 |
| TotalSeq™-C0896 anti-human CD85j (ILT2)<br>Antibody (clone GHI/75) | Biolegend | Cat# 333725<br>RRID: AB_2814226 |
| TotalSeq™-C1021 anti-human GPR56 Antibody (clone 4C3) | Biolegend | Cat# 391907<br>RRID: AB_2832737 |
| CD3 monoclonal antibody (clone OKT3)<br>Functional Grade | Invitrogen | Cat# 16-0037-85<br>RRID: AB_468854 |
| Fluorescein-conjugated Anti-human CCR7 (clone 150503) | R&D Systems | Cat# FAB197F<br>RRID: AB_2259847 |
| Anti-c-Jun antibody [E254] - ChIP Grade | Abcam | Cat# ab32137<br>RRID: AB_731608 |
| Rabbit Anti-Mouse IgG H&L | Abcam | Cat# ab46540<br>RRID: AB_2614925 |
| Chemicals, Peptides and Recombinant Proteins |  |  |
| Brilliant Stain Buffer | BD Biosciences | Cat# 563794 |
| 7-AAD | BD Biosciences | Cat# 559925 |
| Recombinant human IL-2 protein | Peprotech | Cat# 200-0 |
| Recombinant human IL-4 protein | Peprotech | Cat# 200-04 |
| Recombinant human IL-7 protein | Peprotech | Cat# 200-07 |
| Recombinant human IL-9 protein | Peprotech | Cat# 200-09 |
| Recombinant human IL-15 protein | Peprotech | Cat# 200-15 |
| Recombinant human IL-21 protein | Peprotech | Cat# 200-21 |
| Dimethyl sulfoxide | Sigma-Aldrich | Cat# R8418 |
| Cyclosporin A | Sigma-Aldrich | Cat# SML1018 |
| PMA | Sigma-Aldrich | Cat# P1585 |
| Ionomycin calcium salt | Sigma-Aldrich | Cat# I3909 |
| SP600125 | Sigma-Aldrich | Cat# S5567 |
| Formaldehyde solution | Sigma-Aldrich | Cat# F8775 |
| Tacrolimus (FK506) | Selleckchem | Cat# S5003 |
| Everolimus (RAD001) | Selleckchem | Cat# S1120 |
| SR 11302 | R&D Systems | Cat# 2476 |
| Lymphocyte Separation Medium | Corning | Cat# 25-072-CV |
| Trizol reagent | Invitrogen | Cat# 15596018 |
| To-Pro<sup>TM</sup>-3 iodide | Invitrogen | Cat# T3605 |
| Pierce<sup>TM</sup> DSG, No-Weigh<sup>TM</sup> Format | Thermo Fisher | Cat# 20593 |
| Dynabeads<sup>TM</sup> Protein A for Immunoprecipitation | Thermo Fisher | Cat# 10001D |
| Critical Commercial Assays |  |  |
| LIVE/DEAD fixable aqua dead cell stain kit | Invitrogen | Cat# L34957 |
| LIVE/DEAD fixable Red dead cell stain kit | Invitrogen | Cat# L34972 |
| Intracellular fixation & permeabilization buffer set | Invitrogen | Cat# 88-8824-00 |
| Neon<sup>TM</sup> NxT Electroporation System 100-μL Kit | Invitrogen | Cat# N10025 |
| CellTrace Violet Cell Proliferation Kit | Invitrogen | Cat# C34557 |
| TaqMan<sup>TM</sup> Gene Expression Assay (FAM) (Gene Symbol: HCST) | Applied Biosystems | Cat# 4351370 Hs00367159_m1 |
| TaqMan<sup>TM</sup> Gene Expression Assay (FAM) (Gene Symbol: B2M) | Applied Biosystems | Cat# 4331182 Hs00187842_m1 |
| TaqMan Gene Expression Master Mix | Applied Biosystems | Cat# 4369016 |
| RNase A, DNase and protease-free | Thermo Fisher | Cat# EN0531 |
| Anti-FITC microbeads | Miltenyi Biotec | Cat# 130-048-701 |
| CD8 MicroBeads, human | Miltenyi Biotec | Cat# 130-045-201 |
| Human CD8<sup>+</sup> T cell isolation kit | Miltenyi Biotec | Cat# 130-096-495 |
| Ribospin total RNA purification kit | Geneall | Cat# 130-090-101 |
| cDNA synthesis master mix | LeGene | Cat# 6290-20 |
| Diluent C for general membrane labeling | Sigma-Aldrich | Cat# CGLDIL |
| PKH26 Red Fluorescent Cell Linker Kit for General Cell Membrane Labeling | Sigma-Aldrich | Cat# PKH26GL |
| proteinase K | New England Biolabs | Cat# P8107S |
| Recombinant DNA |  |  |
| pcDNA3.1_empty | Genescript | N/A |
| pcDNA3.1_NFATc1_WT | Genescript | N/A |
| pcDNA3.1_NFATc1_T551_MT | Genescript | N/A |
| siRNAs |  |  |
| Silencer™ Select Negative Control No. 1 siRNA | Invitrogen | Cat# 4390844 |
| Silencer™ Select c-Fos siRNA | Invitrogen | Cat# 4392420 |
| Silencer™ Select c-Jun siRNA | Invitrogen | Cat# 4392420 |
| Cell line |  |  |
| K562 | ATCC | Cat# CCL-243 |
| Software and Algorithms |  |  |
| FlowJo (version 10.7.2) | Tree Star | https://www.flowjo.com/solutions/flowjo/downloads |
| Prism (version 8.4) | GraphPad | http://www.graphpad.com/scientific-software/prism/ |
| R v.4.1.0 | The R foundation | N/A |
| Cell Ranger | R package | Version 3.1.0 |
| Seurat package | R package | Version 3.2.2 |
| STAR aligner | R package | Version 2.5.1b |
| DESeq2 | R package | Version 1.24.0 |
| ComplexHeatmap | R package | Version 2.15.4 |
| Bowtie2 | R package | Version 2.2.9 |
| limma | R package | Version 4.3 |
| fgsea | R package | Version 1.22.0 |
| biomaRt | R package | Version 2.50.0 |
| GSEA | PMID:17644558 | http://www.gsea-msigdb.org/gsea/msigdb/index.jsp |
| enrichR database | Maayanlab | https://maayanlab.c |
|  |  | loud/Enrichr/ |
| Dextramers |  |  |
| HLA-A*0201 (NLVPMVATV), APC CMV Dextramer | Immudex | WB2132- APC-150 |
| HLA-A*0201 (GLCTLVAML), APC EBV Dextramer | Immudex | WB2130- APC-150 |
| HLA-A*0201 LLYNCCYHV (HAV) dCODE<br>Dextramer® grade (Gold) | Immudex | dCODELLYNCCYHV -<br>50T |
| HLA-A*0201 NLVPMVATV (CMV) dCODE<br>Dextramer® grade (Gold) | Immudex | dCODENLVPMVATV -<br>50T |
| HLA-A*0201 GLCTLVAML (EBV) dCODE<br>Dextramer® grade (Gold) | Immudex | dCODEGLCTLVAML -<br>50T |
| HLA-A*0201 GILGFVFTL (IAV) dCODE Dextramer®<br>grade (Gold) | Immudex | dCODEGILGFVFTL50<br>T |

### RESOURCE AVAILABILITY

#### Lead contact

Further information and requests for resources and reagents should be directed to and will be fulfilled by the lead contact, Eui-Cheol Shin.

#### Materials availability

The study did not produce novel, distinctive materials or substances.

#### Data and code availability

All analyzed sequencing data will be available online. Bioinformatic analyses were performed using open-source software and in-house scripts in R (version 4.1.0), which are available from the corresponding author upon reasonable request.

### STUDY PARTICIPANT DETAILS

#### Human blood samples

Peripheral blood samples were obtained from healthy volunteers from KAIST Clinic Pappalardo Center, who were not taking any immunosuppressive medications. Peripheral blood samples were also collected from patients with acute HAV infection and also during the convalescent phase, who at admission exhibited clinical and laboratory features of acute hepatitis A (AHA), including seropositivity for anti-HAV IgM and/or IgG antibodies. PBMCs were isolated from whole blood by density gradient centrifugation using Lymphocytes Separation Medium (Corning). The isolated PBMCs were cryopreserved in freezing media containing fetal bovine serum (Corning) with 10% dimethyl sulfoxide (Sigma-Aldrich) until their use. This study was conducted in accordance with the principles of the Declaration of Helsinki, and was approved by the institutional review board of each involved institution (KAIST: KH2018-118 and Chungnam National University Hospital: 2019-03-066-025). Each healthy volunteer and patient with HAV infection gave their written informed consent prior to inclusion in the study.

### METHOD DETAILS

#### Cell sorting

CD8^+^ T cells were negatively isolated from PBMCs of healthy donors and patients with HAV infection using a CD8^+^ T-cell isolation kit (Miltenyi Biotec), according to the manufacturer’s instructions. Isolated CD8^+^ T cells were stained with FITC-conjugated anti-CCR7 antibody (R&D systems), and then further stained with anti-FITC microbeads (Miltenyi Biotec), following the manufacturer’s instructions for negatively isolating CCR7^−^ memory CD8^+^ T cells.

#### In vitro stimulation and proliferation assay

PBMCs were cultured in RPMI 1640 medium (Corning) supplemented with 10% fetal bovine serum (Corning) and 1% penicillin/streptomycin (Sigma-Aldrich). PBMCs were cultured for 48 hours with IL-2 (10 ng/ml; Peprotech), IL-4 (10 ng/ml; R&D systems), IL-7 (10 ng/ml; Peprotech), IL-9 (10 ng/ml; Peprotech), IL-15 (10 ng/ml; Peprotech), or IL-21 (10 ng/ml; Peprotech), and we examined the NKG2D expression among CD8^+^ T cells and their naïve and memory subsets. Sorted CCR7^−^ memory CD8^+^ T cells were cultured with IL-15 (10 ng/ml), anti-CD3 (coated, 1 µg/ml; Invitrogen), PMA (10 ng/ml; Sigma-Aldrich), ionomycin (500 ng/ml; Sigma-Aldrich), or combinations of these additives, for 48 hours followed by analysis of NKG2D and granzyme B expressions, or for 96 hours followed by analysis of CD38 and HLA-DR expressions.

For a proliferation assay, sorted CCR7^−^ memory CD8^+^ T cells were stained with CellTrace Violet (Invitrogen) following the manufacturer’s instructions. CellTrace Violet-labelled cells were cultured with IL-15 (10 ng/ml), anti-CD3 (coated, 1 µg/ml), or the combination of both for 96 hours, and then we measured their proliferation and Ki-67 expression.

We also tested the effects of chemical inhibitors. Sorted CCR7^−^ memory CD8^+^ T cells were pre-treated with everolimus (10 μM; Selleckchem), FK506 (10 ng/ml; Selleckchem), CsA (100 ng/ml; Selleckchem), SP600125 (10 μM; Selleckchem), or SR11302 (10 μM; Selleckchem) for one hour, followed by stimulation with IL-15, anti-CD3, or the combination of both.

#### Flow cytometry

Cells were stained with a mixture of brilliant stain buffer (BD Biosciences) and fluorochrome-conjugated antibodies that bind to surface markers, for 20 min at 4°C. Dead cells were excluded from analysis using the Live/Dead Cell Stain Kit (Invitrogen). To stain intracellular molecules, surface-stained cells were fixed and permeabilized (Invitrogen) for 20 min at 4°C. Subsequently, these cells were stained with fluorochrome-conjugated antibodies targeting intracellular molecules, for 20 min at 4°C. To analyze HCMV- and EBV-specific CD8^+^ T cells, PBMCs were stained with MHC-I dextramers (Immudex) specific for HCMV (HCMV pp65 495–504; NLVPMVATV) and EBV (EBV BMLF-1 259–267; GLCTLVAML), and fluorochrome-conjugated monoclonal antibodies for cell surface proteins. Multicolor flow cytometry was performed using LSR II, LSR Fortessa, and FACSymphony A3 instruments (BD Biosciences), and data were analyzed using FlowJo software (Treestar).

#### Quantitative PCR

Total RNA was isolated using Ribospin^TM^ total RNA purification kit (GeneAll). Complementary DNA (cDNA) was synthesized using cDNA synthesis master mix (LeGene). TaqMan Gene Expression Assays (Applied Biosystems) were used to quantify *HCST* mRNA expression, which was normalized to the endogenous control *B2M* expression. Quantitative PCR was performed on a CFX96 Real-Time PCR Detection System and analyzed using CFX Manager^TM^ IVD Software (Bio-Rad).

#### Bulk RNA sequencing

Bulk RNA-seq was performed on sorted CCR7^−^ memory CD8^+^ T cells that were stimulated with IL-15, anti-CD3, or the combination of both for 48 hours; or with IL-15, ionomycin, or the combination of both for 12 hours. Total RNA was extracted using TRIZOL reagent (Invitrogen). RNA quality and quantity were evaluated using an Agilent 2100 Bioanalyzer with an RNA 6000 Nano Chip (Agilent Technologies) and an ND-2000 spectrophotometer (Thermo Fisher Scientific), respectively. An RNA library was prepared using a QuantSeq 3’ mRNA-seq Library Prep Kit (Lexogen). An individual set of 500 ng total RNA was prepared, an oligo-dT primer containing an Illumina-compatible sequence at its 5’ end was hybridized to the RNA, and reverse transcription was performed. After degradation of the RNA template, second strand synthesis was processed using a random primer with an Illumina-compatible linker sequence at its 5’ end. Magnetic beads were used to purify the double-stranded library, and to eliminate reactive components. The library was amplified for the addition of complete adapter sequences needed for cluster generation and, finally, the amplified library was purified from PCR components.

High-throughput sequencing was performed by single-end 75 sequencing using NextSeq 500 (Illumina). Alignment of the QuantSeq 3’ mRNA-Seq reads based on a human reference sequence (hg19) was performed using Bowtie2.^50^ Gene counts were normalized according to the library size, and DEGs were analyzed using DESeq2 (v.1.24.0).^51^ An experimental batch was corrected using the removeBatchEffect function from limma.^52^ DEGs were identified based on two cut-off values: an adjusted *P* value < 0.05, and an absolute log2 fold change > 1. GSEA was performed using Broad Institute Software (http://www.gsea-msigdb.org/gsea/index.jsp). For upregulated DEGs, pathway analysis was performed using the enrichR database (https://maayanlab.cloud/Enrichr/). The following pathway databases were utilized in this study: MSigDB Hallmark 2020 (interferon gamma response, interferon alpha response, and TNF-alpha signaling via NF-kB), Reactome 2022 (interferon signaling R-HAS-913531, cell cycle R-HAS-1640170, rRNA processing, cytokine signaling in immune system R-HAS-1280215, innate immune system R-HAS-168249, and interferon alpha/beta signaling R-HAS-909733), KEGG 2021 Human (natural killer cell-mediated cytotoxicity, T-cell receptor signaling pathway, and chemokine signaling pathway), Elsevier Pathway Collection (natural killer cell precursor -> NK cell surface expression markers, natural killer cell activation), NCI-Nature 2016 (calcineurin-regulated NFAT-dependent transcription in lymphocyte), and BioPlanet 2019 (regulation of NFAT transcription factors, MYC active pathway, EGFR1 pathway, T-cell receptor calcium pathway).

#### Cytotoxicity assay

Sorted CD8^+^ T cells from patients with acute HAV infection were stimulated with IL-15 (10 ng/ml) alone or in combination with ionomycin (500 ng/ml) for 24 hours. K562 target cells were labelled with PKH26 (Sigma-Aldrich) following the manufacturer’s instructions, and co-cultured with stimulated CD8^+^ T cells at an effector:target cell ratio of 20:1, for 12 hours. To-Pro-3-iodide (0.5 μM; Invitrogen) was added to the samples, and cells were immediately analyzed by flow cytometry. To obtain the background or maximum To-Pro-3-Iodide staining, target cells were incubated in medium without effector cells or detergent, respectively. The specific cytotoxicity was calculated by extracting the percentage of To-Pro-3-Iodide+ PKH26+ cells cultured without effector cells, from the percentage of To-Pro-3-Iodide+ PKH26^+^ cells in co-culture.

#### siRNA transfection

PBMCs were transfected with c-Jun- or c-Fos-specific small interfering RNA (siRNA) (Invitrogen) or negative control siRNA (Invitrogen) using the Neon NxT electroporation system (Invitrogen). Transfection was performed with 200 nM siRNA at 2000 V for three 10 ms pulses. Following transfection, PBMCs were cultured in RPMI 1640 medium (Corning) supplemented with 10% fetal bovine serum (Corning) without antibiotics for 4 days. Subsequently, PBMCs were treated with IL-15 (10 ng/ml) for 48 hours, and expression of NKG2D was analyzed on CCR7^−^ memory CD8^+^ T cells by flow cytometry.

#### NFATc1 plasmids transfection

pcDNA3.1 plasmid containing wild-type and T551A mutant^24^ NFATc1 were purchased from Genescript. PBMCs were transfected with empty pcDNA3.1 plasmids or plasmids containing wild-type and T551A mutant NFATc1 using the Neon NxT electroporation system. Transfection was performed with 5 μg of plasmids at 2000 V for a single 20 ms pulse. Subsequently, PBMCs were cultured with IL-15 (10 ng/ml) for 48 hours and expression of NKG2D was analyzed on CCR7^−^ memory CD8^+^ T cells by flow cytometry.

#### ChIP-qPCR

Sorted CCR7^−^ memory CD8^+^ T cells were stimulated with IL-15 (10 ng/ml) or IL-15 with anti-CD3 (coated, 1 µg/ml) for 48 hours. Cells were double cross-linked with 2mM DSG (Thermo Fisher) and 1% formaldehyde (Sigma-Aldrich) at room temperature for 35 min and 10 min, respectively. Cells were then lysed with 1% SDS buffer (1% SDS, 10 mM EDTA, 50 mM Tris pH 8.1) and chromatin was fractionated by sonication (Covaris, S220). The lysate was diluted with dilution buffer and incubated with pre-coated Dynabeads Protein A (Thermo Fisher) with c-Jun antibody (Abcam) and IgG (Abcam) as negative control overnight at 4°C with rotation. A portion of diluted lysate was stored to quantify the total amount of DNA in each sample as input control. The immunoprecipitated chromatin was washed four times with RIPA buffer (140 mM NaCl, 1 mM EDTA, 0.5 Mm EGTA, 1% Triton X-100, 0.1% SDS, 0.1% sodium deoxycholate, 10 mM Tris pH 8.0) and incubated with RNase A (Thermo Fisher) and proteinase K (New England Biolabs) overnight at 65°C to reverse crosslink. The immunoprecipitated DNA was purified by AMPure XP beads (Beckman, A63881) and analyzed by Real-Time PCR using primers designed based on open chromatin regions identified from publicly available ATAC-seq data^53^ and AP-1 ChIP data^54^ (*HCST*: FW – CAGCAAATTTTCTTGGCCCTACCTC, RV – GTTACTGCCTTTGGTGAACGA; *IFITM3*: FW – GGGGGAATCTCAAAAGGCCA, RV – TGGGCCCATGTTCTCAAGAC; *CCR5*: FW – TTGGGTGGTGAGCATCTGTG, RV – TGTAGAGGGGGATCCTGGAC).

#### scRNA-seq library preparation and sequencing

Sorted CD8^+^ T cells from PBMCs of patients (*n* = 5) at acute and convalescent stages of HAV infection and healthy donors (*n* = 3) were stained with PE-conjugated MHC-I dCODE dextramers (Immudex) specific for HLA-A*0201 HAV 3D_2026_ LLYNCCYHV, HLA-A*0201 HCMV pp65_495_ NLVPMVATV, HLA-A*0201 EBV BMLF1_259_ GLCTLVAML, and HLA-A*0201 IAV MP_58_ GILGFVFTL. Subsequently, these cells were sorted into dCODE-positive and -negative CD8^+^ T cells using a FACSAria II instrument (BD Biosciences). The sorted cells were mixed at a 1:9 ratio, and then further stained with CITE-seq antibodies (TotalSeq-C; Biolegend). Cell suspensions were pelleted, and scRNA-seq libraries were prepared using the Chromium Single Cell 5’ Library & Gel Bead Kit (PN-10000006) and 5’ Feature Barcode Library Kit (PN-1000020) (10X Genomics), following the manufacturer’s instructions. These libraries were sequenced at a depth of approximately 20,000 reads per cell for genes and ADTs using the NovaSeq6000 platform (Illumina).

#### Single-cell transcriptome and ADT expression analysis

Fastq files were generated by demultiplexing the sequenced data using the mkfastq function (Cell Ranger, 10X Genomics, v3.1.0). The demultiplexed fastq files for the expressions of each gene and ADT were aligned to the reference human genome (GRCH38; 10X Cell Ranger reference GRCh38 v3.0.0). The barcode sequencing of each ADT and dCODE dextramer, filtering, barcode counting, and UMI counting were performed to generate feature-barcode matrices using the count function (Cell Ranger, v3.1.0). For basic quality control, we filtered out low-quality cells with mitochondrial genes constituting >10% of their total gene expression, <300 genes or >5,500 genes, or UMI counts of <500 or >55,000. Next, we identified highly variable genes by standardizing the normalization of gene expression in each cell based on the total read count. To correct for a batch effect and harmonize the datasets, we identified ‘anchor’ vectors of genes using the FindIntegrationAnchors function, and integrated the datasets using the IntegrateData function (Seurat, v3.2.2).^55^ The integrated samples were scaled using the ScaleData function (Seurat), and PCA was conducted using the RunPCA function (Seurat) for dimensional reduction. Lastly, the cells were visualized by UMAP using the top 20 principal components (PCs) with the RunUMAP function (Seurat) for whole-cell types. Enrichment of bulk RNA-seq data on scRNA-seq data was performed by measuring the enrichment score with the Addmodulescore function (Seurat). The pseudo-bulk scRNA-seq data were obtained using the AggregateExpression function (Seurat).

#### Analysis of a public RNA-seq dataset

We re-analyzed publicly available bulk RNA-seq datasets for bystander-activated CD8^+^ T cells from a mouse model of influenza infection^20^ (SRX13530605, SRX13530604, SRX13530603, SRX13530601, and SRX13530600ref). Orthologous pairs of human and mouse genes were processed using the getLDS function (biomaRt).^56^ Enrichment of upregulated DEGs from the current study in the public data was performed using the fgsea package

#### Statistical analysis

Statistical analyses were performed using GraphPad Prism (GraphPad Software version 8.4) or R software (R Foundation for Statistical Computing). The paired t test was used to compare continuous variables. The Mann–Whitney U test was used to compare data between two unpaired groups, and the Wilcoxon signed-rank test was used to compare data between two paired groups, and one-way ANOVA with Tukey’s correction was used to compare multiple groups. Two-sided *P* values were determined in all analyses, and a *P* value < 0.05 was considered significant.

## References

1. Kaech, S.M., Wherry, E.J. & Ahmed, R. Effector and memory T-cell differentiation: implications for vaccine development. Nat Rev Immunol. 2, 251–262 (2002).

2. Lee, H., Jeong, S. & Shin, E.C. Significance of bystander T cell activation in microbial infection. Nat Immunol. 23, 13–22 (2022).

3. Maurice, N.J., Taber, A.K. & Prlic, M. The Ugly Duckling Turned to Swan: A Change in Perception of Bystander-Activated Memory CD8 T Cells. J Immunol. 206, 455–462 (2021).

4. Zhang, X.H., Sun, S.Q., Hwang, I.K., Tough, D.F. & Sprent, J. Potent and selective stimulation of memory-phenotype CD8(+) T cells in vivo by IL-15. Immunity. 8, 591–599 (1998).

5. Meresse, B. et al. Coordinated induction by IL15 of a TCR-independent NKG2D signaling pathway converts CTL into lymphokine-activated killer cells in celiac disease. Immunity. 21, 357–366 (2004).

6. Soudja, S.M., Ruiz, A.L., Marie, J.C. & Lauvau, G. Inflammatory monocytes activate memory CD8(+) T and innate NK lymphocytes independent of cognate antigen during microbial pathogen invasion. Immunity. 37, 549–562 (2012).

7. Younes, S.A. et al. IL-15 promotes activation and expansion of CD8+ T cells in HIV-1 infection. J Clin Invest. 126, 2745–2756 (2016).

8. Kim, J. et al. Innate-like Cytotoxic Function of Bystander-Activated CD8(+) T Cells Is Associated with Liver Injury in Acute Hepatitis A. Immunity. 48, 161–173 e165 (2018).

9. Sandalova, E. et al. Contribution of herpesvirus specific CD8 T cells to anti-viral T cell response in humans. PLoS Pathog. 6, e1001051 (2010).

10. Leem, G. et al. Tumour-infiltrating bystander CD8(+) T cells activated by IL-15 contribute to tumour control in non-small cell lung cancer. Thorax. 77, 769–780 (2022).

11. Chu, T. et al. Bystander-activated memory CD8 T cells control early pathogen load in an innate-like, NKG2D-dependent manner. Cell Rep. 3, 701-708 (2013).

12. Ge, C. et al. Bystander Activation of Pulmonary Trm Cells Attenuates the Severity of Bacterial Pneumonia by Enhancing Neutrophil Recruitment. Cell Rep. 29, 4236–4244 e4233 (2019).

13. Sckisel, G.D. et al. Influenza infection results in local expansion of memory CD8(+) T cells with antigen non-specific phenotype and function. Clin Exp Immunol. 175, 79–91 (2014).

14. Arkatkar, T. et al. Memory T cells possess an innate-like function in local protection from mucosal infection. J Clin Invest. 133 (2023).

15. Ostler, T., Pircher, H. & Ehl, S. "Bystander" recruitment of systemic memory T cells delays the immune response to respiratory virus infection. Eur J Immunol. 33, 1839–1848 (2003).

16. Crosby, E.J., Goldschmidt, M.H., Wherry, E.J. & Scott, P. Engagement of NKG2D on bystander memory CD8 T cells promotes increased immunopathology following Leishmania major infection. PLoS Pathog. 10, e1003970 (2014).

17. Crosby, E.J., Clark, M., Novais, F.O., Wherry, E.J. & Scott, P. Lymphocytic Choriomeningitis Virus Expands a Population of NKG2D+CD8+ T Cells That Exacerbates Disease in Mice Coinfected with Leishmania major. J Immunol. 195, 3301–3310 (2015).

18. Huang, C.H. et al. Innate-like bystander-activated CD38(+) HLA-DR(+) CD8(+) T cells play a pathogenic role in patients with chronic hepatitis C. Hepatology. 76, 803–818 (2022).

19. Seo, I.H. et al. IL-15 enhances CCR5-mediated migration of memory CD8(+) T cells by upregulating CCR5 expression in the absence of TCR stimulation. Cell Rep. 36, 109438 (2021).

20. Jeong, S. et al. IFITM3 Is Upregulated Characteristically in IL-15-Mediated Bystander-Activated CD8(+) T Cells during Influenza Infection. J Immunol. 208, 1901–1911 (2022).

21. Park Y.P. et al. Complex regulation of human NKG2D-DAP10 cell surface expression: opposing roles of the γc cytokines and TGF-β1. Blood. 118, 3019–3027 (2011).

22. Lewis, R.S. Calcium signaling mechanisms in T lymphocytes. Annu Rev Immunol. 19, 497–521 (2001).

23. Chatila, T., Silverman, L., Miller, R. & Geha, R. Mechanisms of T cell activation by the calcium ionophore ionomycin. J Immunol. 143, 1283–1289 (1989).

24. Sun, L.J., Peterson B.R. & Verdine G.L. Dual role of the nuclear factor of activated T cells insert region in DNA recognition and cooperative contacts to activator protein 1. Proceedings of the National Academy of Sciences. 94, 4919–4924 (1997).

25. Lodolce, J.P. et al. IL-15 receptor maintains lymphoid homeostasis by supporting lymphocyte homing and proliferation. Immunity. 9, 669–676 (1998).

26. Kennedy, M.K. et al. Reversible defects in natural killer and memory CD8 T cell lineages in interleukin 15-deficient mice. J Exp Med. 191, 771–780 (2000).

27. Becker, T.C. et al. Interleukin 15 is required for proliferative renewal of virus-specific memory CD8 T cells. J Exp Med. 195, 1541–1548 (2002).

28. Goldrath, A.W. et al. Cytokine requirements for acute and Basal homeostatic proliferation of naive and memory CD8+ T cells. J Exp Med. 195, 1515–1522 (2002).

29. Geginat, J., Lanzavecchia, A. & Sallusto, F. Proliferation and differentiation potential of human CD8+ memory T-cell subsets in response to antigen or homeostatic cytokines. Blood. 101, 4260–4266 (2003).

30. Correia, M.P. et al. Hepatocytes and IL-15: a favorable microenvironment for T cell survival and CD8+ T cell differentiation. J Immunol. 182, 6149–6159 (2009).

31. Correia, M.P., Costa, A.V., Uhrberg, M., Cardoso, E.M. & Arosa, F.A. IL-15 induces CD8+ T cells to acquire functional NK receptors capable of modulating cytotoxicity and cytokine secretion. Immunobiology. 216, 604–612 (2011).

32. Liu, K., Catalfamo, M., Li, Y., Henkart, P.A. & Weng, N.P. IL-15 mimics T cell receptor crosslinking in the induction of cellular proliferation, gene expression, and cytotoxicity in CD8+ memory T cells. Proc Natl Acad Sci U S A. 99, 6192–6197 (2002).

33. Balin, S.J. et al. Human antimicrobial cytotoxic T lymphocytes, defined by NK receptors and antimicrobial proteins, kill intracellular bacteria. Sci Immunol. 3 (2018).

34. Seok, J. et al. A virtual memory CD8(+) T cell-originated subset causes alopecia areata through innate-like cytotoxicity. Nat Immunol. 24, 1308–1317 (2023).

35. Jabri, B. & Abadie, V. IL-15 functions as a danger signal to regulate tissue-resident T cells and tissue destruction. Nat Rev Immunol. 15, 771–783 (2015).

36. Ma, S., Caligiuri, M.A. & Yu, J. Harnessing IL-15 signaling to potentiate NK cell-mediated cancer immunotherapy. Trends Immunol. 43, 833–847 (2022).

37. Mishra, A., Sullivan, L. & Caligiuri, M.A. Molecular pathways: interleukin-15 signaling in health and in cancer. Clin Cancer Res. 20, 2044–2050 (2014).

38. Smith-Garvin, J.E., Koretzky, G.A. & Jordan, M.S. T cell activation. Annu Rev Immunol. 27, 591–619 (2009).

39. Roberts, A.I. et al. NKG2D receptors induced by IL-15 costimulate CD28-negative effector CTL in the tissue microenvironment. J Immunol. 167, 5527–5530 (2001).

40. Deshpande, P. et al. IL-7- and IL-15-mediated TCR sensitization enables T cell responses to self-antigens. J Immunol. 190, 1416–1423 (2013).

41. Richer, M.J. et al. Inflammatory IL-15 is required for optimal memory T cell responses. J Clin Invest. 125, 3477–3490 (2015).

42. Vaeth, M., Kahlfuss, S. & Feske, S. CRAC Channels and Calcium Signaling in T Cell-Mediated Immunity. Trends Immunol. 41, 878–901 (2020).

43. Clipstone, N.A. & Crabtree, G.R. Identification of calcineurin as a key signalling enzyme in T-lymphocyte activation. Nature. 357, 695–697 (1992).

44. Chen, L. et al. Structure of the DNA-binding domains from NFAT, Fos and Jun bound specifically to DNA. Nature. 392, 42–48 (1998).

45. Macián, F., García-Rodríguez, C. & Rao, A. Gene expression elicited by NFAT in the presence or absence of cooperative recruitment of Fos and Jun. EMBO J. 19, 4783–4795 (2000).

46. Eferl, R & Wagner, E.F. AP-1: a double-edged sword in tumorigenesis. Nature Reviews Cancer. 3, 859–868 (2003).

47. Marusina, A.I et al. Regulation of human DAP10 gene expression in NK and T cells by Ap-1 transcription factors. J Immunol. 180, 409–417 (2008).

48. Azzi, J.R., Sayegh, M.H. & Mallat, S.G. Calcineurin inhibitors: 40 years later, can’t live without. J Immunol. 191, 5785–5791 (2013).

49. Lee, H.G. et al. Pathogenic function of bystander-activated memory-like CD4(+) T cells in autoimmune encephalomyelitis. Nat Commun. 10, 709 (2019).

50. Langmead, B. & Salzberg, S.L. Fast gapped-read alignment with Bowtie 2. Nat Methods. 9, 357–359 (2012).

51. Love, M.I., Huber, W. & Anders, S. Moderated estimation of fold change and dispersion for RNA-seq data with DESeq2. Genome Biol. 15, 550 (2014).

52. Ritchie, M.E. et al. limma powers differential expression analyses for RNA-sequencing and microarray studies. Nucleic Acids Res. 43, e47 (2015).

53. Giles, J.R. et al. Human epigenetic and transcriptional T cell differentiation atlas for identifying functional T cell-specific enhancers. Immunity. 55, 557–574 (2022).

54. Yukawa, M. et al. AP-1 activity induced by co-stimulation is required for chromatin opening during T cell activation. J Exp Med. 217, e20182009 (2019).

55. Stuart, T. et al. Comprehensive Integration of Single-Cell Data. Cell. 177, 1888–1902 e1821 (2019).

56. Durinck, S., Spellman, P.T., Birney, E. & Huber, W. Mapping identifiers for the integration of genomic datasets with the R/Bioconductor package biomaRt. Nat Protoc. 4, 1184–1191 (2009).

