## Supplemental information for "TCR signaling via NFATc1 constrains IL-15-induced NK-like activation of human memory CD8^+^ T cells"

Contents

10 supplemental figures

4 supplemental tables

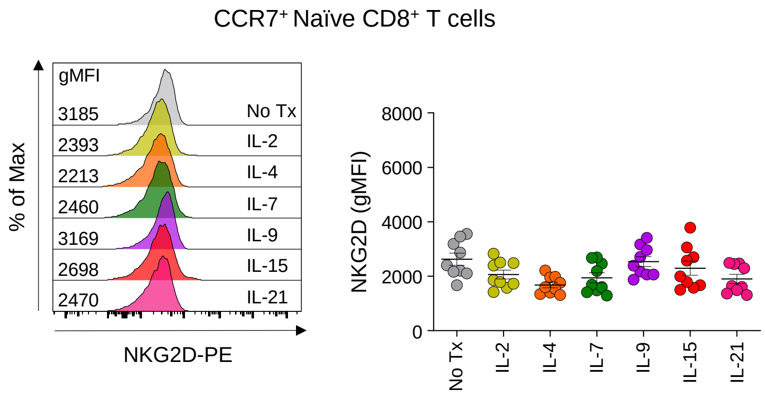


**Figure S1. NKG2D expression on naïve CD8⁺ T cells following stimulation with common gamma chain cytokines.** Peripheral blood mononuclear cells (PBMCs) from healthy donors (*n* = 9) were stimulated for 48 hours with IL-2 (10 ng/ml), IL-4 (10 ng/ml), IL-7 (10 ng/ml), IL-9 (10 ng/ml), IL-15 (10 ng/ml) or IL-21 (50 ng/ml). Representative stacked flow cytometry histograms and cumulative data presenting NKG2D expression on naïve ($CCR7^+$) $CD8^+$ T cells from healthy donor PBMCs following stimulation with indicated cytokines. Error bars represent mean ± SD.

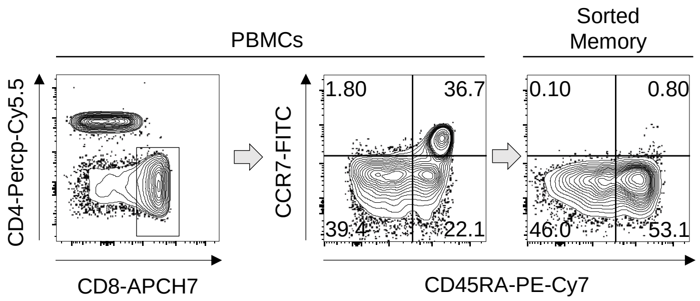


**Figure S2. Purity of sorted memory CD8$^+$ T cells.** CD8$^+$ T cells were negatively isolated from healthy donor PBMCs. The sorted CD8$^+$ T cells were stained with FITC-conjugated anti-CCR7 antibodies and anti-FITC microbeads to negatively isolate CCR7$^-$CD8$^+$ T cells. Representative flow cytometry plots illustrate the expression of CCR7 and CD45RA in the gate of CD8$^+$ T cells from PBMCs and sorted cells.

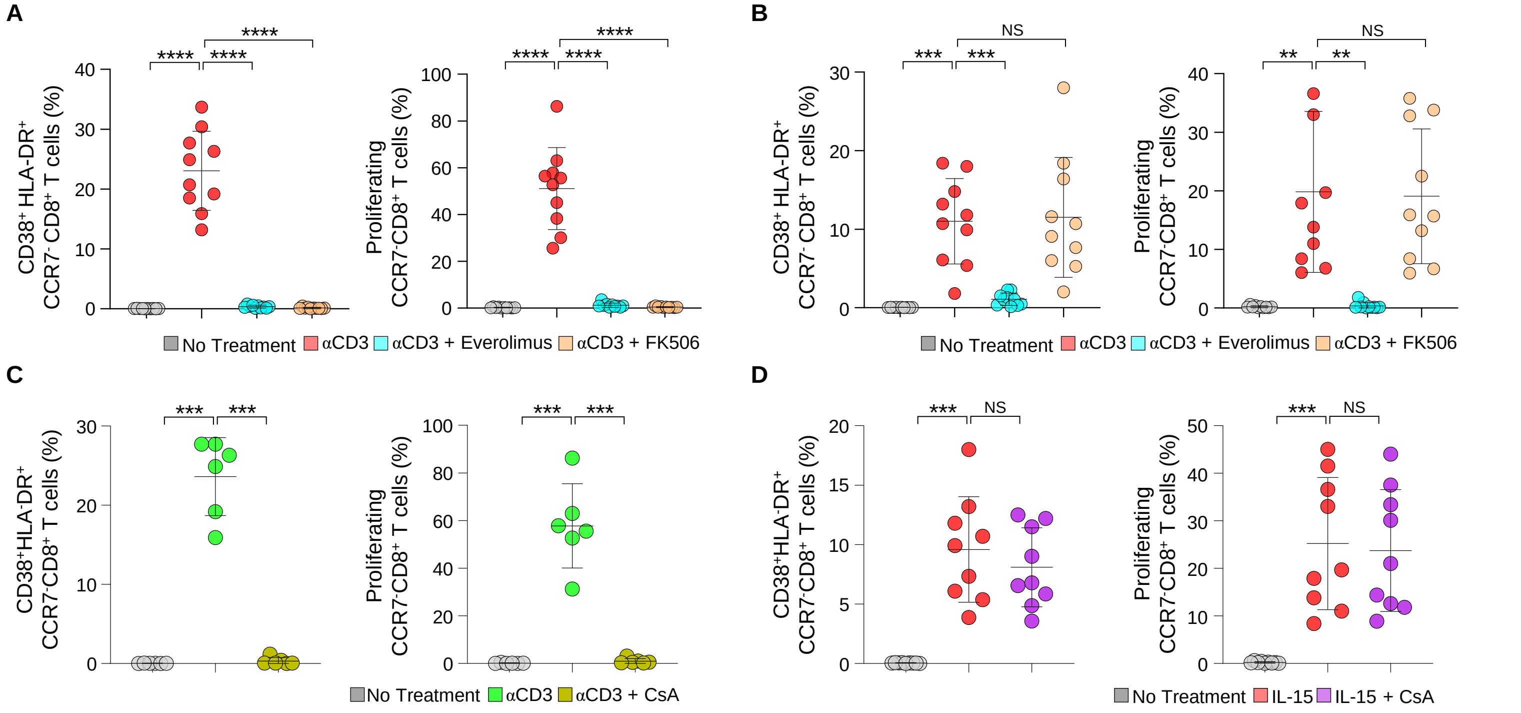


**Figure S3. Influence of everolimus, FK506, and CsA on the IL-15-induced activation and proliferation of memory CD8$^{+}$ T cells.** (**A and B**) Sorted CCR7$^{-}$ memory CD8$^{+}$ T cells from healthy donors (*n* = 10) were stained with CellTrace Violet and pre-treated with everolimus (10 μM) or FK506 (10 ng/ml) for 1 hour, and then cultured with anti-CD3 (coated, 1 μg/ml) or IL-15 (10 ng/ml) for 96 hours. Cumulative data present CD38 and HLA-DR activation markers and proliferation of CCR7$^{-}$ memory CD8$^{+}$ T cells after stimulation with (A) anti-CD3 or (B) IL-15. (**C and D**) Sorted CCR7$^{-}$ memory CD8$^{+}$ T cells from healthy donors (*n* = 6) were stained with CellTrace Violet and pre-treated with CsA (100 ng/ml) for 1 hour, and then cultured with anti-CD3 (coated, 1 μg/ml) or IL-15 (10 ng/ml) for 96 hours. Cumulative data show CD38 and HLA-DR activation markers and proliferation of CCR7$^{-}$ memory CD8$^{+}$ T cells after stimulation with (C) anti-CD3 (*n* = 6) or (D) IL15 (*n* = 6). Error bars represent mean ± SD. Statistical analysis was performed using the paired Student's t test. NS, not significant, $^{**}P < 0.01$, $^{***}P < 0.001$, $^{****}P < 0.0001$.

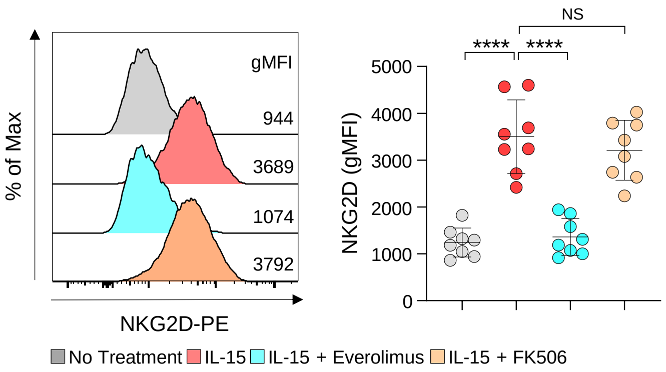


**Figure S4. Influence of everolimus and FK506 on the IL-15-induced upregulation of NKG2D on memory CD8⁺ T cells.** Sorted CCR7⁻ memory CD8⁺ T cells from healthy donors (*n* = 8) were pre-treated with everolimus (10 μM) or FK506 (10 ng/ml) for 1 hour, and then cultured with IL-15 (10 ng/ml) for 48 hours. Representative stacked flow cytometry histograms and cumulative data present NKG2D expression on CCR7⁻ memory CD8⁺ T cells. Error bars represent mean ± SD. Statistical analysis was performed using the paired Student's t test. NS, not significant, ****$P$ < 0.0001.

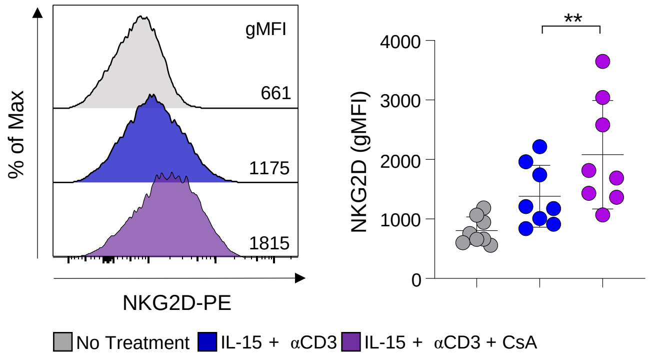


**Figure S5. Paradoxical effect of CsA on NKG2D expression on memory CD8⁺ T cells.** Sorted CCR7⁻ memory CD8⁺ T cells from healthy donors (*n* = 8) were pretreated with CsA (100 ng/ml) for 1 hour, and then stimulated with IL-15 (10 ng/ml) and anti-CD3 (coated, 1 μg/ml) for 48 hours. Representative stacked flow cytometry histograms and cumulative data present NKG2D expression on CCR7⁻ memory CD8⁺ T cells. Error bars represent mean ± SD. Statistical analysis was performed using the paired Student's t test. $^{**}P < 0.01$.

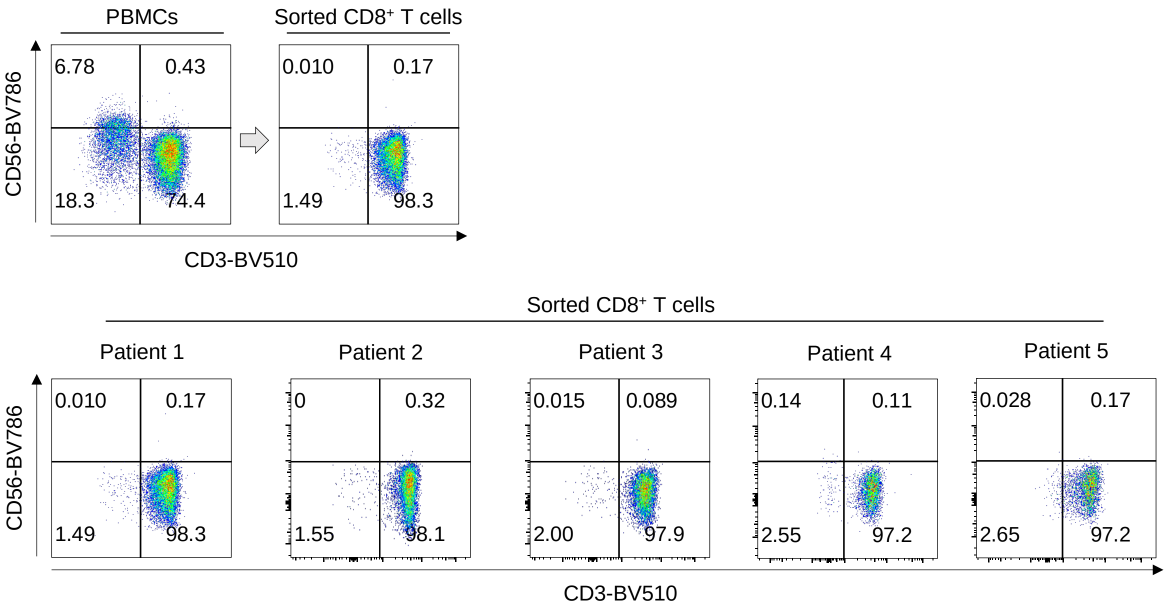


**Figure S6. Expression of CD56 among sorted CD8$^+$ T cells from patients with acute HAV infection.** CD8$^+$ T cells were negatively isolated from PBMCs from patients with acute HAV infection. Representative flow cytometry plots illustrate the expression of CD56 and CD3 in the gate of CD8$^+$ T cells from sorted cells of individual patients with acute HAV infection.

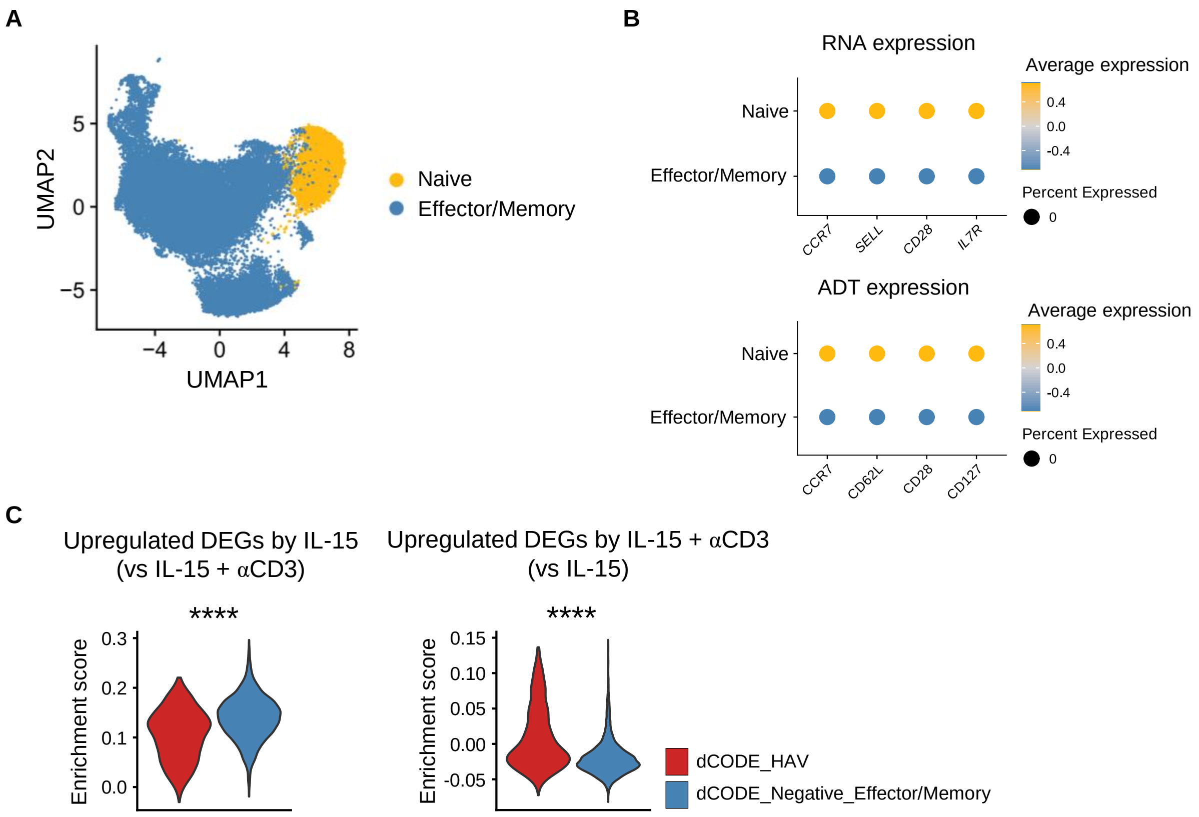


**Figure S7. Enrichment of differentially expressed genes upregulated by IL-15 or IL-15 plus anti-CD3 in dCODE⁻ effector/memory and HAV-specific CD8⁺ T cells from patients with acute HAV infection.** (**A**) UMAP projections of naïve and effector/memory dCODE dextramer-negative CD8⁺ T cells from patients with acute HAV infection. (**B**) Dot plots showing the normalized expression of marker genes and ADTs in naïve and effector/memory dCODE dextramer-negative CD8⁺ T cells. (**C**) Violin plots showing enrichment score for DEGs upregulated by stimulation with IL-15 compared to IL-15 plus anti-CD3 in memory CD8⁺ T cells in dCODE dextramer-negative effector/memory CD8⁺ T cells and dCODE dextramer-positive HAV-specific CD8⁺ T cells from patients with acute HAV infection in scRNA-seq analysis. Statistical analysis was performed using the Mann-Whitney U-test. ****$P < 0.0001$.

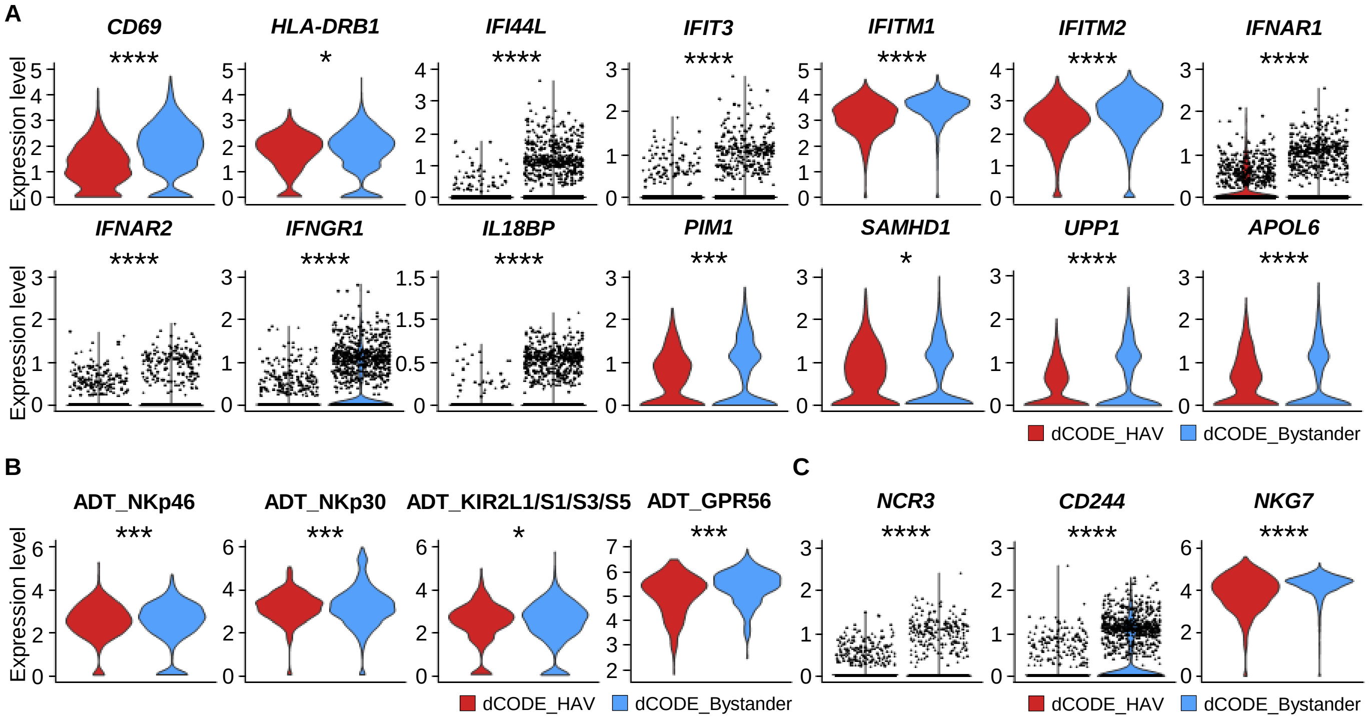


**Figure S8. Expression of genes and ADT differentially upregulated in bystander CD8⁺ T cells compared to HAV-specific CD8⁺ T cells from patients with acute HAV infection.** (**A**) Violin plots showing normalized expression of genes that are associated with IFN response in HAV-specific versus bystander CD8⁺ T cells from patients with acute HAV infection. (**B** and **C**) Violin plots showing (B) ADT expression of NKp46, NKp30, KIR2L1/S1/S3/S5, and GPR56 and (C) normalized gene expression of *NCR3*, *CD244*, and *NKG7* in HAV-specific versus bystander CD8⁺ T cells from patients with acute HAV infection. Statistical analysis was performed using the Mann-Whitney U-test $*P < 0.05$, $***P < 0.001$, $****P < 0.0001$.

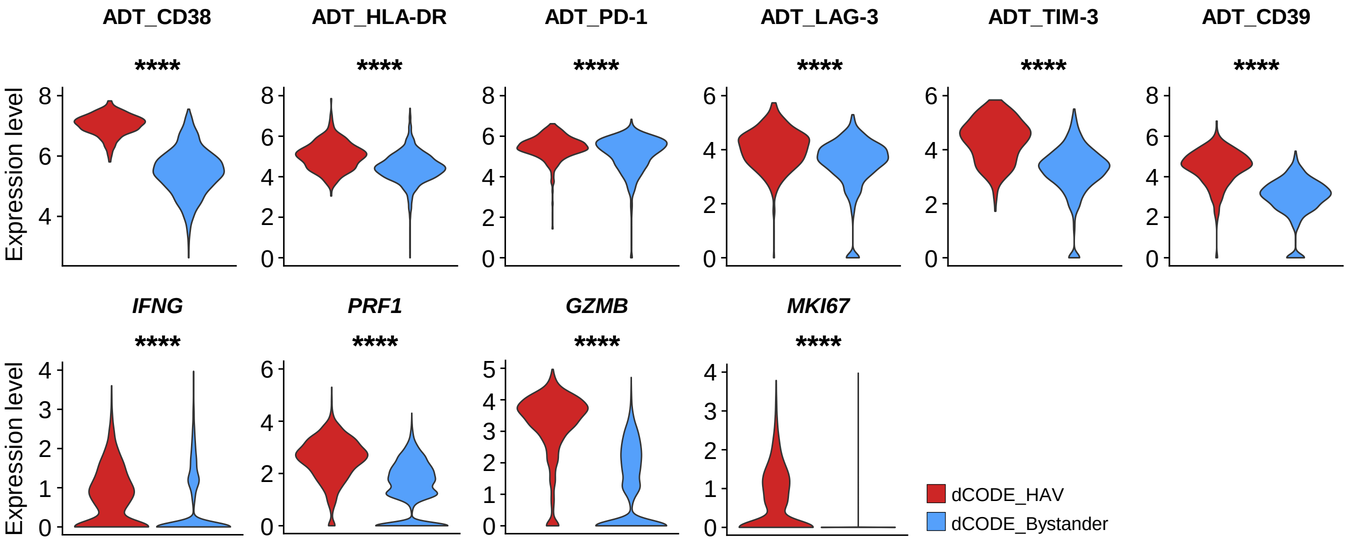


**Figure S9. Expression of ADT and genes differentially upregulated in HAV-specific CD8⁺ T cells compared to bystander CD8⁺ T cells from patients with acute HAV infection.** Violin plots showing ADT expression of CD38, HLA-DR, PD-1, LAG-3, TIM-3, and CD39, and normalized gene expression of *IFNG*, *PRF1, GZMB,* and *MKI67* between HAV-specific and bystander dCODE dextramer-positive $CD8^+$ T cells from patients with acute HAV infection and healthy donors. Statistical analysis was performed using the Mann-Whitney U-test. ****$P < 0.0001$.

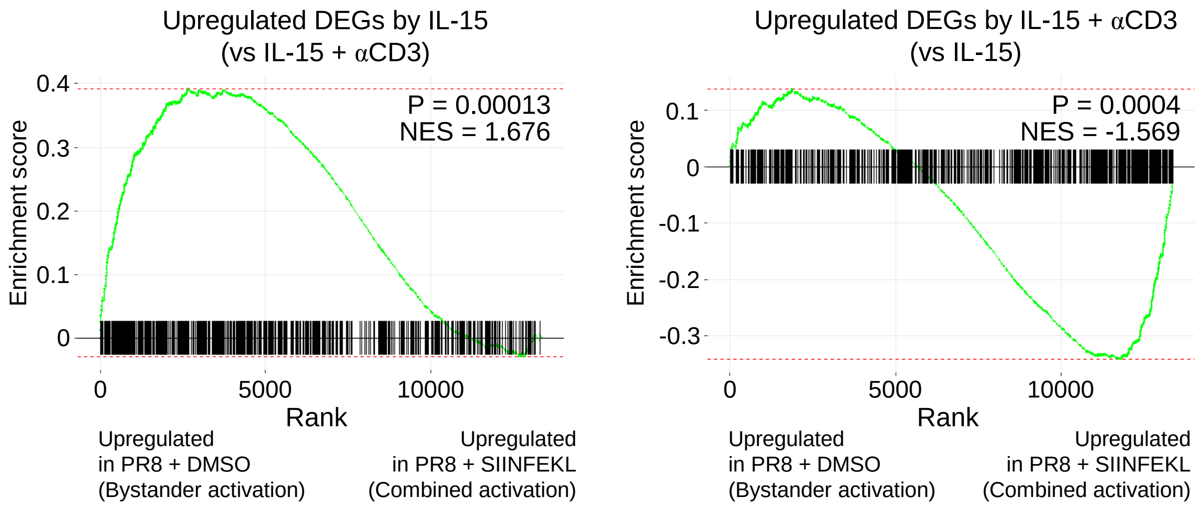


**Figure S10. Enrichment of gene sets induced by stimulation with IL-15 or IL-15 plus αCD3 in OT-1 T cells isolated from mice infected with influenza virus, with or without SIINFEKL.** GSEA of upregulated DEGs obtained from bulk RNA-seq of memory $CD8^+$ T cells stimulated with IL-15 or IL-15 plus anti-CD3 using the previously reported transcriptome of OT-1 T cells from mice with PR8 IAV infection, with or without SIINFEKL injection.[20]

### Supplemental Tables

| **Table S1. List of differentially expressed genes in memory CD8⁺ T cells clustered by upregulation patterns following stimulation with IL-15, anti-CD3 or the combination of both.** | |
|---|---|
| **Cluster 1** | ERRFI1, CASZ1, UQCRHL, FAM131C, TIE1, TCTEX1D4, MCOLN3, EXTL2, ATP5F1, CTTNBP2NL, DENND2C, HIST2H2BE, ENSA, NTRK1, FCRL3, SLC19A2, TNFSF4, NCF2, RGL1, DUSP10, GALNT2, TFB2M, IL15RA, ARL5B, C10orf25, UBE2D1, SLC25A16, SRGN, FAM149B1, HECTD2, UBTD1, INPP5F, ECHS1, PAOX, DENND5A, MDK, VPS37C, TKFC, ROM1, TIGD3, UCP2, NDUFC2-KCTD14, MTMR2, DIXDC1, PHLDB1, NLRX1, ARHGEF12, PUS3, SLC2A13, SLC48A1, RARG, SMUG1, SPRYD4, CYP27B1, NAP1L1, MAPKAPK5, ANKLE2, ZNF891, RGCC, CCDC122, GTF2F2, TBC1D4, ATP11A, FOS, ADCK1, DPH6, PAK6, LCMT2, SHF, GABPB1, RSL24D1, C2CD4A, BCL2A1, LRRC28, BRICD5, IL21R, MYLPF, RRAD, TANGO6, CYB5B, ZNRF1, SLC43A2, CYB5D2, CAMTA2, TMEM88, UTP6, CCL13, COA3, MAP3K14, SNX11, SEC14L1, RAB40B, PSTPIP2, DYM, HMSD, GPX4, IZUMO4, GADD45B, C19orf44, RBM42, NFKBID, SPINT2, PLA2G4C, ETFB, ZNF628, ZNF324B, FKBP1B, MEMO1, FANCL, FAM161A, GPAT2, DUSP2, COX5B, BCL2L11, TANK, METTL8, NCKAP1, STAT1, NDUFB3, SPAG16, UBE2F, LOC388780, RBBP9, TP53INP2, CD40, KCNQ2, RWDD2B, HMGN1, SEC14L2, SLC35E4, TNFRSF13C, PRRT3, RFTN1, LZTFL1, PARP3, PSMD6, CLDND1, GPR15, NDUFB4, MGLL, NDUFB5, EHHADH, PIGX, MYL5, MFSD7, MFSD10, EVC, SMIM20, CXCL9, RASGEF1B, LSM6, OTULIN, ITGA1, ERCC8, TAF9, UTP15, SRFBP1, TSPAN17, TMEM170B, TPMT, HIST1H2BF, UBD, HSD17B8, MAPK13, FOXP4, RRP36, GSTA4, COX7A2, BACH2, RTN4IP1, FAM229B, PLAGL1, SUMO4, ZC3H12D, ULBP2, PPP1R14C, PLEKHG1, SFT2D1, FGFR1OP, MRPL32, GBAS, CAV1, ST7, LRRC4, RDH10, TMEM70, PTP4A3, GRINA, C9orf64, ASMTL, NHS, CXorf21, CHST7, FTSJ1, SLC35A2, MAGED1, CXorf57, PSMD10, NKRF, TMEM185A, CXorf40B |
| **Cluster 2** | B3GALT6, LYPLA2, AUNIP, HMGN2, TXLNA, TMEM234, MANEAL, PPT1, PPIH, EIF2B3, ZSWIM5, FAF1, GPX7, TYW3, RABGGTB, SETSIP, KCNC4, ANKRD34A, RBM8A, HIST2H2AA4, SF3B4, PIP5K1A, PSMD4, MRPL9, OAZ3, UBAP2L, MUC1, FAM189B, UBQLN4, IQGAP3, ILDR2, MPZL1, C1orf53, TMCC2, IL10, TMEM206, PARP1, HIST3H2A, NTPCR, KMO, GDI2, RBM17, ATP5C1, PHYH, ACBD7, HACD1, PDSS1, BMS1, VSTM4, SLC16A9, COMTD1, KIF20B, RPP30, MYOF, NDUFB8, BCCIP, DHX32, ZNF511, HRAS, PIDD1, CD151, PRMT3, CAPRIN1, TMX2, MRPL16, VWCE, DDB1, C11orf84, MACROD1, CCDC85B, LRP5, PPME1, CLNS1A, DCUN1D5, POU2AF1, PPP2R1B, PIH1D2, OAF, NRGN, RPUSD4, ST3GAL4, GLB1L2, NOP2, ETV6, DNM1L, AAAS, METTL7B, ITGA7, METAP2, IKBIP, UHRF1BP1L, CHPT1, MMAB, MVK, PTPN11, DDX54, GCN1, MLEC, CAMKK2, PITPNM2, DHX37, PGAM5, EBPL, MZT1, GPR180, CCNB1IP1, DHRS4, FANCM, L2HGDH, PSMA3, SYNJ2BP-COX16, COX16, FLVCR2, ANGEL1, POMT2, EFCAB11, TTC7B, ITPK1, GLRX5, CINP, CKB, C14orf2, RPUSD2, EHD4, HAUS2, HYPK, EIF3J, CEP152, MNS1, FAM96A, PIF1, CIB2, CHRNA3, CTSH, MRPL46, MESP1, FURIN, FES, LYSMD4, NPRL3, NME4, RPUSD1, UBE2I, BAIAP3, TSR3, TELO2, TRAF7, RNPS1, HCFC1R1, CORO7-PAM16, PAM16, EEF2KMT, EEF2K, TBC1D10B, ZNF768, CTF1, FBXL19, ITGAX, ARMC5, C16orf87, HEATR3, CCL17, NAE1, FAM96B, E2F4, SLC9A5, ACD, NOB1, DDX19A, DHODH, USP10, SPATA33, GAS8, CLUH, EMC6, GSG2, PELP1, KIF1C, ZNF232, SLC16A13, NAA38, SREBF1, RAB34, PSMD11, ACACA, NT5C3B, ACLY, KAT2A, KCNH4, PSMC3IP, DBF4B, EFTUD2, UTP18, SRSF1, MKS1, MRC2, FTSJ3, PSMC5, TEX2, PSMD12, GALK1, SRP68, TBC1D16, BAIAP2, CCDC137, OGFOD3, CLUL1, TWSG1, SPIRE1, TMEM241, ME2, LMAN1, PHLPP1, C19orf24, GAMT, ADAT3, GNA11, MFSD12, FSD1, STAP2, PRR22, PNPLA6, FDX1L, ILF3, CARM1, DOCK6, TNPO2, ASNA1, SYCE2, ILVBL, RAB8A, GTPBP3, COLGALT1, RHPN2, KCTD15, TMEM147, C19orf47, ZNF576, PVR, AP2S1, NAPSA, ZSCAN5A, ADGRF3, SIX3, PRKCE, POLR1A, FAHD2A, ARID5A, ANKRD39, FAHD2B, IL1R2, TTL, IL1RN, CBWD2, RALB, R3HDM1, PKP4, CERS6, NUP35, MYO1B, DNAH7, MARS2, SPATS2L, FZD5, ARMC9, PTMA, HDLBP, THAP4, IDH3B, SLC4A11, SMOX, TRMT6, NDUFAF5, SNX5, CTNNBL1, TTI1, PKIG, PTGIS, BCAS4, CBR1, SIK1, RRP1B, PWP2, GNB1L, ASPHD2, AP1B1, THOC5, OSBP2, RBFOX2, TXN2, MPST, GCAT, RPS19BP1, PANX2, MAPK12, RAD18, FANCD2OS, SEC13, TSEN2, CHCHD4, SATB1, ANO10, ZNF35, ELP6, CELSR3, NCKIPSD, HEMK1, FRMD4B, FILIP1L, TRMT10C, TOPBP1, P2RY13, GFM1, USP13, PARL, XXYLT1, FBXO45, NCBP2, TACC3, AFAP1, ACOX3, DHX15, PI4K2B, TBC1D1, RASL11B, HNRNPD, TRMT10A, RWDD4, CASP3, TRIO, SKP2, OXCT1, KIF2A, GTF2H2C_2, GTF2H2, WDR41, ATG10, CETN3, CCNI2, SHROOM1, DNAJC18, NDST1, LARP1, GEMIN5, BNIP1, CLTB, PRR7, DDX41, BPHL, PSMG4, HIST1H1A, HIST1H2BC, HIST1H1E, HIST1H1D, HIST1H2AG, HIST1H2BM, FLOT1, GTF2H4, PPT2, RPS10-NUDT3, FANCE, SRPK1, GLO1, C6orf132, MRPS10, RPL7L1, TMEM14A, KHDC1, SYNCRIP, NDUFAF4, COQ3, FAM184A, SOGA3, KIAA0408, OLIG3, SYTL3, PDCD2, PSMG3, ARL4A, AEBP1, VKORC1L1, BUD31, PTCD1, ATP5J2-PTCD1, SPDYE3, SRRT, LRWD1, PRKAR2B, CALU, TBXAS1, ZNF282, LRRC61, ABCF2, NOM1, FAM86B1, REEP4, WRN, POLB, PRKDC, NDUFAF6, NCALD, SLC25A32, NDUFB9, TOP1MT, PUF60, SLC39A4, CBWD1, CLTA, FXN, TJP2, HNRNPK, AUH, CENPP, OMD, ECM2, ZNF782, NANS, DNAJC25-GNG10, GNG10, NDUFA8, STXBP1, SLC27A4, URM1, MED27, WDR5, C9orf116, PMPCA, TRAF2, NPDC1, SSNA1, NRARP, MRPL41, DHRSX, RBBP7, EIF1AX, MAOA, ELK1, CCDC22, FOXP3, HUWE1, PHKA1, VMA21, NSDHL, NAA10, HCFC1, FUNDC2 |
| **Cluster 3** | SAMD11, NOC2L, AGRN, C1orf159, TNFRSF4, AURKAIP1, ATAD3B, PARK7, CTNNBIP1, APITD1-CORT, SDHB, RCC2, GPN2, NUDC, MECR, EIF3I, SFPQ, GNL2, C1orf109, YRDC, FHL3, NDUFS5, PABPC4, MFSD2A, YBX1, ATP6V0B, ERI3, TOE1, TESK2, MMACHC, PRDX1, TMEM69, UQCRH, COA7, MAGOH, TTC4, FGGY, IL12RB2, SERBP1, GADD45A, CTH, GNG5, MCOLN2, LRRC8B, EVI5, ALG14, VCAM1, WDR3, PPIAL4G, PPIAL4A, PPIAL4C, CHD1L, HIST2H2AB, PSMB4, TUFT1, ILF2, SLC39A1, CREB3L4, SLC50A1, MTX1, LAMTOR2, LMNA, PMF1, CCT3, HDGF, TAGLN2, PEA15, PFDN2, F5, C1orf112, ACBD6, GLRX2, SLC30A1, NENF, RRP15, PSEN2, URB2, FAM89A, SLC35F3, GNG4, GTPBP4, PITRM1, NUDT5, CDC123, HNRNPF, CISD1, TFAM, CCDC6, DNAJC12, DDX21, LRRC20, MRPS16, CHCHD1, ADK, VDAC2, NOC3L, ALDH18A1, DPCD, NOLC1, PSD, CUEDC2, PDCD11, MGMT, FUOM, PSMD13, PTDSS2, CHID1, IGF2, INS-IGF2, PGAP2, PRKCDBP, IPO7, PSMA1, NCR3LG1, RCN1, NAT10, ABTB2, APIP, PDHX, CD82, PACSIN3, ACP2, PTPMT1, CCDC86, PTGDR2, PRPF19, INCENP, LBHD1, C11orf98, WDR74, ARL2, FOSL1, YIF1A, NDUFV1, MRPL21, CTTN, DHCR7, FAM86C1, RNF121, CLPB, COA4, POLD3, DGAT2, NARS2, EED, BIRC3, SLC35F2, ZC3H12C, RDX, LAYN, FDXACB1, C11orf1, DLAT, TIMM8B, PTS, IFT46, SLC37A4, DPAGT1, CRTAM, GRAMD1B, ACRV1, FOXRED1, DYRK4, MRPL51, PTMS, USP5, ATN1, PHB2, YBX3, STRAP, LDHB, MRPS35, ZCRB1, HDAC7, LIMA1, LARP4, GRASP, NR4A1, C12orf10, TARBP2, HNRNPA1, CS, ATP5B, TSFM, TMEM5, IFNG, MRPL42, CEP83, TMPO, SLC25A3, UTP20, NUP37, SLC41A2, C12orf45, ALKBH2, GPN3, FAM216A, RNFT2, VSIG10, RPLP0, TRIAP1, COQ5, POP5, HNF1A, C12orf43, RILPL2, GTF2H3, PUS1, NOC4L, MRPL57, CENPJ, USP12, EXOSC8, NUFIP1, NUDT15, SPRYD7, UCHL3, IPO5, KDELC1, APEX1, HNRNPC, SUPT16H, PSMB5, ZFHX2, DHRS2, IPO4, CBLN3, DTD2, MBIP, C14orf166, TOMM20L, TIMM9, SNAPC1, FNTB, GPHN, EIF2S1, BATF, GSTZ1, AHSA1, SLIRP, VRK1, WDR25, HSP90AA1, SIVA1, NIPA2, ARHGAP11B, AVEN, NOP10, ITPKA, SORD, MAPK6, GTF2A2, TLN2, LACTB, STOML1, TSPAN3, PSMA4, CHRNA5, TMED3, HOMER2, AEN, RCCD1, SNRPA1, POLR3K, MRPL28, METRN, FAM173A, C16orf91, HN1L, MRPS34, TBL3, SYNGR3, NTHL1, MLST8, PGP, ECI1, PRSS21, DNASE1, TRAP1, TFAP4, GLIS2, RSL1D1, ITPRIPL2, KNOP1, ACSM3, ERI2, LOC81691, POLR3E, NSMCE1, EIF3CL, EIF3C, TUFM, SLX1A, BCKDK, NUDT21, MT1G, MT1H, NUP93, CCL22, CCDC102A, KATNB1, GOT2, RFWD3, TMEM231, CMC2, GCSH, HSBP1, EMC8, APRT, TRAPPC2L, GEMIN4, SRR, TSR1, MYBBP1A, SLC25A11, RPAIN, DHX33, TXNDC17, ELP5, CLDN7, WRAP53, SCO1, CENPV, NT5M, PEMT, TMEM97, POLDIP2, TLCD1, TRAF4, ANKRD13B, BLMH, ATAD5, TEFM, CCL1, RAD51D, RDM1, CCL4, CCL3L3, CCL3L1, CCL4L1, CCL4L2, DHRS11, MRM1, TADA2A, MRPL45, CCR7, STAT5A, AARSD1, PTGES3L-AARSD1, G6PC3, SLC25A39, GJC1, KPNB1, PNPO, CBX1, HOXB7, HOXB9, MRPL27, NME2, AKAP1, MRPS23, RAD51C, TACO1, CCDC47, FDXR, ATP5H, SLC25A19, GALR2, SPHK1, SRSF2, SYNGR2, AFMID, CBX2, DCXR, ENOSF1, TUBB6, SEH1L, FAM210A, CABLES1, SETBP1, ATP5A1, MBD2, SEC11C, CYB5A, PQLC1, TXNL4A, FSTL3, PTBP1, WDR18, DAZAP1, MBD3, DOT1L, SLC39A3, NCLN, SIRT6, EBI3, UHRF1, LONP1, NDUFA11, KHSRP, TNFSF9, TRIP10, PEX11G, TGFBR3L, ELAVL1, NDUFA7, HNRNPM, MRPL4, QTRT1, YIPF2, ZNF788, JUNB, PRDX2, GCDH, RAD23A, GADD45GIP1, TRMT1, HAUS8, NR2F6, ANKLE1, INSL3, DDX49, ZNF93, UQCRFS1, POP4, PDCD5, FAAP24, PDCD2L, IGFLR1, WDR62, POLR2I, MRPS12, FBL, SNRPA, KCNN4, FOSB, EML2, CALM3, PTGIR, GNG8, DACT3, GRWD1, PRMT1, NUP62, JOSD2, VSIG10L, LIM2, ZNF534, PRPF31, NAT14, ZNF584, TRIM28, TSSC1, CPSF3, E2F6, RHOB, SF3B6, CENPO, DNMT3A, SLC5A6, CAD, PPM1G, WDR43, LCLAT1, TTC27, CYP1B1, HNRNPLL, GALM, LRPPRC, CCT4, MDH1, PNO1, FAM136A, SPR, SFXN5, SMYD5, CCT7, MOB1A, HTRA2, LOXL3, MAT2A, GGCX, MRPL35, RPIA, STARD7, EIF5B, MRPS9, FHL2, ACOXL, POLR1B, POLR2D, DARS, NR4A2, GPD2, METTL5, METAP1D, OLA1, MTX2, FKBP7, WDR75, STRADB, NOP58, CTLA4, ICOS, EEF1B2, ATIC, XRCC5, BCS1L, TTLL4, DNPEP, GMPPA, TMEM198, CCL20, PSMD1, NCL, COL6A3, RAMP1, MRPS26, ITPA, MCM8, SNRPB2, DTD1, PLAGL2, POFUT1, MAP1LC3A, GSS, EIF6, PABPC1L, TOMM34, CSE1L, PFDN4, PSMA7, CABLES2, BTG3, MRPL39, ATP5J, CCT8, SOD1, MIS18A, URB1, GART, RCAN1, ETS2, TMPRSS3, WDR4, U2AF1, MRPL40, SMARCB1, DRG1, RTCB, HMOX1, MB, TST, TOMM22, RANGAP1, PHF5A, XRCC6, SREBF2, NAGA, RRP7A, MCAT, TTLL12, SAMM50, PIM3, NCAPH2, SCO2, LMCD1, OGG1, RPUSD3, JAGN1, CMTM6, XIRP1, RPSA, EXOSC7, LARS2, CCDC51, TREX1, UQCRC1, GPX1, TRAIP, RRP9, ACY1, SMIM4, GNL3, TKT, KBTBD8, RYBP, CPOX, ALCAM, CD200, CD80, CD86, CCDC58, FAM162A, PDIA5, UMPS, CHCHD6, RUVBL1, EEFSEC, ACAD9, MRPL3, PCCB, WWTR1, SIAH2, NMD3, GOLIM4, ACTL6A, MRPL47, MAP6D1, ALG3, PSMD2, EIF4G1, POLR2H, MAGEF1, TRA2B, BDH1, IQCG, LETM1, TNIP2, NOP14, JAKMIP1, GRPEL1, CD38, QDPR, LAP3, PACRGL, RBPJ, SRD5A3, PPAT, HOPX, CXCL13, ENOPH1, EIF4E, NFKB1, SLC9B2, GSTCD, HADH, GAR1, C4orf32, TNIP3, EXOSC9, LARP1B, MGST2, ABCE1, GATB, NAF1, ANP32C, SNX25, AHRR, MRPL36, NDUFS6, SUB1, BRIX1, NDUFAF2, AK6, PTCD2, ELL2, WDR36, HSD17B4, HINT1, IL3, CSF2, PDLIM4, IL5, IL4, KIF3A, GDF9, HSPA4, IL9, ETF1, HSPA9, MRPL22, TIMD4, FNDC9, THG1L, NUDCD2, RARS, SPDL1, NPM1, THOC3, ARL10, RMND5B, GMDS, RPP40, RIOK1, EEF1E1, |

| | |
|---|---|
| | HIVEP1, NOL7, HIST1H2AB, HIST1H1C, HIST1H4H, HIST1H2AM, MRPS18B, CSNK2B, LSM2, WDR46, PFDN6, HMGA1, SRSF3, TMEM217, TBC1D22B, TOMM6, POLR1C, XPO5, NFKBIE, CD109, UBE3D, ORC3, RRAGD, GTF3C6, HDAC2, ENPP1, MYB, AHI1, LTV1, SF3B5, TIAM2, ZDHHC14, SNX9, ACAT2, TCP1, MRPL18, PSMB1, PDGFA, DNAAF5, C7orf50, EIF3B, AIMP2, MALSU1, CYCS, HNRNPA2B1, CBX3, MRPS24, URGCP-MRPS24, TBRG4, MDH2, HSPB1, YWHAG, DBF4, SRI, CYP51A1, TFR2, TRIP6, DLD, MET, FAM3C, SND1, IRF5, TSPAN33, MEST, COPG2, HIPK2, MRPS33, SSBP1, CDK5, PINX1, POLR3D, EGR3, UBE2V2, LYPLA1, PLAG1, MSC, MRPS28, MTERF3, POP1, STK3, FBXO43, POLR2K, DCAF13, WDYHV1, TRIB1, TIGD5, PYCRL, CYC1, HGH1, TMEM249, FBXL6, TONSL, C8orf33, CD274, PLAA, DNAJA1, NOL6, VCP, STOML2, CD72, HINT2, GRHPR, POLR1E, EXOSC3, CBWD6, CBWD5, NINJ1, ANP32B, SEC61B, NR4A3, TEX10, MRPL50, PRPF4, TRAF1, MRRF, PSMB7, NR6A1, RABEPK, DPM2, TRUB2, COQ4, ODF2, SET, NUP188, EXOSC2, DDX31, MRPS2, CARD9, PHPT1, ZMYND19, PNPLA4, PRPS2, PDHA1, SMS, GK, SLC9A7, SLC38A5, SUV39H1, TIMM17B, PIM2, PPP1R3F, APEX2, LAS1L, CENPI, GLA, PRPS1, ACSL4, MCTS1, GLUD2, UTP14A, AIFM1, CXorf40A, MAMLD1, DKC1, NLGN4Y, EIF1AY |
| **Cluster 4** | ISG15, AGTRAP, TNFRSF1B, IFI6, PTAFR, KIAA1522, CITED4, IFI44L, GBP1, CDC7, RHOC, S100A11, ETV3L, ETV3, FASLG, SMYD2, NLRP3, AKR1C3, VIM, MAP3K8, IFIT3, PDLIM1, ARHGAP19, GOLGA7B, IFITM2, IFITM1, IFITM3, STX3, CFL1, CDK2AP2, IL18BP, TPBGL, CTSC, HYLS1, KCNA6, SLC2A14, CD69, PRPF40B, GALNT6, SOAT2, MYL6, NDUFA12, TXNRD1, ARPC3, LCP1, CLN5, NRL, GNG2, PTGER2, SYNE3, JAG2, NUDT14, CRIP1, GOLGA8J, GOLGA8R, GOLGA8O, SPRED1, PDIA3, C15orf48, MYO1E, ANXA2, RAB8B, LCTL, CYP1A1, C15orf39, PTPN9, IDH2, TNFRSF12A, IL32, CORO1A, BCL7C, CKLF-CMTM1, CBFB, NOL3, SMPD3, MTSS1L, COTL1, ZNF469, P2RX5, SPNS3, PFN1, MAPK7, MFAP4, ALDOC, CDK5R1, TNS4, IFI35, CALCOCO2, ABI3, CD79B, SLC16A6, WIPI1, TMEM104, RNF157, UBALD2, ACTG1, MYL12A, ZNF519, TICAM1, TNFSF14, ZNF699, ZNF823, HSH2D, TSHZ3, HCST, CEACAM1, VASP, NUCB1, FLT3LG, KIR2DL3, KIR2DL1, KIR2DL4, KIR2DS4, KIR3DL2, ZNF460, ZNF132, CMPK2, RSAD2, SPRED2, IL18R1, IL18RAP, ARL6IP6, SSFA2, PGAP1, TUBA4A, RASSF2, CD93, COMMD7, CEBPB, ADAMTS1, TRPM2, RGL4, XBP1, APOL6, LGALS1, MAFF, SMC1B, ADM2, COLQ, GLB1, POMGNT2, CCR5, PFKFB4, NFKBIZ, HEG1, PLXNA1, GYG1, FNDC3B, TNFSF10, IL1RAP, SLC51A, WFS1, COMMD8, MTHFD2L, FAM47E-STBD1, GPRIN3, HPGDS, ELOVL6, SETD7, HPGD, ITGA2, GZMA, CYSTM1, PFDN1, PDGFRB, CD74, ZNF300, NEURL1B, NCR3, DDAH2, HLA-DRA, HLA-DRB5, HLA-DRB1, CPNE5, PIM1, TREML2, CCND3, IL17F, FAM26F, TMEM200A, RPS6KA2, LFNG, ACTB, FAM126A, UPP1, PHTF2, ARPC1B, PTPRN2, DMTN, DPYSL2, GSR, MYBL1, C8orf88, SNTB1, DERL1, TMEM65, LY6E, CNTLN, DCTN3, SIT1, NTRK2, AKAP2, BSPRY, LOC100505478, TNFSF8, AGPAT2, UAP1L1, NSMF, LANCL3, FAM104B, RPA4, BEX5, NXT2, HTATSF1 |
| **Cluster 5** | DHRS3, CASP9, MINOS1-NBL1, IQCC, PLK3, BTF3L4, NRAS, TSEN15, UEVLD, CCDC15, DERA, GGACT, SPTSSA, CES4A, SAMD14, B4GALT6, MPND, CEBPG, LAIR2, TTC7A, VRK2, DENND6B, TIGIT, LAMP3, GNPDA2, COPS4, BASP1, HLA-DQA2, VLDLR, PLGRKT, ZXDA, F8 |
| **Cluster 6** | FBXO6, SCMH1, RIMKLA, PRMT6, BCL9, IDI1, PFKFB3, NEURL1, ASRGL1, FIBP, CNIH2, PC, RELT, HYOU1, STYK1, EMP1, RHEBL1, CBX5, GTSF1, STAC3, POC1B, POC1B-GALNT4, GALNT4, CIT, RAB15, DPF3, TRAF3, COMMD4, ABHD17C, LYRM1, KIAA0895L, PSMB10, TUBB3, MYO1C, RILP, FAM64A, LGALS9, ITGA3, SKA2, TRIM47, CHMP6, RNF165, TNFRSF11A, ZNF257, YIF1B, LIPE, PLAUR, DHDH, NLRP2, KLF11, TPRKB, CAPG, CHST10, CCDC74B, CYTIP, DPP4, IGFBP2, STK16, PER2, GPC1, TRIB3, SIRPG, TOX2, EEF1A2, SUMO3, CECR5, USP18, GGT1, CRELD2, XYLB, CCR1, CCR2, NICN1, MON1A, C3orf18, EOGT, GBE1, EAF2, C4orf48, ABCG2, SLC7A11, SCOC, P4HA2, MZB1, HBEGF, F12, PDLIM7, TMED9, FARS2, NEDD9, LTB, CLIC1, HLA-DQB1, RCAN2, RAB23, ULBP1, DBNL, SDHAF3, ARHGEF5, RNF32, MFHAS1, LZTS1, NECAB1, LAPTM4B, JRK, GPAA1, C9orf66, IPPK, PHF19, FAM129B, SURF4, TOR4A, EBP, ZC4H2, MORC4, SLC25A43, C1GALT1C1, CD40LG |
| **Cluster 7** | TNFRSF18, ATAD3A, RNF207, ICMT, ACOT7, TNFRSF9, ENO1, SLC25A33, TMEM201, PGD, SRM, MAD2L2, TNFRSF8, AGMAT, ALDH4A1, MRTO4, NBL1, E2F2, GALE, CLIC4, STMN1, ZNF593, RCC1, YARS, AK2, PSMB2, CLSPN, MRPS15, DNALI1, CDCA8, MYCBP, CTPS1, EBNA1BP2, CDC20, DPH2, B4GALT2, CCDC24, SLC6A9, TMEM53, KIF2C, HPDL, AKR1A1, NASP, MAST2, STIL, CDKN2C, ORC1, LRP8, NDC1, LRRC42, MRPL37, DHCR24, ITGB3BP, PGM1, AK4, DEPDC1, PSMA5, ATXN7L2, CSF1, WDR77, SLC16A1, DCLRE1B, VANGL1, TTF2, PHGDH, FAM72B, FAM72D, FAM72C, ANKRD35, ANP32E, PMVK, CKS1B, ADAM15, FDPS, SEMA4A, GPATCH4, MRPL24, DUSP23, FCER1G, TOMM40L, NUF2, UCK2, XCL2, XCL1, PRDX6, CENPL, DARS2, CACYBP, IER5, RGS16, NPL, RGS1, ASPM, KIF14, PHLDA3, TIMM17A, PTPN7, UBE2T, SNRPE, FAM72A, NEK2, DTL, ATF3, BATF3, CENPF, LIN9, COA6, EDARADD, FH, EXO1, SCCPDH, PFKP, NET1, IL2RA, MCM10, SUV39H2, MASTL, CREM, C10orf128, TIMM23, ZWINT, CDK1, ADO, EGR2, DNA2, PPA1, PCBD1, DDIT4, DNAJC9, PPIF, EIF5AL1, FAM213A, FAS, KIF11, CEP55, HELLS, RRP12, PGAM1, ZDHHC16, GOT1, SCD, NPM3, PPRC1, NFKB2, SFXN2, SFR1, GSTO1, DUSP5, SFXN4, PRDX3, MKI67, GLRX3, STK32C, TALDO1, SLC25A22, POLR2L, TSPAN4, MRPL23, PHLDA2, RRM1, MRPL17, C11orf16, WEE1, RRAS2, SAAL1, LDHA, ZDHHC13, E2F8, CCDC34, KIF18A, PRRG4, HSD17B12, PSMC3, MTCH2, SSRP1, SLC43A3, TIMM10, FAM111B, FEN1, FADS1, FADS2, COX8A, STIP1, PPP1R14B, GPR137, SAC3D1, CDCA5, POLA2, SSSCA1, BANF1, MRPL11, DPP3, ZDHHC24, RBM14, LRFN4, GSTP1, NUDT8, NDUFS8, C11orf24, PAK1, NDUFC2, ALG8, DDIAS, TMEM126A, SMCO4, KDELC2, ZW10, ZPR1, HMBS, H2AFX, HSPA8, FEZ1, CHEK1, DCPS, NCAPD3, FKBP4, FOXM1, RHNO1, CCND2, RAD51AP1, NDUFA9, NCAPD2, GAPDH, LAG3, CDCA3, TPI1, EMG1, MAGOHB, HEBP1, BCAT1, ARNTL2, DDX11, VDR, TMEM106C, TUBA1B, TUBA1A, TUBA1C, TROAP, RACGAP1, ESPL1, CDK2, IKZF4, PA2G4, MYL6B, NABP2, SLC39A5, TIMELESS, PRIM1, NEMP1, SHMT2, NDUFA4L2, MARS, CDK4, METTL1, XPOT, IL26, YEATS4, CCT2, PHLDA1, E2F7, DUSP6, UBE2N, SOCS2, SNRPF, PARPBP, PMCH, NT5DC3, CORO1C, UNG, FAM222A, OAS3, TESC, RFC5, COX6A1, KNTC1, CDK2AP1, EIF2B1, SCARB1, BRI3BP, RAN, POLE, PXMP2, SKA3, MIPEP, POMP, SLC7A1, HSPH1, BRCA2, RFC3, DNAJC15, LACC1, DIAPH3, NDFIP2, ABCC4, TEX30, TFDP1, MRPL52, PRMT5, HAUS4, PCK2, EMC9, PSME2, GZMB, PSMA6, NFKBIA, LRR1, POLE2, CDKN3, WDHD1, DLGAP5, HIF1A, MTHFD1, ACTN1, ERH, ACOT4, C14orf1, TMED8, ASB2, IFI27L1, WARS, ANKRD9, TRMT61A, XRCC3, CEP170B, CDCA4, TMEM121, ARHGAP11A, BUB1B, KNSTRN, RAD51, CHAC1, OIP5, NUSAP1, WDR76, FRMD5, SQRDL, DUT, ATP8B4, SLC27A2, CCNB2, TIPIN, ZWILCH, AAGAB, CLN6, KIF23, PKM, SEMA7A, COX5A, RPP25, FBXO22, ETFA, IDH3A, FAH, TM6SF1, HDGFRP3, NMB, MRPS11, HAPLN3, TICRR, KIF7, BLM, PRC1, SNRNP25, WDR90, CCDC78, CHTF18, GNG13, CCNF, C16orf59, PAQR4, PKMYT1, THOC6, MMP25, CIITA, SOCS1, RMI2, EARS2, NDUFAB1, PLK1, ERN2, IL4R, BOLA2B, BOLA2, HIRIP3, ALDOA, DCTPP1, ZNF267, SHCBP1, ORC6, GPT2, NETO2, RPGRIP1L, CRNDE, MT2A, MT1E, MT1X, CFAP20, GINS3, FHOD1, PARD6A, ENKD1, NUTF2, SLC12A4, DUS2, COG8, PDF, NQO1, AARS, KARS, CENPN, MPHOSPH6, GINS2, IRF8, C16orf95, SLC7A5, CDT1, FAM57A, YWHAE, HIC1, SPNS2, PSMB6, C1QBP, C17orf49, EIF5A, CD68, MPDU1, SOX15, TP53, AURKB, PFAS, COPS3, DHRS7B, C17orf51, SPAG5, ERAL1, COPRS, NLE1, CCL3, PIGW, CISD3, PSMB3, PSMD3, MED24, CDC6, TOP2A, ZNF385C, TUBG1, PSME3, BRCA1, C17orf53, KIF18B, COPZ2, ATP5G1, GNGT2, PHB, EME1, LRRC59, NME1-NME2, NME1, MMD, DYNLL2, PRR11, PTRH2, BRIP1, C17orf58, KPNA2, HN1, TEN1, TK1, BIRC5, SOCS3, EIF4A3, MRPL12, SLC25A10, ALYREF, ANAPC11, MAFG, PYCR1, FASN, SLC16A3, TYMSOS, TYMS, YES1, NDC80, NDUFV2, IMPA2, SNRPD1, RBBP8, TAF4B, INO80C, TPGS2, HAUS1, IER3IP1, ACAA2, SKA1, PMAIP1, BCL2, RBFA, TIMM13, LMNB2, THOP1, GNA15, DOHH, CHAF1A, MYDGF, C19orf70, DUS3L, RFX2, CLPP, CD70, SH2D3A, CD320, DNMT1, ICAM1, PDE4A, LDLR, KANK2, FBXW9, RNASEH2A, FARSA, CALR, NACC1, C19orf57, ASF1B, DDX39A, TPM4, CCDC124, IFI30, MPV17L2, LSM4, ARMC6, NR2C2AP, NDUFA13, ZNF85, ZNF681, CCNE1, GPI, ZBTB32, OVOL3, PSMD8, NFKBIB, SARS2, TIMM50, DLL3, PSMC4, EXOSC5, PAFAH1B3, SMG9, BCL3, TOMM40, RELB, CD3EAP, ERCC1, PPP5C, SLC1A5, SAE1, KPTN, LIG1, MAMSTR, BCAT2, RUVBL2, IL4I1, ATF5, POLD1, C19orf48, HSPBP1, UBE2S, ISOC2, RRM2, C2orf48, ODC1, ATP6V1C2, PDIA6, GEN1, ADCY3, CENPA, FAM98A, GEMIN6, MORN2, EPAS1, MSH2, MSH6, CHAC2, COMMD1, SLC1A4, SNRPG, FBXO41, BOLA3, MTHFD2, SLC4A5, HK2, MAL, ITPRIPL1, NCAPH, PDCL3, BUB1, CKAP2L, DBI, TMEM177, NIFK, IMP4, CCDC74A, MCM6, PSMD14, GCA, NOSTRIN, SPC25, KLHL23, HAT1, ZAK, CDCA7, ATP5G3, CCDC150, HSPD1, HSPE1-MOB4, HSPE1, TMEM237, WDR12, BARD1, MREG, CHPF, FARSB, SERPINE2, HJURP, DTYMK, FKBP1A, SNRPB, NOP56, C20orf27, PCNA, CRLS1, NXT1, GINS1, NANP, BCL2L1, TPX2, E2F1, EIF2S2, AHCY, ROMO1, DSN1, FAM83D, MYBL2, SDC4, UBE2C, AURKA, CTSZ, C20orf197, ADRM1, SLCO4A1, SLC17A9, DONSON, CBR3, CHAF1B, PSMG1, CBS, RRP1, FAM207A, SLC19A1, C21orf58, PEX26, UFD1L, CDC45, RANBP1, SDF2L1, C22orf15, CHCHD10, DERL3, MIF, CHEK2, NEFH, LIF, OSM, MTFP1, PES1, SMTN, YWHAH, MCM5, FOXRED2, NCF4, PDXP, DMC1, APOBEC3A_B, ATF4, ADSL, RBX1, DESI1, CENPM, NDUFA6, PARVB, RIBC2, GTSE1, MAPK11, PLXNB2, TYMP, BHLHE40, CAMK1, LOC401052, FANCD2, LSM3, MLH1, CSRNP1, KIF15, KIF9, CDC25A, IMPDH2, LSMEM2, IFRD2, HYAL3, CISH, MAPKAPK3, MANF, POC1A, STAB1, NT5DC2, RFT1, C3orf14, THOC7, SHQ1, CMSS1, NIT2, KIAA1524, ZBED2, ARHGAP31, ADPRH, COX17, POLQ, MCM2, SRPRB, ATP1B3, GMPS, SMC4, RPL22L1, ECT2, ECE2, CAMK2N2, DNAJB11, RPL39L, TFRC, SLBP, S100P, TRMT44, NCAPG, PGM2, PTTG2, SCFD2, PAICS, MRPL1, COQ2, HPSE, H2AFZ, SLC39A8, CENPE, OSTC, TIFA, ZGRF1, MAD2L1, CCNA2, PLK4, MND1, C4orf46, TMA16, HMGB2, SAP30, NEIL3, CENPU, ANKRD37, TRIP13, CCT5, ANKRD33B, TARS, NUP155, HMGCS1, DEPDC1B, CENPK, NLN, CCNB1, CENPH, ARSB, DHFR, C5orf30, LMNB1, ISOC1, IL13, SEPT8, UQCRQ, VDAC1, H2AFY, KIF20A, CDC25C, EGR1, SPATA24, GM2A, ATOX1, KIF4B, HAVCR2, ADAM19, PTTG1, HMMR, RPL26L1, SFXN1, HRH2, NOP16, GPRIN1, PRELID1, MXD3, NHP2, HNRNPAB, IRF4, TUBB2A, TUBB2B, LYRM4, BMP6, TXNDC5, PAK1IP1, TBC1D7, MCUR1, CD83, GMNN, HIST1H3C, HIST1H4C, HIST1H2AE, HIST1H2BH, HIST1H2AI, HIST1H2AJ, HIST1H1B, NRM, TUBB, IER3, VARS2, TCF19, LTA, TNF, LST1, AIF1, APOM, MSH5, VARS, NELFE, HLA-DQA1, KIFC1, UQCC2, SNRPC, CDKN1A, PPIL1, CCDC167, KCNK5, MED20, BYSL, DNPH1, MRPS18A, VEGFA, MRPL14, SLC29A1, HSP90AB1, CENPQ, MCM3, BAG2, PRIM2, PTP4A1, SDHAF4, KCNQ5, MB21D1, TTK, MMS22L, BEND3, CEP57L1, RPF2, DSE, CENPW, EPB41L2, SGK1, MTFR2, STX11, GINM1, LRP11, MTHFD1L, FBXO5, PRKAR1B, NUDT1, SNX8, IQCE, TTYH3, FSCN1, MIOS, RPA3, NDUFA4, BZW2, AHR, TWISTNB, DNAH11, CDCA7L, NFE2L3, SNX10, GGCT, GARS, LSM5, ANLN, PSMA2, BLVRA, POLD2, PPIA, FIGNL1, SEC61G, LANCL2, MRPS17, PSPH, CCT6A, CHCHD2, ASL, STX1A, ABHD11, LAT2, RFC2, CDK6, ASNS, PDAP1, ATP5J2, MCM7, POP7, AP1S1, PUS7, NAMPT, MDFIC, HILPDA, STRIP2, CHCHD3, BPGM, TTC26, NDUFB2, EZH2, XRCC2, NCAPG2, AGPAT5, ERI1, |

| | |
|---|---|
| | SLC39A14, CDCA2, ESCO2, PBK, DUSP4, UBXN8, EIF4EBP1, GINS4, AP3M2, MCM4, MRPL15, GGH, RRS1, FABP5, TMEM64, RAD54B, FSBP, COX6C, NUDCD1, DSCC1, MRPL13, MTBP, ATAD2, SQLE, MYC, TSTA3, FAM83H, EXOSC4, BOP1, SLC52A2, RECQL4, RCL1, TMEM261, CDKN2A, NDUFB6, KIAA1161, SIGMAR1, DNAJB5, FANCG, MELK, TOMM5, ALDH1B1, C9orf40, PSAT1, CKS2, GADD45G, NFIL3, IARS, ASPN, FANCC, ZNF367, AAED1, SMC2, NIPSNAP3A, TMEM38B, CTNNAL1, TXN, SLC31A1, RBM18, NEK6, ARPC5L, HSPA5, SLC2A8, PTRH1, FPGS, WDR34, PKN3, ZDHHC12, NCS1, VAV2, NOTCH1, SAPCD2, TUBB4B, FAM166A, FANCB, PRDX4, APOO, POLA1, RBM3, CLCN5, HSD17B10, PDZD11, KIF4A, ERCC6L, COX7B, PGAM4, PGK1, CSTF2, NOX1, TIMM8A, PGRMC1, SLC25A5, UBE2A, ZDHHC9, HPRT1, HMGB3, HAUS7, IRAK1, LAGE3, G6PD, MPP1, VBP1 |

### Table S2. List of IL-15-specific and TCR-specific gene sets.

| | |
|---|---|
| **IL-15-specific gene set (bystander activation gene set)** | ZNF683, CES1, CXCR1, NME8, KLF2, NLRP3, FRMPD3, LY9, NHLRC1, MEOX1, PCSK5, CMKLR1, ASAP2, WNT10B, ZBP1, NMUR1, SLC22A3, SLC1A7, LGR6, TREML2, TMEM163, TGFBI, CX3CR1, DNAJC28, MS4A1, PTGDS, SLC40A1, KLRF1, HKDC1, NCF1, TRPM2, KCNA6, RSAD2, KLF8, ZNF540, KIF19, CEACAM1, MYO3B, CFH, ARRDC5, CYP4F22, MTSS1L, PDGFRB, TPBG, ADM, RPA4, TMEM229B, ENPP5, PAPSS2, SPON2, CHI3L2, GOLGA7B, MYOM1, FCRL6, RAB27B, ADRB2, SNX30, CPNE5, HSPA6, FHAD1, PYROXD2, ZNF781, ENC1, TSPAN32, SIGLEC9, GIMAP1, GDPD5, GIMAP1-GIMAP5, TRIB2, MYBL1, TGFA, CD300A, C3AR1, ZNF547, PNMA3, ZNF365, PTGDR, IFIT2, COL5A1, ANXA2R, C1orf162, RASGRP2, ATL1, CLIC3, LZTS1, AKR1C3, ZNF300, RHOU, SLC51A, LOC100505478, CFAP53, COL9A3, GCNT2, FGFBP2, FAM26F, ADAMTS1, FAM229A, ERP27, PLCH2, CDK5R1, RGS17, DNAH10, SOAT2, SLC4A4, C2CD2, HPGDS, SLFN5, RAB37, IFIT1, CLU, POU5F1B, DOCK5, TXNIP, ZNF155, PTPRN2, TNFSF13B, ACVR2A, CDC14B, PRH1, SMC1B, PLAC8, GOLGA8K, GNRH1, ZNF416, LAIR1, TMCC3, SMPD3, SPRED2, PDGFB, FGL2, GOLGA8R, SLCO4C1, PTPN4, ARHGEF28, VLDLR, CNTLN, GOLGA8N, RARRES3, ZNF879, S1PR5, ADHFE1, ITGAM, AMY1C, SLC2A14, TBC1D3B, FCGBP, SPRED1, LPAR6, IL6R, FRY, FCGR3B, CD244, GOLGA8O, GSN, TCF7, CD52, RCBTB2, TRANK1, WNT1, CALHM2, FCGR3A, LYZ, BNC2, CD93, ISG20, SUSD1, SPNS3, ITGA6, AMY1A, EXOC6B, SCML4, ZNF460, PPL, UBXN11, PARP15, GOLGA8H, CTSO, ERBB3, FAM47E-STBD1, NTNG2, MCF2L2, AQP3, ZNF570, APOBR, ARMC2, SYNE1, JAZF1, SLCO3A1, VNN2, C1orf204, IGFBP4, CECR1, C20orf197, PXN, NEIL1, FZD6, ZMYM1, C15orf53, TANC2, FOXD2, EPHA4, PDGFD, HRASLS2, IL16, CCDC146, XAF1, C16orf54, DDAH2, SNTB1, LGALS9C, APBA2, RAP2A, BFSP1, MTSS1, IFITM1, CCDC141, GIMAP7, SLC9A9, GOLGA8J, CXorf65, ZSCAN12, ZFYVE9, FBXO48, AUTS2, DDX60L, BTLA, FAM231D, RHOBTB3, PDLIM2, ADARB1, GRAP2, MTMR7, RGL4, SAMD3, SBK1, AMY1B, TXK, DDX60, ZNF443, KIAA1551, SNX20, ADM2, SERPINI1, DNMBP, PKD2L2, ETV3, ATM, RASSF4, TMIGD2, C5orf58, GIMAP2, LILRB1, PHLDB2, CDRT4, CCR6, CARD6, MEGF6, AKTIP, GPRASP1, SSBP2, LGALS1, CCDC65, PLEKHF1, TCF7L2, DNAH14, TNC, FMN1, TICAM2, ZNF799, SLC4A8, C1QTNF3, AHNAK, RGS9, TCEANC, WDFY1, GIMAP4, ANXA1, BCL6, DOK2, B3GAT1, RNF122, ST8SIA1, ARHGEF3, SUN2, TARP, HDAC4, PATL2, CD101, JAG2, CHN2, ADAMTS10, MMP25, LMTK3, PIM1, ARL4C, MCTP1, PPP2R2B, SYNGR1, SBF2, ZFP3, BIN2, KIAA1841, SGK3, MDGA1, FGR, CD69, CARD16, TMEM71, ANTXR2, SPTB, KIR2DL4, XYLT1, AMT, SMAGP, NOL3, IRF7, DNM3, AGTRAP, SEMA6A, UTRN, GZMA, CCDC126, SAMD9L, S100A4, CACNA2D2, TRAT1, FAM129B, SLC16A6, ZFP36, MGAT4A, ZNF417, EFNA4, CYTH3, SYTL1, RAP1GAP2, KLRB1, CASP10, SYNE2, AMPD3, ZNF550, ZNF470, APOBEC3H, FBLN5, ITM2B, PDP1, NDRG1, ATP10D, BCL7A, TMEM154, C21orf62, CAPG, ZNF627, PDLIM1, TNFSF8, SLC2A12, TIMP1, RNF166, TRIM22, STAT4, GATS, LMO7, MYO1F, TMEM50B, MBP, BBS10, LDLRAP1, TM6SF1, ENPP4, IL18RAP, SNAI3, CELF6, ELOVL6, GNG2, LAPTM4B, PTAFR, AKAP5, ZBTB20, TSHZ3, FAM46A, SMIM14, IL18BP, CXCR5, SLAMF7, RTKN2, DMTN, GZMM, SELPLG, ZNF154, B3GALT4, C1orf21, IFI44, KIR2DL1, PREX1, GOLGA8M, FAM102B, B3GALT2, CBX4, ZFYVE28, CLIC5, ZNF226, GPR68, TSPAN2, EVI2B, CMPK2, ZNF789, TSPAN14, TC2N, TBC1D10C, PDZD4, RTP4, TMEM116, ZNF652, KLRK1, ZFP14, MCC, SAMD9, GIMAP5, PITPNC1, KLHL28, ZNF235, PLCB1, FAM65B, DNAJB9, FAM131B, GAB3, TBC1D19, HELZ2, PLA2G4B, KLRC4-KLRK1, ITGB2, RASGRP1, GLIPR2, EPSTI1, IFI44L, YPEL1, ADAMTSL5, LIPT1, SLC5A10, MXD4, HOXB3, HBEGF, CA5B, HCST, XBP1, SPOCK2, CCR5, LIG4, KPNA5, KIR3DL2, ZNF674, FLT3LG, ZNF14, ZBTB41, GIMAP6, DTX3L, GYG1, HERC6, SLC25A53, PIK3IP1, IL18R1, ISG15, C14orf28, PLEKHG3, PARP8, ZNF304, ZNF549, MX2, PRSS23, PLEK, IFITM2, CD200R1, RNF157, MX1, CC2D2A, ZXDB, HLA-DRB5, NATD1, EIF5A2, ARNTL, RBL2, RASA3, ZNF845, STOM, ZNF75D, FBXO6, RAB33B, ELMOD2, GIMAP8, SLAMF6, CITED4, ZNF559, ARRDC3, SH2D3C, ALOX5AP, SCRN1, KIF21B, MYL12A, OASL, ZBTB18, KIR3DL1, TOX2, LTB, ZNF568, C5orf63, ARSD, TCF7L1, ZFP36L2, ZNF681, CROT, SLC46A3, ZNF484, NOL4L, LRMP, TMSB4X, PTGER4, SH3BP5, GPR35, GLI1, PBXIP1, ASCL2, CYP2R1, THEMIS, BTD, A2M, KAT2B, ZMAT1, RDH5, ARRB1, AKNA, SOS1, THEMIS2, PYCARD, SIT1, SSH2, TSC22D3, ERMP1, TSC22D1, SORL1, PVRIG, TYROBP, TMEM107, GGT7, BTN3A1, FBXL16, SESN3, TTC16, CLEC2B, KIAA0040, CASC1, IQSEC1, FAM50B, ZNF831, SPSB2, TNFSF12, ZNF274, PRNP, NUAK2, CFAP44, LIME1, RNPEPL1, ZNF134, EVL, IFNGR2, ZNF790, MOAP1, ITGA5, GBP3, HS3ST3B1, TNRC6C, KBTBD7, ZCCHC14, FAM8A1, ADCY7, MAD1L1, KIAA1468, ITGAV, ERO1B, LNX1, CD53, PRKACB, APOBEC3G, TRPV2, PLOD3, MANBA, DLG5, IFITM3, IFIT5, MIDN, TRAPPC2, DPYD, MORC3, SENP7, WTH3DI, SIGIRR, PTGER2, CEBPD, AKAP7, IKBKE, RGS19, ZNF28, C1RL, OSTM1, ZNF786, ITGB1, ZNF792, CPQ, ANKRD44, NSG1, ITK, MATK, PCSK7, NUCB2, ZNF575, PZP, CLUAP1, ZNF573, MYO5B, ZNF528, TNIK, ZNF260, CASP8, BCL9L, ZNF33B, MMD, TBC1D32, C3orf18, ZNF264, GPR18, KLRG1, VAMP4, TOB1, DYRK2, GALNT6, BEX2, TMEM63A, YPEL5, NLRC3, FAM63B, GPSM3, OSBPL7, BLZF1, TGFBR1, IL17RA, ZNF480, SH3BGRL3, GSAP, ZNF429, CRIP1, FYB, ABHD13, CYTH4, POLI, SAP25, SIDT1, LAPTM5, ARHGAP15, ZNF84, RPS6KA2, FLJ44635, AIM1, SLC26A11, KLF6, GNAL, TRIM62, NLRP1, EPOR, C1orf56, PARP9, PHKB, ADD3, IFIT3, EVI2A, PRKCH, YPEL3, GPR137B, NKG7, SLC17A5, RGAG4, CNN2, PTPRE, NRIP2, ZNF493, RNF44, FAM199X, NLGN3, CDKN2D, ACVR1B, UBA7, PAFAH2, ADRB1, FNDC3B, RAP2B, LFNG, SYNRG, STK38, GLRX, ZFAND6, LRIG2, C6orf47, CTSD, LAIR2, SVIL, RASSF1, ARHGDIB, ZNF350, PLCL2, VAV3, TPST2, CD96, GLG1, AGAP2, C11orf63, METTL18, PCED1B, RGS14, TLE4, ATXN7L1, TRDMT1, CALCOCO2, IFNAR2, ST3GAL5, STK10, CLN5, INPP5D, PTPRCAP, LNPEP, ZNF567, CHST12, ZNF200, IL10RA, SGTB, FBXO44, LYSMD3, GPR65, KLF3, CDC14A, C20orf194, FAM126A, CASP1, C3orf58, FAM107B, SEC31B, FAM84B, APBB3, IL7R, S1PR1, PTCH1, TMEM80, KANSL1, UBL3, SHISA5, CDC42EP3, MAPK7, TLDC2, GIN1, CD99, ADAM10, CPD, MFAP4, TENM1, OAS1, OSBPL5, SKI, SLC27A3, SFMBT2, TRIM21, APOL6, TES, GTF2IRD2B, CCDC112, TRIM5, MTHFR, PTPRC, CORO2A, GLUL, PLSCR1, ACAP1, CCDC107, PNRC1, KLRC3, FAM117A, ARAP2, SLC25A42, RAB6C, SLC2A3, PYHIN1, HLA-DPB1, CD48, BCL11B, PLCG1, ANKAR, IRF9, SAMHD1, JAKMIP2, PIKFYVE, PLCD1, KLF12, DIP2A, PTPN9, KLHL24, COLQ, VAMP2, FAM210B, GZMH, TMEM65, CREBRF, ZNF319, CCND3, ZDHHC20, LPCAT4, IRF2BPL, TULP4, WDR37, ZNF718, CLCN6, IPCEF1, KIAA0355, ARHGAP12, ATP11B, RTN4R, DAZAP2, DBP, RHOC, HEG1, PRR5L, ARHGAP9, SYTL2, SLC35F6, CD5, NBEAL2, SUSD3, ZNF18, ZC3H6, GABARAPL1, SARAF, RFK, LPP, TTC14, ARHGAP27, CAPN2, ARL6IP6, PLEKHG2, RNF144A, BTN3A2, NCR3, MAN2B2, KRCC1, SETD7, HLA-E, GBP1, IFNLR1, AGO4, F11R, HACD4, ZNF554, WBP1L, TRIM14, PWWP2B, PLEKHO2, GRAP, PAN2, AXIN1, FABP3, ZNF189, FAM131A, P2RY8, VIM, PPP1R3B, SESN1, MPP1, TMC6, IL24, UMAD1, WIPI1, KIR2DS4, SLC15A4, PRKD2, IFI6, ZMAT3, PPP1R18, CCDC88C, CISH, FGFR1, PTPN12, BID, FPGT, EIF4E3, RNF111, APOL3, F2RL2, PARP10, SASH3, PQLC3, MICAL1, BLOC1S3, PLEKHA1, MYD88, LPGAT1, HPGD, LIPA, DDX58, PTPN22, STX3, ERAP2, MAPK1, ITGAL, CTSW, SNX29, TMEM251, ZKSCAN8, FGD3, GNB4, TNFRSF1A, ZNF641, S100A10, MAP4K4, TBC1D20, LGALS9, SNRK, FGFR1OP2, S100A6, GLCCI1, ARHGEF6, RORA, DEDD2, DCK, MR1, MAN2C1, TNFSF14, RNASEL, NEMP2, ARHGAP35, ZNF160, LYSMD1, ARHGEF1, SLC44A2, ANO9, ZNF708, USP9Y, NEDD9, ZNF700, SYPL1, CTSC, CPEB2, C7orf49, NABP1, RAB8B, CHAC1, RASSF2, DOCK9, SP100, NLRC5, CAMK2G, PDE7A, C7orf25, ZYX, MPPE1, CD8A, ZNF394, ARHGEF39, SLFN12L, ZNF805, SLC9A3R1, FAM200B, MYADM, TTC39C, FEZ2, CDC42SE1, TCTA, SH3BP2, PDE4D, YES1, PIK3R1, H2AFJ, TBC1D23, IFNGR1, ZNF217, ARL2BP, IGIP, IGF2R, UTY, C12orf75, FAM49A, CMIP, DENND3, NPHP3, TRAF5, NBPF3, PHYKPL, UTS2, STK17A, PHF1, FCMR, HBP1, TRIM38, PARVG, ATP1B1, GK5, HLA-DRB1, ZNF737, IRS2, TMC8, ARHGAP30, LZTS2, RBMS1, LOC728392, CAPRIN2, APBB1IP, FBXL5, COLGALT2, INPP4A, ZNF548, CPEB4, MYO6, LEMD3, STMN3 |
| **TCR-specific gene set (TCR-induced activation gene set)** | IL9, CXCL13, IL5, DHRS2, IL3, NR4A3, DACT3, GALR2, COL6A3, PTGIR, XIRP1, FNDC9, IGF2, INS-IGF2, ZBED2, TIMD4, F5, IL1RN, GNG8, PANX2, OLIG3, CCL17, PHKA1, IL1R2, MYOF, ACOXL, C1orf53, VWCE, DNAJC12, IL4, ADCY1, CD200, EBI3, SPR, TRIB1, NPW, IL2RA, MNS1, SHF, IL10, TMEM217, COL5A3, PDLIM4, EVI5, ANKLE1, TMEM198, IL13, CTF1, GBP6, POU2AF1, SMAD1, SHC4, DGAT2, GNG4, HOMER2, TNFRSF4, POU3F1, C17orf78, CARD9, MYB, MEX3A, ABTB2, CTTN, FSTL3, MAMLD1, NDST1, CSF2, AEBP1, PSD, LMCD1, LIM2, FZD5, PRKAR2B, STAP2, VSTM4, DIXDC1, CCL3, NR4A1, CCR4, SIX3, CTH, NR6A1, CTLA4, KCNH4, EPAS1, SOX8, TWIST1, GLB1L3, TTC8, PRKCDBP, SLC41A2, TNFRSF9, CCL3L3, HOXB9, CLUL1, PRSS27, CCL3L1, DOCK6, PEX11G, EGR3, VDR, ZBTB32, SLC4A11, CCL4L1, CCL4L2, CCR7, RYR2, MT1G, CCDC38, SLC35F3, SORBS1, CENPI, CD86, LOC388780, EGR2, FKBP7, IFNG, NUP210L, CD109, HPDL, P2RY13, CLEC11A, ERRFI1, MB, TNFRSF8, RGS16, LYPD3, LKAAEAR1, DUSP4, C6orf132, NCR3LG1, HACD1, NAPSA, GLIS2, PDGFA, MET, TUFT1, MYC, PPP1R3F, SCAMP5, VSIG10L, ZC3H12C, CCL22, IL17D, LGMN, CLPB, TP53INP2, IRF4, SNX33, FOSL1, SMOX, DNAJC6, RAMP1, FABP5, GCAT, MT1H, MED12L, ADGRF3, ELL2, MFSD2A, NR4A2, CHRNA5, PTGIS, TMPRSS3, LAYN, RBFOX2, FOSB, FOS, STAB1, CCL4, PIM2, TMEM178B, GPR19, NT5DC2, B3GNT5, SHROOM1, MAPK12, HIST1H4E, ZFHX2, HIST3H2A, HRH1, NARS2, TMEM234, SOCS6, MESP1, FSD1, CHRNA10, PLAGL1, PYCR1, C9orf116, CRTAM, NAMPT, ANKRD13B, SETMAR, ACTN1, SCARB1, FBXW9, DLL3, PACSIN3, HIST1H4H, ATP6AP1L, BCL2L11, ZNF768, DUSP2, SLC9A7, ACY1, TLN2, GNA11, METAP1D, BATF3, IL12RB2, POLR3G, CSF1, LAG3, SOX4, FKTN, PDIA5, GLB1L2, CCL20, GSG2, AFMID, SORD, TPRG1, TFRC, GJC1, ARL10, RDH10, SRR, NPM3, FAM216A, LMNA, MYL6B, ST6GALNAC2, MACROD1, HSPD1, AUH, HLA-DOB, GALM, ECM2, IQCG, PRDM11, RGS1, ICAM1, NME1, FBXO43, TFAP4, ADGRE2, PTGR2, OAZ3, DZANK1, GRAMD1B, SLC43A2, HIST1H2AE, TNFRSF18, PHLDA1, DHRS11, |

| | |
|---|---|
| | *CORO6, KCNK5, PHLDB3, SMYD5, ZC3H12D, KCNC4, AGRN, C4orf32, MYOT, ARHGAP11B, TRAP1, RRP12, MRC2, NLN, DNAH11, ACBD7, MCAT, LNP1, MRPL12, CCNE1, FLT1, HOOK2, TSEN2, YBX3, NLE1, CDCA7L, CAMK1, SNX9, NR2F6, NFIL3, COPRS, CYP1B1, CCNB1, FOXRED2, OAF, ARID5A, SLC29A1, TOMM20L, GADD45G, ZNF713, HOPX, PSEN2, TICRR, POLD2, C17orf53, TYMSOS, PMCH, AUNIP, PEMT, IGFLR1, PIM3, PUS7, KCNN4, PRMT5, ADAM9, MEGF8, B4GALT2, HNF1A, CCDC6, IL4I1, BRICD5, KIF7, ALDH1B1, DNAJC18, LOC81691, TNIP3, RBPJ, SLC38A5, NEFH, SLC35F2, DNASE1, EXO1, CLUH, MECR, TMEM121, TSPAN33, GCSH, SRM, VARS2, MRM1, TMEM201, SFXN4, HOMER1, SLC27A2, FCRL3, LTA, ZNF584, HAGHL, GDF9, PYCRL, NPM1, MYBBP1A, IMPDH2, NFIA, MRPL4, HK2, HIST1H1B, KDM6B, MRTO4, IL23R, TROAP, ESCO2, PIH1D2, PRMT1, FAHD2B, IMMP2L, WDR12, TIAM2, CTPS1, MEST, ORC1, ARNTL2, COPG2, PLCD4, PFAS, MRPL37, NOP16, NETO2, SKA3, AURKB, MAF, P4HA2, AGMAT, LRRC8B, MLST8, SLC25A10, MAP1LC3A, RRM2, ACACA, GZMB, HMGA1, KLHL23, CBR3, TAF4B, OGG1, IER3, RPL26L1, CAD, IL23A, LSMEM2, STRIP2, CCDC134, CDCA5, NINJ1, GAMT, PLAGL2, HSPE1, KIF3A, TRIP6, PAOX, CD3EAP, TBRG4, ERN2, SLC25A19, C20orf27, BIRC5, PES1, FHL3, CDC20, CENPA, WDR3, SUCLA2, ETV6, TNFSF9, GK, IFRD2, NOP14, TLCD1, ANKRD16, C19orf48, DPH2, TRIM16, TST, ALCAM, FASN, NCS1, HSPE1-MOB4, LRRC20, PAICS, NOLC1, ACTA2, C16orf87, SLC43A3, TOMM40, C10orf128, TSR1, PHLPP1, SKA1, AHI1, C16orf59, RPL22L1, TTLL12, EED, FRMD4B, FUOM, HSP90AB1, PLK1, ANKRD34A, TIMM13, ALKBH2, SLC27A4, CMSS1, ANKS6, HDGFRP3, PRKCE, PLD6, RCC1, ITPRIPL2, CCNI2, ECE2, SHCBP1, BOP1, STAT5A, SLC5A6, SPIRE1, RYBP, METTL1, PRMT3, KPNA2, C16orf95, CDC25A, PARS2, PALLD, SLIRP, FKBP4, PCCB, CCDC86, ZSCAN5A, ODC1, EGR1, GNL3, LIMA1, BATF, PTMS, PBK, PPARGC1B, ZNF771, SPNS2, SWI5, SLC17A9, SRD5A1, ITGAX, TRIP10, SPC25, SFXN2, DTWD2, C2orf48, DRAM1, DIAPH3, MKS1, NCL, PELI1, SATB1, TJP2, ESPL1, CDCA3, MRPL3, KIAA0408, SLC29A2, NDUFA4L2, GPX1, PINX1, IMP4, MRPS26, DNPH1, HIVEP1, RRP9, ZNF692, RPP40, DKC1, MAPK6, GNB1L, FAM83D, TRAF4, TOMM34, FAM69B, ASPHD2, PARVB, SIGMAR1, MTURN, PSMB5, MTRNR2L4, CCDC51, PDCD2L, WDR43, THG1L, NOP56, RRNAD1, MRPS2, FADS1, CXorf40A, ISOC2, NFKB2, DNALI1, UCK2, RRP1, ATIC, BUB1B, RECQL4, AHCY, NCAPG, MTA3, THOP1, POLQ, WDR74, SIAH2, FURIN, SDC4, NCALD, PPAT, ZMYND19, FDX1L, ATP8B4, NHP2, USP12, YRDC, FAM136A, PKMYT1, MYH10, GOT2, BZW2, FAM72A, FOXP3, C17orf51, TASP1, HIST1H2AI, WDR4, GMDS, EIF2D, DHX34, PKM, ADCY3, NOP2, GEMIN4, HIST1H2AJ, TRAIP, TMCC2, ARMC6, NFE2L3, ADK, ACOX3, TIMM50, MRPL17, CKB, TIMM8A, PTGDR2, SSSCA1, APEX1, ATF5, NDUFV2, MSC, CA11, CD70, FARSB, TMEM97, PHLDA2, EXOSC5, KNSTRN, NME1-NME2, STOML2, RANBP1, CCDC58, HDAC5, DHX33, UBE2T, SLC39A14, CTNNAL1, C22orf15, WARS, GEMIN6, MELK, MTRNR2L6, HSPA4, MORN2, SOX12, TNFAIP3, TUFM, SCD, CDK6, CD151, PPA1, CCT3, FAM72B, HMBS, LYST, DLGAP5, B4GAT1, EBNA1BP2, NCAPH, DERL3, GOLIM4, POLR2H, MRPL21, FAHD2A, ARL2, NOB1, RUVBL1, TRMT1, TFB2M, TRABD2A, HMGN5, MRPS17, ATAD3B, FAM207A, CENPO, GART, TKT, C17orf58, ZNF593, DLAT, THAP4, HIST2H2AA4, RFC3, FAAP24, DDX31, GEMIN5, TGFBR3L, RAN, HMMR, YIF1A, MYBL2, CYB5D1, SLC37A3, NDFIP2, LARS2, CHPF, PNPO, SHMT2, ADAP1, CHCHD10, HDLBP, SLC25A33, PIF1, LYRM4, DDX49, TRAF1, FAM212B, BCL2L1, NAT14, PRKAR1B, ABCF2, KAT2A, PPM1N, EXOSC4, SMS, RANGAP1, RPL7L1, UBE2C, CDCA2, MND1, SNAPC4, PHB, SERBP1, SMTN, DOT1L, PGAM1, MPV17L2, OIP5, MFSD3, HIST2H2AA3, GPATCH4, LRPPRC, MRPL15, PAM16, SLC37A4, L2HGDH, DCTPP1, MTX3, SLC39A4, RPS6KA4, CPSF3, PRDX4, CCT7, DHCR24, CHCHD6, DHODH, NUMBL, BCS1L, COA7, UBQLN4, FSCN1, CHCHD4, PDE4A, RCN1, CHID1, ATAD3A, PABPC4, EIF5A, TUBG1, SLC19A1, DCPS, PARP2, AIMP2, WDR18, SUV39H1, TARBP2, MRPL13, HNRNPAB, CCNB2, ATF3, MRRF, IRF5, ETF1, ENDOG, PSMG4, EIF5AL1, CREB3L4, PQBP1, GPT2, TYMS, MRPS12, CENPM, DCAF13, GRWD1, CKS2, CORO7-PAM16, DNAAF5, YIPF2, GPN2, CARM1, DDX21, KIF2C, BCKDK, NOL6, NTPCR, PVR, TTC26, HDAC7, KLF7, TMEM241, CDK4, KIF2A, ZNF232, NFKBIA, SPHK1, GINS4, MRPS15, PITRM1, ARL3, ELP2, RBFA, DPAGT1, RHOB, LONP1, FARSA, TIPIN, RUVBL2, INSL3, EIF4EBP1, GTF2H2C_2, LDHB, WDR77, TK1, CHAC2, PPME1, TOE1, GNL2, RAD51C, HNRNPA1, FAM72C, MTHFD1L, NDC1, RAD50, SNRPD1, AGAP3, SAAL1, IGF1R, SOGA3, MASTL, ZWINT, RRS1, ARID5B, TXLNG, AVEN, ERCC1, POC1A, DTYMK, MBD3, PRDX1, SARM1, GLRX2, SEMA4A, PGAM4, EXOSC7, CENPV, LRRC61, TMA16, LRRC75A, POLR1C, PABPC1L, AGO2, SYNGR2, CCNB1IP1, SLC27A5, GTPBP3, PKP4, UHRF1BP1L, KIF4A, FAM89A, NUDT8, TTF2, EZH2, FAM72D, CKS1B, NCDN, PTPRK, RCL1, ZNF282, GNA15, IPO5, WRAP53, ATRIP, HEATR5A, PDSS1, XRCC3, PHB2, CHKA, CYC1, NUP35, CDC25C, UBE2S, NEK2* |

**Table S3. List of differentially expressed genes in memory CD8⁺ T cells clustered by upregulation patterns following stimulation with IL-15, ionomycin or the combination of both.**

| | |
|---|---|
| **Cluster 1** | *ERRFI1, MINOS1, ZSCAN20, MRPS15, ZFP69, FUBP1, DNAJB4, CIART, ENSA, RORC, ARHGEF2, CD84, GPR161, CENPL, DARS2, SUSD4, GALNT2, NTPCR, ADSS, FAM107B, APBB1IP, C10orf35, PPRC1, NFKB2, ADD3, MXI1, SFXN4, BNIP3, UBQLNL, PLEKHA7, TIMM10, CD6, VPS37C, PRCP, MTMR2, BIRC3, FXYD6, ARHGEF12, ZNF202, RPUSD4, GAPDH, TPI1, KLRC1, CCDC91, CPM, FAM216A, FITM1, NID2, TMEM30B, ESR2, HSPA2, TTC9, EFCAB11, TNFAIP2, CEP170B, SERINC4, HDC, PIF1, CLN6, KIF23, CIB2, SNN, TXNDC11, ARL6IP1, MYLPF, ZNF747, FBXL8, HSF4, CENPN, GCSH, COTL1, NLGN2, MTRNR2L1, ANKRD13B, CPD, CCT6B, CDC6, STAT5B, HOXB2, CUEDC1, MRC2, PRR29, PRKCA, BAIAP2, DSEL, NFATC1, ADAT3, TRIP10, TMEM205, ZNF20, ILVBL, FXYD1, NFKBID, LYPD3, QPCTL, DHX34, PLEKHA4, ZNF432, ZNF841, MBOAT7, LAIR1, KLF11, CYS1, POMC, DNMT3A, RASGRP3, FAM98A, COMMD1, RMND5A, GPAT2, TBC1D8, TMEM182, CERS6, CERKL, CD28, FASTKD2, ISM1, CHMP4B, AHCY, SOGA1, MRAP, EVA1C, RCAN1, DDTL, DDT, NEFH, MTFP1, SETMAR, ITPR1, ARL8B, XYLB, KIF9, PFKFB4, SMIM4, ITIH4, IL17RB, LNP1, TRMT10C, ADCY5, PLXNA1, ACKR4, WWTR1, BCL6, CD38, SRD5A3, AREG, FAM175A, EIF4E, PAPSS1, HADH, C4orf32, PRDM5, TRIP13, MTX3, TMEM167A, FAM53C, REEP2, ZNF300, THG1L, RNF145, BTN2A2, HLA-DQA1, SAYSD1, DST, FAM135A, FYN, PLN, TPD52L1, ADAT2, PEX3, SERAC1, ETV1, PSMA2, LANCL2, PEG10, SLC25A13, TRIP6, UBE2H, ASIC3, DLC1, BMP1, RIPK2, TP53INP1, GRHL2, RFK, COL27A1, NR6A1, C9orf16, GPSM1, DYNLT3, CDK16, EFNB1, MID2, NSDHL, DDX3Y, CD24* |
| **Cluster 2** | *TNFRSF4, RCC1, YRDC, CCDC24, SLC6A9, GPX7, LMO4, ALG14, PRMT6, GPR89B, JTB, AIM2, CAMSAP2, IL10, TFB2M, PFKP, PITRM1, NET1, ANKRD16, MAP3K8, TIMM23, ADO, DDX21, NUTM2A, NOLC1, INPP5F, KCNJ11, ACP2, SLC43A3, CLP1, TAF6L, NDUFC2-KCTD14, ZBTB16, DPAGT1, STT3A, FAM118B, PTPN6, PLEKHG7, PLXNC1, PARPBP, BTBD11, MLEC, TMED2, MTUS2, NUFIP1, SLAIN1, DNAJC3, DTD2, FRMD6, FERMT2, AHSA1, TDP1, TMEM251, NIPA2, CHSY1, FAM173A, NETO2, CSNK2A2, MAF, ZCCHC14, CLUH, PSMB6, UBB, MRM1, STAT3, VPS25, G6PC3, HOXB7, ATP5G1, SLC35B1, PCTP, MRPS23, HSF5, P4HB, HRH4, TIMM13, C19orf70, RFX2, ZNF699, S1PR2, ZGLP1, ABHD8, PDCD2L, TMEM147, SPINT2, CTU1, LDAH, WDR43, LBH, CRIM1, IL1R1, IL1RN, TMEM177, GORASP2, HSPE1-MOB4, TMEM237, FZD5, TTLL4, EBF4, HM13, NNAT, CD40, SULF2, STAU1, SPATA2, CXADR, TIAM1, FAM207A, RANBP1, DERL3, SMTN, FOXRED2, TOMM22, NDUFAF3, IMPDH2, ARF4, SEC61A1, PCCB, TSC22D2, P2RY14, ATP13A3, JAKMIP1, TRMT44, SPINK2, CXCL10, NUDT6, LARP1B, DCTD, SUB1, NDUFAF2, SLC30A5, IL5, UQCRQ, IL9, ETF1, HSPA9, SPDL1, ZNF879, GCNT2, NOL7, GMNN, APOM, NOTCH4, PFDN6, BAG2, KCNQ5, SH3BGRL2, UBE2J1, MDN1, GPR63, PRDM1, ATG5, BEND3, CYCS, C7orf25, CYP51A1, MET, TPK1, ABCB8, RAB11FIP1, LYPLA1, NECAB1, POLR2K, SPAG1, NCALD, PTPRD, PIGO, MSMP, DNAJC25, UGCG, SLC27A4, SLC2A6, SH3KBP1, CHST7, SLC9A7, TIMM17B, HSD17B10, ITM2A, GLA* |
| **Cluster 3** | *UTS2, PARK7, ZBTB17, NBL1, STMN1, ZBTB8OS, GNL2, ZNF684, CTPS1, USP24, GADD45A, HHLA3, HS2ST1, GBP2, LRRC8B, LRRC8D, EVI5, EXTL2, VAV3, TTF2, POLR3C, CD160, GPR89A, HIST2H2AA3, HIST2H2AA4, OTUD7B, PMF1, FCRL3, SLAMF1, SH2D1B, CREG1, XCL1, F5, FASLG, GLUL, PIK3C2B, RASSF5, ATF3, JMJD4, FAM89A, SIPA1L2, GNG4, LYST, EDARADD, GTPBP4, PLXDC2, ZNF438, CREM, CCDC6, ARID5B, EGR2, SRGN, SGPL1, VCL, GLUD1, FAM35A, IFIT3, PCGF5, HECTD2, ALDH18A1, RRP12, PGAM1, ZDHHC16, UBTD1, GOT1, NPM3, PRDX3, RGS10, GLRX3, FUOM, PSMD13, TRIM5, TRIM22, MRPL17, NUCB2, NCR3LG1, PRRG4, CAT, CD82, PACSIN3, DDB2, FKBP2, ZDHHC24, CTTN, FCHSD2, MAML2, SLC35F2, DIXDC1, REXO2, SIK3, CD3D, PHLDB1, CRTAM, FKBP4, CD27, LAG3, KLRD1, MAGOHB, YBX3, AEBP2, KIF21A, TMEM117, AMIGO2, VDR, SMAGP, NR4A1, ATG101, TESPA1, IFNG, MDM2, PHLDA1, UHRF1BP1L, EID3, CRY1, ATP2A2, PTPN11, POP5, P2RX7, P2RX4, ORAI1, CDK2AP1, RILPL2, PUS1, USP12, HSPH1, RGCC, NEK3, TBC1D4, GPR18, PRMT5, SAMD4A, PRKCH, HIF1A, AKAP5, ZFP36L1, DCAF4, PTGR2, ZNF410, NPC2, TMED10, RIN3, ATG2B, TRAF3, CKB, EMC4, SLC12A6, NOP10, BMF, EHD4, PDIA3, ATP8B4, SLC27A2, GABPB1, MAPK6, C2CD4A, C2CD4B, SNX33, TSPAN3, PEAK1, NMB, TBC1D24, PRSS21, HCFC1R1, MTRNR2L4, C16orf45, ABCC1, ITPRIPL2, ZNF768, BCKDK, CCL22, KIFC3, PLEKHG4, NFAT5, PHLPP2, NUDT7, GAN, KIAA0513, KDM6B, TMEM88, FAM106A, C17orf51, CORO6, EVI2B, EVI2A, COPRS, CCL4, CCL4L1, CCL4L2, DUSP14, MRPL45, PNPO, HOXB5, HOXB6, MSI2, ICAM2, MAFG, CD7, SECTM1, ENOSF1, TGIF1, SEH1L, RBBP8, MAPRE2, DOK6, CD226, SLC39A3, TNFSF9, ANGPTL6, SYDE1, ZBTB32, IGFLR1, SERTAD1, LTBP4, ERF, TOMM40, RELB, IL4I1, NAT14, RHOB, HNRNPLL, GALM, FOXN2, PELI1, POLE4, EIF2AK3, IL18RAP, SLC20A1, GTDC1, NR4A2, GPD2, UBE2E3, ITGAV, NAB1, HSPD1, HSPE1, PLCL1, ICOS, IKZF2, CHPF, SERPINE2, SP140, HTR2B, UBE2F, CPXM1, ATRN, SMOX, PRNP, MAP1LC3A, SPAG4, ROMO1, PKIG, ADA, PABPC1L, TOMM34, PIGT, ZNFX1, PTPN1, TSHZ2, GNAS, HSPA13, BTG3, SOD1, URB1, ZBTB21, C21orf2, SLC35E4, DUSP18, TNFRSF13C, CENPM, PRR5, GRAMD4, TATDN2, CSRNP1, CMSS1, FILIP1L, ALCAM, CBLB, ZBED2, CD200, BTLA, TIGIT, B4GALT4, TIMMDC1, CD86, PDIA5, RPN1, SRPRB, ATP1B3, CHST2, SIAH2, GOLIM4, MAP6D1, PARL, PPP1R2, RHOH, TXK, PPAT, CXCL9, RASGEF1B, HPSE, PKD2, GPRIN3, OSTC, CAMK2D, SEC24D, FGF2, SPRY1, INPP4B, SRD5A1, ANKRD33B, ZNF622, PTGER4, TTC33, OXCT1, HMGCS1, ADAMTS6, MAST4, LHFPL2, HOMER1, PAM, WDR36, STARD4, IL13, IL4, CTNNA1, LRRTM2, SIL1, NDFIP1, EBF1, FBXW11, SFXN1, B4GALT7, SERPINB9, RPP40, TXNDC5, TMEM170B, GFOD1, RANBP9, JARID2, UBD, GTF2H4, PPT2, HLA-DMA, FOXP4, POLR1C, MTRNR2L9, CD109, NT5E, BACH2, NUS1, MAN1A1, NCOA7, MYB, AHI1, LTV1, PLAGL1, EPM2A, FBXO30, TAB2, OPRM1, IPCEF1, TIAM2, SNX9, TAGAP, IGF2R, SLC22A1, FSCN1, RPA3, GLCCI1, BZW2, MACC1, CDCA7L, OSBPL3, NFE2L3, HNRNPA2B1, ELMO1, RHBDD2, NPTX2, NAMPT, CCDC71L, DNAJB9, CAV1, FAM3C, SND1, LRRC4, WDR91, CREB3L2, C7orf55-LUC7L2, NDUFB2, MTRNR2L6, ATP6V1B2, AP3M2, SDCBP, ASPH, RDH10, TMEM70, PKIA, ZNF704, FABP5, TMEM64, RNF19A, TRIB1, ASAP1, SLA, CHRAC1, AGO2, BNC2, TESK1,* |

| | |
|---|---|
| | CD72, KLF9, GCNT1, PRUNE2, CKS2, GADD45G, NFIL3, TMOD1, ANKS6, NR4A3, CTNNAL1, BSPRY, PSMD5, MRRF, SLC2A8, ZNF79, PRRX2, NCS1, PPP1R26, GK, MAGEH1, OGT, PGAM4, BEX5, PAK3, UBE2A, GLUD2, ZDHHC9, PHF6, FHL1, MAP7D3, MAMLD1, HMGB3 |
| **Cluster 4** | PERM1, ACOT7, ENO1, FBXO6, MAD2L2, CLIC4, AZIN2, LRP8, HOOK1, DNAJC6, PDE4B, CRYZ, GBP1, CD53, C1orf162, PHGDH, MLLT11, S100A11, DUSP23, MPZL1, KIAA0040, LIN9, ITIH2, MCM10, VIM, SVIL, C10orf128, ZWINT, CISD1, COMTD1, CEP55, PDLIM1, GOLGA7B, IFITM1, IFITM3, PPFIBP2, RRAS2, SLC35C1, FAM111B, FADS2, PPP1R14B, CFL1, CCDC85B, DPP3, CORO1B, IL18BP, PAK1, HYLS1, GLB1L3, CCND2, CD9, SLC2A14, MRPS35, CCDC65, TUBA1B, TUBA1C, NDUFA12, PITPNM2, DNAJC15, CLN5, RAP2A, MRPL52, PSME2, GZMB, PSMA6, WDHD1, GPHN, CATSPERB, ASB2, CINP, INF2, JAG2, AVEN, SPRED1, C15orf48, ANXA2, PTPN9, CTSH, MTHFS, ST20-MTHFS, TM6SF1, RCCD1, TNFRSF12A, MMP25, IL32, CORO7, CIITA, CPPED1, MKL2, IL4R, CORO1A, MT1E, CPNE2, CBFB, NQO1, AARS, ATP2C2, CDT1, RILP, P2RX5, PFN1, C17orf49, EIF5A, TNFSF12, TNFSF13, SOX15, TP53, COX10, PLD6, MAPK7, MFAP4, CISD3, RARA, IGFBP4, TNS4, PSMC3IP, MMD, PTRH2, INTS2, TACO1, RGS9, SLC16A6, RNF157, UBALD2, TK1, NPTX1, ACTG1, ARHGDIA, YES1, EMILIN2, MYL12A, ACAA2, MYO5B, GZMM, IZUMO4, ZNF77, GNA15, SH2D3A, KANK3, ICAM1, RAVER1, PDE4A, LDLR, TPM4, IFI30, PDCD5, GPI, HCST, KCNK6, IFNL1, CEACAM1, PLAUR, PPP5C, FLT3LG, CLDND2, KIR2DL3, KIR2DL4, RNF144A, IAH1, RRM2, TRIB2, TTC27, SOS1, EPAS1, SPRED2, BOLA3, TMSB10, CAPG, POTEF, CCDC74A, SLC4A10, WDR12, CXCR1, BCS1L, STK16, TUBA4A, FARSB, PCNA, GINS1, BCL2L1, TLDC2, SAMHD1, MYBL2, UBE2C, TMEM189, C20orf197, SLCO4A1, RGS19, ADAMTS1, CHAF1B, PSMG1, WDR4, SLC19A1, OSM, APOL6, LGALS1, APOBEC3G, DESI1, PARVB, GTSE1, DENND6B, ADM2, SCO2, TYMP, BHLHE40, KAT2B, CCR1, CCR2, CCR5, HYAL3, HYAL2, C3orf18, MAPKAPK3, DUSP7, TWF2, ARHGAP31, FNDC3B, PGM2, PTTG2, SCFD2, SLC39A8, CISD2, ELOVL6, CASP3, AHRR, TARS, GZMA, COMMD10, EGR1, HBEGF, PDGFRB, FAM196B, LCP2, HRH2, PRELID1, PDLIM7, EEF1E1, NEDD9, TUBB, LTA, LTB, LST1, NCR3, DDAH2, CLIC1, HSPA1A, KIFC1, ETV7, PIM1, CCDC167, TREML2, CCND3, DNPH1, NFKBIE, RCAN2, MANEA, ARHGAP18, AIG1, STX11, SLC22A3, LFNG, TTYH3, ACTB, GARS, UPP1, STX1A, SLC25A40, ASNS, BPGM, NUP205, TBXAS1, FAM131B, MFHAS1, LZTS1, DOK2, UBXN8, SLC20A2, MYBL1, MSC, MRPS28, TPD52, NDUFB6, KIAA1161, PSAT1, IARS, ZNF367, DNAJC25-GNG10, SUSD1, SNX30, TNC, NEK6, ARPC5L, ZBTB34, FAM129B, SDCCAG3, NOTCH1, NPDC1, SAPCD2, UAP1L1, TUBB4B, FAM166A, TOR4A, NSMF, PRKX, EBP, RBM3, PIM2, PDZD4 |
| **Cluster 5** | CITED4, CD69, BCAT1, SNRNP25, IMPA2, TNFSF14, QTRT1, ETV2, TIMM50, KIR2DL1, KIR3DL1, KIR2DS4, PNO1, CKAP2L, NANP, MIF, RRP9, NLN, PTTG1, RPL26L1, CEP57L1, MICAL1, SQLE, NOL6, NOXA1 |
| **Cluster 6** | SRM, PIGV, KIAA1522, CKS1B, SEMA4A, GPATCH4, UCK2, IVNS1ABP, SFMBT2, RTKN2, PPA1, LRRC20, DUSP5, STIP1, BANF1, EMP1, DUSP6, PMCH, SCARB1, CHAC1, SNX22, HAPLN3, CCDC78, THOC6, SOCS1, GPT2, GINS3, CES4A, PIGW, AARSD1, SOCS3, RNF165, DUS3L, CD320, ANKLE1, KPTN, HSPBP1, CAD, SMYD5, DPP4, ATIC, TOX2, SLC17A9, XBP1, LIF, CRELD2, CISH, FRMD4B, LPP, ACSL1, CCT5, GNPDA1, PPARGC1B, ADAM19, CDKN1A, SNX8, AHR, CDK6, ERI1, EXOSC4, BOP1, ALDH1B1, GNG10, TNFSF8, MBTPS2, KLHL15, ELK1, C1GALT1C1 |
| **Cluster 7** | ATAD3A, TNFRSF9, SLC25A33, AGMAT, MRTO4, ZNF593, DNALI1, DPH2, B4GALT2, MMACHC, PRDX1, BTF3L4, AK4, CSF1, SLC16A1, XCL2, BATF3, SCCPDH, IDI2, IL2RA, SCD, CTSC, HYOU1, HSPA8, FEZ1, PCED1B, ACVR1B, METTL1, UBE2N, SOCS2, GNPTAB, HSP90B1, SLC7A1, ALG5, DGKH, NUDT15, NDFIP2, PTGER2, WARS, TRMT61A, RAD51, PPIB, PKM, SEMA7A, RPP25, PIGQ, BOLA2B, BOLA2, DCTPP1, IRF8, SLC7A5, HIC1, SPNS2, MYBBP1A, PFAS, PEMT, CCL3, PSMB3, PTGES3L-AARSD1, NME1, AXIN2, MRPL12, PYCR1, TYMS, TAF4B, SEC11C, BCL2, RBFA, MYDGF, MRPL4, RNASEH2A, CALR, NR2C2AP, EXOSC5, CD3EAP, SLC1A5, ATP6V1C2, PDIA6, ADCY3, MTHFD2, HK2, MAL, IMP4, FMNL2, ACVR1, PGAP1, GMPPA, CCL20, PER2, NXT1, CECR5, SDF2L1, ZNRF3, CMTM6, LARS2, MANF, MRPL3, ECE2, DNAJB11, MB21D2, NRROS, WFS1, AFAP1, PAICS, HOPX, SLC9B2, ARSB, HAVCR2, NUDCD2, NOP16, RMND5B, NHP2, IRF4, LYRM4, BMP6, IER3, TNF, VARS, WDR46, KCNK5, BYSL, SLC29A1, HSP90AB1, RPF2, SGK1, MTHFD1L, KDELR2, DNAH11, SNX10, SEC61G, PUS7, MDFIC, PDIA4, SLC39A14, POLB, RRS1, LAPTM4B, NUDCD1, SNTB1, MYC, RCL1, FANCC, SEC61B, TXN, HSPA5, VAV2, PRDX4, TIMM8A, OCRL, CD40LG, MPP1 |

**Table S4. List of differentially upregulated genes in memory CD8$^+$ T cells induced by IL-15 or IL-15 + ionomycin stimulation.**

| | |
|---|---|
| **Upregulated DEGs by IL-15** | ITGB1BP1, IFFO2, SH2D2A, PLP2, ITGA5, GNAI2, ITGA6, CTU2, DAXX, CISD3, MID1IP1, MGAT4A, CARS, KIAA0754, ST3GAL1, ITPR3, ZBTB2, TIAF1, DHRSX, MTHFD2, CCR7, NFATC2, ZNF641, ISG15, PTPRE, YES1, DHCR7, PPM1M, TSPAN14, PTK2B, TWF2, PLEKHG2, TAPBP, MYO5B, TTC7A, SLC25A25, ST20, ACTR3, ARPC1B, CROCC, RNF125, FAM89B, CIDEB, EPB41, PLA2G16, FGFBP2, CACNB3, MLLT6, FLI1, ZNF789, ICAM3, TUBA1C, SLC35B4, GAB3, ZNF319, FNDC3B, FBXO44, LOC400927-CSNK1E, PPP1R14B, IFITM3, SLC2A1, UBTF, MR1, FAM84B, NDUFS4, PPARD, DBP, TSC22D4, CAP1, TOR1AIP1, PLOD3, ZBTB7A, OASL, TRIM21, UNC13D, PECAM1, CPPED1, C14orf159, CD244, CKLF, PROK2, ISG20, MX2, TTC13, OPTN, TMEM200B, P2RY10, TMIGD2, PARP10, FAM63B, CRIP1, RPL39L, NLRP1, GCH1, MRPL2, MVP, PBXIP1, APOBEC3G, BTG2, SAMD3, LIME1, IFI35, NFIA, NOTCH2NL, RBCK1, GRIP2, ZC3HAV1, ATP10A, KIR2DL3, TCIRG1, CKLF-CMTM1, RASGRP1, SLCO3A1, 45901, RUNX3, PDSS1, DHRS1, TYROBP, ZNF652, CD101, TTC32, F2RL2, TMEM39B, TET3, GZF1, TRABD2A, KNSTRN, SLC25A4, ASNS, S100A6, MKNK2, LIX1L, CLN3, ZBTB45, ARHGEF9, CASP8, PLEKHF1, C3AR1, TRERF1, CCHCR1, DENND1C, F11R, FAM166A, S1PR1, FAM105A, SCFD2, POC1B, FERMT3, PPIF, STK40, IRF9, ARRB2, MT2A, RSU1, PIK3AP1, EPHB6, PDP2, GRK6, RELT, LASP1, CRYZ, OXNAD1, GMIP, TUBB4B, MAP7D1, IFIT2, RAP2A, BCL9L, TRIB2, SYNE1, UBA7, SSH2, ST3GAL5, TGFBR1, TMSB4X, GALNT10, AES, CYFIP2, CCR6, FADS1, TMEM104, PLCB2, INPP1, INTS2, IFIH1, USP28, PTPN4, TNFRSF25, GSDMD, GYPC, FCHO1, LPXN, CCDC112, PSMB10, CRYL1, ARHGEF19, MYBL1, ZNF101, RBM38, KATNAL1, FLOT1, OAS2, HSPA1A, PITPNC1, CRB3, TRANK1, EVL, ACAA2, CCDC88C, NQO2, CSNK1E, NSG1, ABR, IER5, PLAUR, CAMK2G, FAM213B, APOBEC3H, GMFG, NLGN3, SLC39A10, TNFRSF1A, GBP1, C1RL, PNP, DUSP5, MBP, EML4, GIMAP5, ELP4, MANEA, ACSL5, CASP6, DDX60, MFHAS1, PRKD2, IFITM2, NR1D1, PIK3CG, DHCR24, SLC25A42, BBC3, CFL1, ZAP70, SP4, SLC2A14, COMMD7, TMEM63A, FGD3, ZNF33B, IL4R, CCM2, PIEZO1, SLCO4C1, LDLR, TREX1, FAM65B, LDLRAP1, CASP1, BIN2, ZNF80, CAPN2, IER2, C9orf142, PRKCB, CCND2, ABCB1, IRF1, TUBA1B, PPP3CA, ANKRD13D, TMEM14A, FAM46C, GRAP, CMTM3, BCL11B, SYNGR1, HN1, IFNLR1, EEF2K, SH3BP5, NLRC5, SLC25A20, MYH9, TUBB, TCF7, RCSD1, EGR1, ZFP36L2, TSNARE1, CXCR5, THRA, ARMT1, ZNF792, GYG1, MPZL1, TMEM19, GM2A, DGKA, ZYX, PAK1, STX11, RASSF3, SMYD2, TMOD2, ARHGAP30, AGAP2, SLC9B1, SLC18B1, VANGL1, CYB561, ACOT1, TNFRSF14, CDK5R1, BCAT1, CDC42SE1, IL11RA, SAMD9L, INTS7, MX1, CNPY4, TRDMT1, VAV1, OXR1, FANCA, ALDOC, LCP2, CLDND2, GNB5, ST3GAL4, STX3, NEDD9, HIP1, HS3ST3B1, HPGD, LSS, LAPTM5, PTPN9, ADAM19, CLIC1, AMPD3, ANKS1B, TACC3, HYAL2, GPSM3, LST1, RNF44, CDC25B, KIAA1841, CLYBL, CA5B, GSTK1, APOL1, RNF165, ZNF449, CD53, DDX43, TRIM14, KIF3C, FLOT2, SKI, MATK, VASP, MRPS28, SEMA4B, ARHGAP11A, FHOD1, RFTN1, OTUD3, TK2, OCIAD2, AKNA, UBALD2, RCBTB2, LMNA, MAP4K1, S100A11, ARHGEF1, SLC22A5, MAPKAPK3, ADCY9, XAF1, SLC27A3, LCP1, ERV3-1, MICAL1, LDLRAD4, AKAP2, TTC38, RALGDS, TPM4, RASA3, BHLHE40, NOL4L, ARHGEF26, ZNF276, AGPAT2, TMEM120B, ENPP4, TUBB4A, STK16, GBP3, EHD1, DOCK10, SLFN5, DDAH2, SLC1A4, TUBA1A, RTP4, CHAC1, ATXN7L1, SQRDL, IDH2, GOLGA8R, FAM102A, PLXND1, SLC26A11, TMCC3, C16orf54, SH2D3A, SSBP4, PLEKHO2, C5orf58, LCK, ISYNA1, SH3BGRL3, PPP1R18, TAGLN2, LAT2, FAM231D, GPD1L, RASSF1, ABI3, C4orf33, ITM2C, GOLGA8N, RASAL3, C1orf56, FAM8A1, DENND1B, AP1S3, PIK3CD, EFHD2, ZNF600, ASCL2, FAM78A, IMPDH1, SLC2A4RG, GOLGA8O, MSC, DEF6, CTSH, IFI30, EHBP1L1, MFNG, HDAC4, RRM2, IL10RA, CYP4V2, ASB2, CLUAP1, LPAR5, RGS19, BICD1, CECR1, CALHM2, ENPP5, CORO1B, RNASEL, CCL5, TOR4A, DCK, NPRL2, POC1B-GALNT4, SPOCK2, GPR55, GALNT4, CSK, PATL2, UPP1, DOK2, TCF19, ANXA2R, KIAA0040, IL2RB, PDZD4, CARD6, MBOAT1, TBC1D10A, IRF2BPL, PARVG, TBKBP1, SIGIRR, SLC25A43, PRDM11, CARD16, ACOT7, TXNIP, SLC25A53, PLCG1, PIK3R5, TNFSF13, ARHGAP4, SLC9A9, CORO7, TGFB1, VIM, PLEKHN1, TUBA4A, AHNAK, TFEB, GIMAP7, ABCA7, NOTCH1, TRIB3, CMIP, CAPG, STK10, UAP1L1, ANO9, RHOF, CEBPD, IQSEC1, EPHX1, PTP4A3, CUBN, LRFN1, LPAR2, SLC12A7, EFCAB7, CCDC30, CCNI2, ADCY7, TCTEX1D2, TC2N, TUBB2A, GSAP, CTNND1, SPTB, SOCS1, TRAF3IP3, CYP2R1, ZNF710, KIR3DL1, CD5, GALNT3, TNFSF12, GIMAP6, BCL2L1, E2F3, DDIT4, EIF4EBP1, LSP1, SIPA1, APOL6, CYTH4, TMSB10, HEMK1, EBP, PDE4D, UBASH3A, EPHA1, OAS1, ACAP1, PSD4, ACTG1, KLF13, TNFRSF12A, IFITM1, SH3BP2, PROCA1, LGALS9, PYCARD, RARA, RHBDF2, MYL12A, KIR2DL4, B3GAT1, C1orf21, KCNAB2, FAIM, GPR132, SLC16A6, SLC14A1, HSPA1L, SLC9A3R1, SMAD9, PFN1, DUSP7, PSAT1, MOB3A, ZFYVE28, STMN3, VNN2, SLC7A11, GOLGA8J, MAD1L1, GRAP2, LYZ, SYNM, PXN, ATP2A3, APOL3, RNF43, TLDC2, NEK6, MCM10, RNF213, ARHGDIB, TMEM229B, GOLGA8K, RARRES3, LGALS3BP, TK1, DRAM1, ACTB, C1orf74, TSC22D3, CCL27, PTTG2, MSRA, PRR5L, LTK, TMEM156, USP18, CCDC74B, ADGRG1, PECR, OSBPL5, ADGRG5, PLAC8, ACACB, KDELC2, KLF10, FAM179A, RNF166, LAT, PIM2, SAMD10, SH3BP1, MB21D1, KIF21B, NQO1, PAQR8, SNX30, PRMT7, EMILIN2, BPGM, RHEBL1, ITPRIPL1, HIVEP3, MYADM, ALOX5AP, SPN, 45909, RGMB, TPM1, BHLHB9, NBEAL2, SLCO4A1, TSEN54, LRP8, SASH3, CNN2, HCST, ADGRE5, C5orf63, DUSP6, HSD11B1L, ICAM1, LIPT1, TMEM204, TMEM55A, SAPCD2, PREX1, TMEM71, TTYH2, APBA2, CTSW, FLT3LG, S1PR4, MIDN, CORO2A, SNX21, SH2D3C, TOR2A, OAS3, LIN9, DHRS3, RNF144A, CX3CR1, IL16, PLCD1, C11orf21, FBXL16, LYSMD1, DDI2, SELPLG, ITGAM, BIN1, RAC2, KANK3, ZBTB46, PDE4A, PWWP2B, AHRR, GIMAP4, NCR3, RASSF2, SBK1, GTSF1, ARAP3, POU5F1, SUN2, EMP3, CCDC74A, FMNL1, TSPAN32, PDLIM1, LGALS1, FBXO6, GNGT2, NLRC3, OGFRL1, FAM26F, AQP3, CYBA, AGTRAP, DUSP10, KYNU, CIITA, P2RY8, TJP2, SAMHD1, FOXM1, TRPV2, GIMAP8, LTA, OTOF, DBN1, PTGDS, SEMA4C, SLC4A8, OSM, BCL7A, IKBKE, CPNE8, TMOD4, SEMA4G, SERF1A, PLK4, CCR5, ATL1, S100A4, RNF157, SORL1, TRPA1, IL32, PPM1N, PIM1, GZMM, CLU, FLT4, NHSL1, RGS14, ORC6, PHGDH, RAB33A, ATP2C2, TMC6, BEST1, PTPRCAP, NUDT14, FAM167B, TBXAS1, GZMH, PAOX, CMTM4, DAB2, |

| | |
|---|---|
| | *RGS16, CXCR6, DOCK5, NCF1, WNT1, SIRPG, FAM171A1, EPHA4, CACNA2D2, PROSER3, CAPS2, IRAK2, IL6R, MAN1C1, CLIC5, CD52, ITGB2, RGL4, POLN, NMUR1, MYO1F, ZWINT, BCL3, TMEM154, PLEKHG3, NKD1, TNFAIP8L2, CDKL1, FCGR3A, MYO1G, CMPK2, GZMK, CSGALNACT1, ICOSLG, GLIPR2, PCDH1, FAM131B, TIMP1, ABCA2, FAM129B, CEACAM1, TTC16, AKR1C3, CRIP2, SUSD1, FXYD2, PRR7, SYTL1, CXCR3, FXYD6-FXYD2, ITGAL, LTB, S1PR5, TBC1D10C, RAB37, CORO1A, GDPD5, ELOVL6, PDLIM7, RGS9, PLEC, LGR6, NLRP3, LXN, IZUMO4, E2F2, GALNT6, PYROXD2, SPRED2, FLNA, AOAH, N4BP3, ADAMTSL5, DNAJC6, FBLN5, KLF8, RASGRP2, LLGL2, KCNN4, RGS18, ADAM8, DNAH7, KBTBD11, OR2A1, HEATR9, SH3TC1, KIF19, APOBR, H2AFY2, ZNF257, ANKEF1, POU5F1B, PDGFRB, MMP25, GPR68, CISH, DNAH14, BUB1, RTN4R, CITED4, TMEM200A, PVRIG, SGCD, ITGB7, LRRC63, PDE6C, TMC8, GLB1L3, KLF2, CDKN2D, MAP2K6, MEGF6, FCER1G, C3orf18, FCRL6, JAG2, PERM1, MLLT11, RAP1GAP2, C1orf162, CCDC65, ERCC6L, PDGFB, CACNA1I, MTFR2, FCGR3B, UBXN11, ATP1A3, ETV7, IFNL1, SERPINI1, DTX4, LFNG, CD79B, CDT1, DPEP2, FGR, CD300A, MUC12, GIPC3, FCRLB, CCR1, SPNS3, SLAMF8, ADM2, SPON2, TNC, HBEGF, CPNE2, GZMA, CDK20, CMKLR1, DAPK2, FAM196B, CXCR1, PDLIM2, PLXNA4, KIAA1671, CXCR2, LZTS1, TREML2, RCAN2, ADAMTS10, NOXA1, FRMPD3, P2RX5, HPCAL4, RAB27B, RDM1, ZNF365, CCR2, CATSPERB, CLIC3, ERP27, IGFBP4, TNS4, ARRB1, SUSD3, IGFBP3, C20orf197, NHSL2, ZNF683, HLA-DMB, SNAI3, SLC22A3, GOLGA7B* |
| **Upregulated DEGs by IL-15+ionomycin** | *FILIP1L, CD200, F5, C2CD4A, C2CD4B, PRRX2, EGR2, TG, CTTN, PACSIN3, CXCL9, NPTX2, CD160, HECTD2, CD109, MAP6D1, CPXM1, SAMD4A, SERPINE2, IL9, SPAG4, SMOX, HOXB5, NR4A2, NOTCH4, HOXB6, SH2D1B, IL31, IL1RN, PKIG, COL6A3, BNC2, RASGEF1B, IFNG, BSPRY, TMEM88, LRRC4, ZBED2, PRH1, KCNJ11, KANK2, PAK3, SLC27A2, IL4, TNFSF9, CCL4L1, CCL4L2, MET, ADAMTS6, UBTD1, REEP2, TRPM8, SIPA1L2, HSF5, NR4A3, TRPC1, NR4A1, ZNF704, HSPA2, TNFRSF9, NCR3LG1, GPR34, EFNB2, SLC22A1, C11orf96, SPINK2, NMB, SPRY1, UBD, EBF1, VCAM1, GPR63, KCNS1, C16orf46, XCL2, KLF5, CEP72, XCL1, CTNNAL1, CRTAM, ATF3, PRDM5, NID2, ITGB8, TUBA8, LCA5L, FSTL3, CENPM, NNAT, AREG, SLA, GPR19, CAV1, RBFOX2, CAMK2N2, FGF2, GFOD1, SULF2, PRRG4, FABP5, MAML2, IL5, CCL3, CCL4, MAP1LC3A, EVI5, CHN1, MACC1, HPSE, SLC35E4, TMEM117, HRH4, TMEM86A, DST, IL10, FCRL3, PRUNE2, ZKSCAN2, C16orf86, DCHS1, CXCL10, MASP2, TP53INP2, MOK, PPFIA3, FAM167A, NFKBID, C16orf45, NBL1, JAG1, PLCL1, EXOC6B, TMEM170B, COL5A3, PEG10, PAM, CCL20, CCL22, RBBP8, CHST2, CD72, AKAP5, EBF4, BTLA, CCDC168, GADD45G, IL1R1, LYST, ADCY5, PLXDC2, CD40, CD82, EVI2A, EXTL2, GABPB1, TRIB1, STRIP2, KIAA0513, TSHZ2, IL23R, MTRNR2L4, FAM209B, FRMD6, SNX9, ADA, GBP2, SLC25A34, DIXDC1, PDIA5, NFE2L3, IL13, FBXO30, RND2, RDH10, TGIF1, LRRC8B, HOXB7, SECTM1, GPD2, GATB, ZNF768, CDC42BPB, GNG4, C17orf97, PRNP, HIST2H2AA4, TIAM2, PMEPA1, ZNF629, GALM, NEK3, HIST2H2AA3, TESK1, CTLA4, MAMLD1, ZFP36L1, KLF9, RILPL2, TRIM22, HNRNPLL, MYO3B, JARID2, ITFG2, AFAP1, NT5E, EVI2B, RGCC, VAV3, AIM2, PDGFA, BBS12, FASLG, GK, ADAM9, FHL1, CKB, MYLPF, OPRM1, AMIGO2, PLEKHG7, KLRC4, ALCAM, GRAMD4, LTBP4, IPCEF1, BTBD11, IGF2R, CD226, FDXACB1, AHI1, SRGN, ANKRD33B, ERRFI1, THBS1, OTUD7B, CHPF, MTRNR2L9, NET1, EHD4, BCL6, FAHD2B, MCF2L2, IGFLR1, LRRTM2, ITPRIPL2, GLCCI1, PPCDC, TRIM5, TAGAP, PDZK1, ZBTB16, TDP1, SIAH2, CD84, NUDT7, PKIA, HIST2H2BE, WDR91, ATP6V1B2, CAMSAP2, KCNRG, SIK3, CTNNA1, ARID5B, TAF6L, ZCCHC4, ZNF821, ANKRD9, HOPX, DUSP18, CAMK2D, POLR3C, TP53INP1, CREM, CDKN1C, PCGF5, MED12L, HSPH1, DNAJB5, NCS1, CBLB, PFN2, RASGRF2, TM7SF2, NUTM2B, PPT2, NAMPT, GLUD1, BEND3, FAM3C, ANKS6, BCL2L11, CD7, RNF19A, PLEKHA4, CDCA7L, IL4I1, UBE2E3, ZEB2, MAP7D3, CD27, FSCN1, LBH, FZD4, MRM1, RIN3, SYCE1L, IKZF2, MPP1, IRF4, PKD2, TBC1D4, KIF21A, PEX3, PLAG1, KDM6B, PAICS, YBX3, BRSK2, LAG3, TSPYL2, GCNT1, PHLDA1, ADAT2, KIFC3, KLRF1, USP24, PMCH, NFAT5, ZBTB21, IL2RA, MAPRE2, CREG1, N6AMT1, NCOA7, BMF, PEAK1, P2RY14, RGS2, MSI2, OXCT1, GAN, ERF, KBTBD8, ZCCHC14, TSPAN3, ITGAV, VSTM4, INPP4B, PGAM4, POMC, FBXW11, DNAJB4, PGAM1, MAP3K9, BACH2, EOMES, PLEKHG4, ASPH, HMGCS1, OSBPL3, FCHSD2, DDB2, TMEM64, NDFIP2, DDX3Y, MAST4, MPZL3, BACH1, BATF3, KLRC1, ATP9A, ATP1B3, TIGIT, IL18RAP, SPAG1, EFNA5, GLUD2, NUTM2A, OGT, TSC22D2, CRY1, SNX33, SLC12A6, TTC9, MYB, RGS10, ZNF438, PON2, SRD5A3, INPP5F, TMEM70, HOXB2, ANK1, SRD5A1, HEATR5A, ZBTB10, CDKN2C, STARD4, HOMER1, FYN, GNAS, TAS2R30, SERPINB9, CCDC6, MTRNR2L6, ARHGEF12, IFT57, FAM200A, UMAD1, FAM216A, TRMU, MBOAT7, PTGER4, DUSP2, SRGAP1, MAP3K8, CREB3L2, NDFIP1, UHRF1BP1L, PARL, CMSS1, ZC3H12D, GPR18, SPDL1, DUSP16, ATG2B, CCDC24, IRF8, ENOSF1, RNF115, GADD45A, ASAP1, YPEL2, NFIL3, POU6F1, ARL3, RANBP9, COQ2, SP140, ELL2, PYCRL, PPP4R2, AXIN2, TPK1, PAQR7, RHOB, UBE2F, PTPN22, SNTB2, PLGLB2, EBLN2, PAXBP1, MTRNR2L1, FOXD2, ABCC1, WARS2, RHBDD2, CCDC141, GFI1, BMP8A, URB1, BZW2, PLGLB1, ELMO1, RORC, STAT5B, SGPL1, RBBP9, TTC33, SLC2A13, ACTA2, SLAMF6, IGHMBP2, MBIP, TTL, MXI1, CDK2AP1, IL6ST, ADD3, ITPKB, ICOS, GLUL, ZFP91, ATP13A3, LMO4, NCALD, GPR89A, PPP1R2, SLC2A8, EED, MTRNR2L3, DYNLT3, MAGEH1, WWC3, HAVCR2, DTNB, GPR89B, DNMT3A, KLRC2, RELB, CSNK2A2, MAF, SPRYD4, AGO2, PHLPP1, PHF6, TOMM34, WBP4, TIMMDC1, KLRC3, CA11, TMEM251, ZNF350, ZNF358, KIAA1524, KIAA1324, FAM89A, FOXP1, RRNAD1, ZNFX1, SDCBP, NAT14, SCD5, NEU1, COPRS, LRRC8D, SNTB1, CCDC91, FOSL2, KLF12, MTSS1, WDR36, TP53BP1, SOD1, PIGT, SNAPC4, PTPRM, EMC4, HERPUD2, DMXL1, PIK3C2B, AP1AR, P2RX4, POLB, HDAC5, HLA-DMA, RAD54L2, CTPS1, ITM2A, RFX7, TNFAIP3, CEP126, DOK6, HSPA13, ZNF831, TMEM2, MTRNR2L10, MPP7, GALNT2, TMEM245, RGPD5, WARS, RCL1, TTF2, CD6, SLC9A7, PRKCA, MDM2, MTX3, FOXP4, NFATC1, SMAGP, PRKCH, GPR137B, LRRFIP2, CTBS, STMN1, RBMX, SPAG9, SDC4, SND1, PRDM1, BTG3, CAT, ALDH5A1, HELQ, PDE8A, CMTM6, PPP3CC, ZSWIM6, ADSS, DCAF17, TIMM17B, TENM1, TIMELESS, TESPA1, ISCU, NPC2, HERC5, PHACTR2, SCAPER, TTLL5, PHF13, UTS2, ERLEC1, BTN2A2, USP12, TBC1D24, PPAT, HGSNAT, BEX5, SRGAP2, GPR155, TRIM34, PAPD5, LAX1, C7orf55-LUC7L2, NR3C1, AMFR, GOT1, MSANTD4, GOLIM4, ZNF75D, NRROS, BTG1, CD40LG, LRRC37A, B4GALT4, THADA, ZBTB38, ATP2A2, PKM, STAT3, NIN, DLG1, SPATA1, ZNF410, APBB1IP, UBAC2, TNF, RAB11FIP1, FGL2, ACAD11, FBXO32, ZC3H8, BTRC, PRSS23* |
